# Chemical Reversible Crosslinking Enables Measurement of RNA 3D Distances and Alternative Conformations in Cells

**DOI:** 10.1101/2021.11.19.469208

**Authors:** Ryan Van Damme, Kongpan Li, Minjie Zhang, Jianhui Bai, Wilson H. Lee, Joseph D. Yesselman, Zhipeng Lu, Willem A. Velema

## Abstract

Three-dimensional (3D) structures dictate the functions of RNA molecules in a wide variety of biological processes. However, direct determination of RNA 3D structures in vivo is difficult due to their large sizes, conformational heterogeneity, and dynamics. Here we present a new method, Spatial 2’-Hydroxyl Acylation Reversible Crosslinking (SHARC), which uses chemical crosslinkers of defined lengths to measure distances between nucleotides in cellular RNA. Integrating crosslinking, exonuclease (exo) trimming, proximity ligation, and high throughput sequencing, SHARC enables transcriptome-wide tertiary structure contact maps at high accuracy and precision, revealing heterogeneous RNA structures and interactions. SHARC data provide constraints that improves Rosetta-based RNA 3D structure modeling at near-nanometer resolution. Integrating SHARC-exo with other crosslinking-based methods, we discover compact folding of the 7SK RNA, a critical regulator of transcriptional elongation. These results establish a new strategy for measuring RNA 3D distances and alternative conformations in their native cellular context.

## INTRODUCTION

RNA plays critical roles throughout the cell, ranging from carrying genetic information to regulation and catalysis ^1^. To perform these tasks, RNA must fold into complex three-dimensional (3D) structures that undergo intricate conformational transitions ^2-7^. Physical methods can be applied to elucidate RNA structure, such as NMR, cryo-EM, and crystallography. These approaches have helped characterize RNA structures, often at atomic resolution, but require well-behaved and purified samples, whereas cellular RNA structures can be highly dynamic and heterogenous^2^. Alternatively, numerous low-resolution approaches, such as chemical mapping and crosslinking, are high-throughput and can be applied in vivo. These low-resolution methods can be coupled with ever-improving computational tools to build 3D models ^8^.

Chemical probing, such as selective 2’-hydroxyl acylation (SHAPE) and dimethyl sulfate (DMS) alkylation, report various aspects of nucleotide flexibility and have been used to constrain local secondary structure predictions ^9-13^. Correlated chemical probing methods such as multiplexed •OH cleavage analysis (MOHCA), mutate-and-map (M^2^), and RNA interacting group mutational profiling (RING-MaP) infer spatial proximity of nucleotides but provides fuzzy distances to constrain 3D modeling ^14-18^. While these methods are improvements over 1D DMS chemical mapping, they are often limited to smaller RNAs as they require the two correlated nucleotides on the same sequencing read, and the sequencing coverage scaled exponentially with RNA length. Furthermore, MOHCA and M^2^ are only applicable to in vitro synthetic RNAs, while RING-MaP is limited by the noisy background and low correlation levels^14^.

Crosslinking and proximity ligation represents an alternative strategy to capture spatial distances among nucleotides, overcoming the limitations of correlated chemical probing ^19^. Recently developed psoralen-crosslinking-based methods, such as PARIS, LIGR-seq, SPLASH, and COMRADES directly capture base pairs either within or between different RNA molecules in high throughput ^20-24^. Psoralen crosslinks staggering pyrimidines in opposite strands through [2+2] photocycloadditions. At the cost of low efficiency, this reaction offers high specificity, is challenging to reverse, and is limited to uridines in helical regions. Even though the gapped reads from such methods can go down to 15 nucleotides on each arm, unambiguous identification of base pairs remains challenging. Recently reported bifunctional acylating crosslinkers, BINARI, reacts with the 2’-OH on all four nucleotides and offers a new approach to capturing nucleotide pairs in spatial proximity crosslinking capacity to 3D space ^25^. However, the nine-step synthesis, large molecular size, and complex reversal mechanism rendered the BINARI compounds unsuitable for cellular application to measure RNA tertiary contacts on a transcriptome-wide level ^25^.

This study develops highly efficient and accessible 2’-hydroxyl acylation chemistry for crosslink-formation and -reversal in living cells (SHARC), overcoming the technical challenges in the preparation and application of BINARI reagents. We develop an exonuclease (exo) trimming approach to pinpoint crosslinked nucleotides, improving the precision of distance measurements to the crosslinked atoms (2’-O in ribose). The integration of SHARC crosslinking, exo trimming, proximity ligation, and high throughput sequencing (SHARC-exo) enables transcriptome-wide analysis of spatial distances between nucleotides at nanometer resolution in cells, without sequence length limitations. We rigorously benchmarked the distance measurement and structure capture using complex, yet well-studied models in cells, such as the ribosome, spliceosome, 7SL, and RNase P, revealing both static structures, interactions, and alternative conformations. The incorporation of distance measurements into Rosetta-based 3D modeling dramatically improved structure resolution. We combined SHARC-exo with established methods, such as PARIS and CLIP, to discover compact folding of the 7SK RNA, a critical regulator of transcriptional elongation in higher eukaryotes. These experiments demonstrate the power of integrating multiple orthogonal approaches to capture proximity constraints in complex RNAs to study their structures. Together, we developed cheap and easily synthesized compounds that dramatically outperform known crosslinking tools, providing the community with a novel strategy for understanding RNA 3D structures and alternative conformations in cells.

## RESULTS

### Quantitative RNA crosslinking with bifunctional 2’-hydroxyl acylation

Unlike proteins, RNA’s overall structure is governed by sparse tertiary contacts (**Fig. 1a**). Highly structured RNAs as large as 500 nucleotides may only contain a few critical tertiary contacts ^6,26-28^. Knowledge of these tertiary contacts significantly improves modeling of complex RNAs ^15,18,29^. To determine these constraints for RNAs in cells, we sought to develop a new set of bifunctional and reversible 2’-hydroxyl acylation reagents with flexible linkers (**Fig. 1b**). To improve accessibility and facilitate optimizations, we focused on a modular design using simple dicarboxylic acids, the length of which can be easily adjusted. Subsequent reversal allows facile analysis of the crosslinked sequences. To synthesize such reagents, we activated simple dicarboxylic acids using 1,1’-carbonyldiimidazole (CDI) in a one-step reaction (**Fig. 1c**, see methods in Supplementary Information). We tested crosslinking efficiency on a model self-complementary duplex RNA **1** *in vitro*, where acylations are expected to occur on unconstrained nucleotides (**Fig. 1d**).

**Figure 1.**
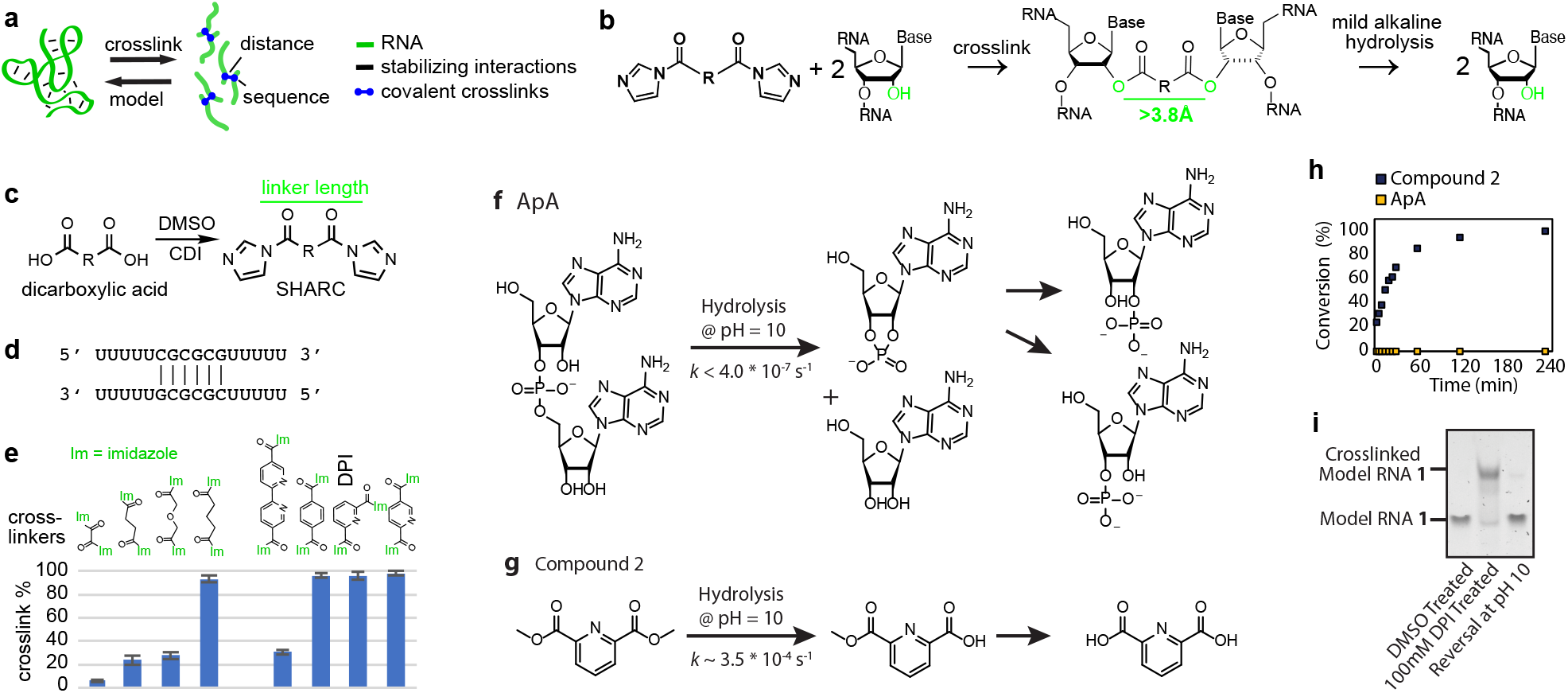
Efficient RNA crosslinking and reversal using SHARC reagents. **a**, Principle of using crosslinking to determine RNA 3D structures. Spatial distances among nucleotides in an RNA should be sufficient to rebuild its 3D structure. **b**, SHARC crosslinking and reversal of ribose 2’ hydroxyls. **c**, One-step activation of dicarboxylic acids to produce SHARC reagents. CDI: carbonyldiimidazole. **d**, Model RNA 1 homodimer duplex for testing SHARC crosslinking and reversal. **e**, Crosslinking efficiency of a series of SHARC reagents on the model RNA 1 duplex. Common names for the 9 dicarboxylic acids to prepare the crosslinkers: oxalic, succinic, diglycolic, glutaric, 6,6’-binicotinic, terephthalic, dipicolinic and isocinchomeronic. Crosslinking condition: MOPS buffer (pH 7.5 0.1 M KCl, 6 mM MgCl_2_), 4 hours at room temperature. **f**, Hydrolysis kinetics of RNA phosphodiester bonds in a model dinucleotide ApA. **g**, Hydrolysis of a model SHARC crosslinking product, compound 2. **h**, Measurements of hydrolysis rates for the ApA and model compound 2 (5mM starting concentration). **i**, Example SHARC crosslinking and reversal of model RNA 1 duplex on a 20% urea-denatured TBE polyacrylamide gel.

We activated a set of eight dicarboxylic acids with diverse linker lengths and chemical properties (**Fig. 1e**, see characterization in **Supplementary Fig. 1, 2a**). The efficiency of crosslinking RNA **1** was measured by polyacrylamide electrophoresis (**Fig. 1e, Supplementary Fig. 2b**). Activated oxalic and succinic acids showed low to modest crosslinking of 1-24% (**Fig. 1e**), possibly due to the short linker lengths that might be insufficient to bridge 2’-OH groups on opposing strands (see estimated crosslinker lengths in **Supplementary Fig. 2a**). Activated glutaric acid showed 94% crosslinking, while diglycolic acid, which is similar in size, exhibited significantly lower crosslinking of 27%. The differences between these aliphatic compounds can potentially be explained by the inductive effect of the beta oxygen that substantially increases the reactivity of the activated ester ^30^, making it more susceptible to hydrolysis. The activated aromatic compounds, terephthalic, isocinchomeronic, and dipicolinic acids all exhibited excellent crosslinking efficiencies between 97-99%, likely due to optimal spacing and conformation of the linkers to bridge opposing 2’-OH groups and favorable reactivity towards 2’-OH moieties as previously demonstrated by aromatic SHAPE reagents ^13^. The activated bipyridine compound 6,6’-binicotinic acid showed an apparent crosslinking efficiency of 31%, though solubility in aqueous solution was limited, hampering the exact determination of its crosslinking performance. We selected dipicolinic acid imidazolide (DPI) as a candidate to test further based on these results. To characterize the reaction kinetics of DPI at physiological pH 7.4 at room temperature, we measured its hydrolysis with NMR and found that 50% was hydrolyzed after 5 min (**Supplementary Fig. 2c-e**). This reaction is significantly faster than the structurally related cell-permeable SHAPE reagent NAI (half-life ∼30 min, HEPES buffer pH = 8.0) due to the additional electron-withdrawing group ^31^, suggesting its potential for rapid RNA crosslinking in cells.

### Reversing 2’-hydroxyl crosslinking under mild alkaline conditions without RNA damage

Reversal of the crosslinks is necessary for subsequent sequence analysis. We hypothesized that the lower stability of the 2’-acylation products relative to the phosphodiester bonds could allow selective crosslink reversal without causing RNA chain breaks. To test this, we first analyzed the rate of phosphodiester cleavage in a model RNA dinucleotide ApA (**Fig. 1f**). We compared it to the methyl ester of dipicolinic acid **2** (**Fig. 1g**), a simple ester derivative of the SHARC reagent DPI. The two compounds were incubated in a 3:1 mixture of 100 mM borate buffer and DMSO at pH 10.0, and the stability was monitored over time by ^1^H NMR (**Fig. 1h** and **Supplementary Fig. 2f-g**). No degradation of ApA was observed even after 48 hours, and the rate constant was estimated to be below 4.0*10^−7^ s^-1^ (**Fig. 1f**). In contrast, compound **2** was fully hydrolyzed after ∼120 min, with a rate constant of 3.5*10^−4^ s^-1^ (R^2^=0.99) (**Fig. 1g**). From this we concluded that the ∼1000-fold difference in rate constant should provide sufficient opportunity to selectively reverse the crosslinks under mild alkaline conditions without RNA damage.

To investigate if the alkaline conditions can be successfully applied to reverse SHARC crosslinks in longer RNA, the model RNA **1** was crosslinked with DPI, purified, and 10 µM of crosslinked RNA was incubated in 100 mM Borate buffer pH 10.0 for two hours at 37 °C. The crosslinked RNA was nearly fully reversed without apparent degradation (**Fig. 1i**). Increasing pH to 11.0 did not result in noticeable degradation, suggesting a broad window for robust reversal of crosslinks (**Supplementary Fig. 2h-i**). Together, we showed for the first time that 2’-OH acylation could be easily reversed at moderately alkaline pH without significant RNA damage, opening the possibility for subsequent sequence analysis in various applications. For larger RNAs, degradation may be unavoidable. However, fragmentation is an inherent step in sequencing library preparation, so the residual RNA degradation does not affect subsequent sequence analysis.

### Exonuclease trimming: a new strategy to determine crosslinking sites at near nucleotide resolution

Having demonstrated efficient SHARC crosslinking and reversal, next we developed a new strategy, exonuclease (exo) trimming, to measure inter-nucleotide distances, based on our previously established PARIS method ^21,32^ (**Fig. 2a**). Crosslinked RNA samples are first digested with RNase III, which fragments both single and double-stranded RNA into short pieces ^21^. RNA fragments are fractionated on a denatured-denatured 2-dimension (DD2D) gel ^24^, where the second dimension is denser than the first (e.g., 16% vs. 8%). The differential gel densities enable the separation of crosslinked from non-crosslinked fragments. The crosslinked fragments migrate as a smear above the diagonal (**Fig 2a**), therefore achieving near 100% purity without the contamination of RNA with monoadducts of the crosslinker. The purified cross-linked fragments are then trimmed by an exonuclease, e.g., RNase R, which removes nucleotides from the 3’ end until it is blocked by the crosslink sites ^33^. The trimmed fragments are ligated so that the two arms are joined to form a continuous RNA molecule. After mild alkaline crosslink reversal, the bipartite RNA molecules are reverse transcribed for cDNA library preparation and sequenced. The gapped reads are clustered into duplex groups (DGs, similar to our previous definition ^21,34^, but includes all gapped reads from secondary and tertiary structures). Each group corresponds to one specific pair of nucleotides that are close to each other. The gapped reads should reveal trimmed 3’ ends at a fixed distance from the actual crosslinking sites (∼5 nts, see details below). The spatial distances between the crosslinked nucleotides (the 2’-OH groups, to be precise) are determined by the length of the linkers and the flexibility of the RNA structure.

**Figure 2.**
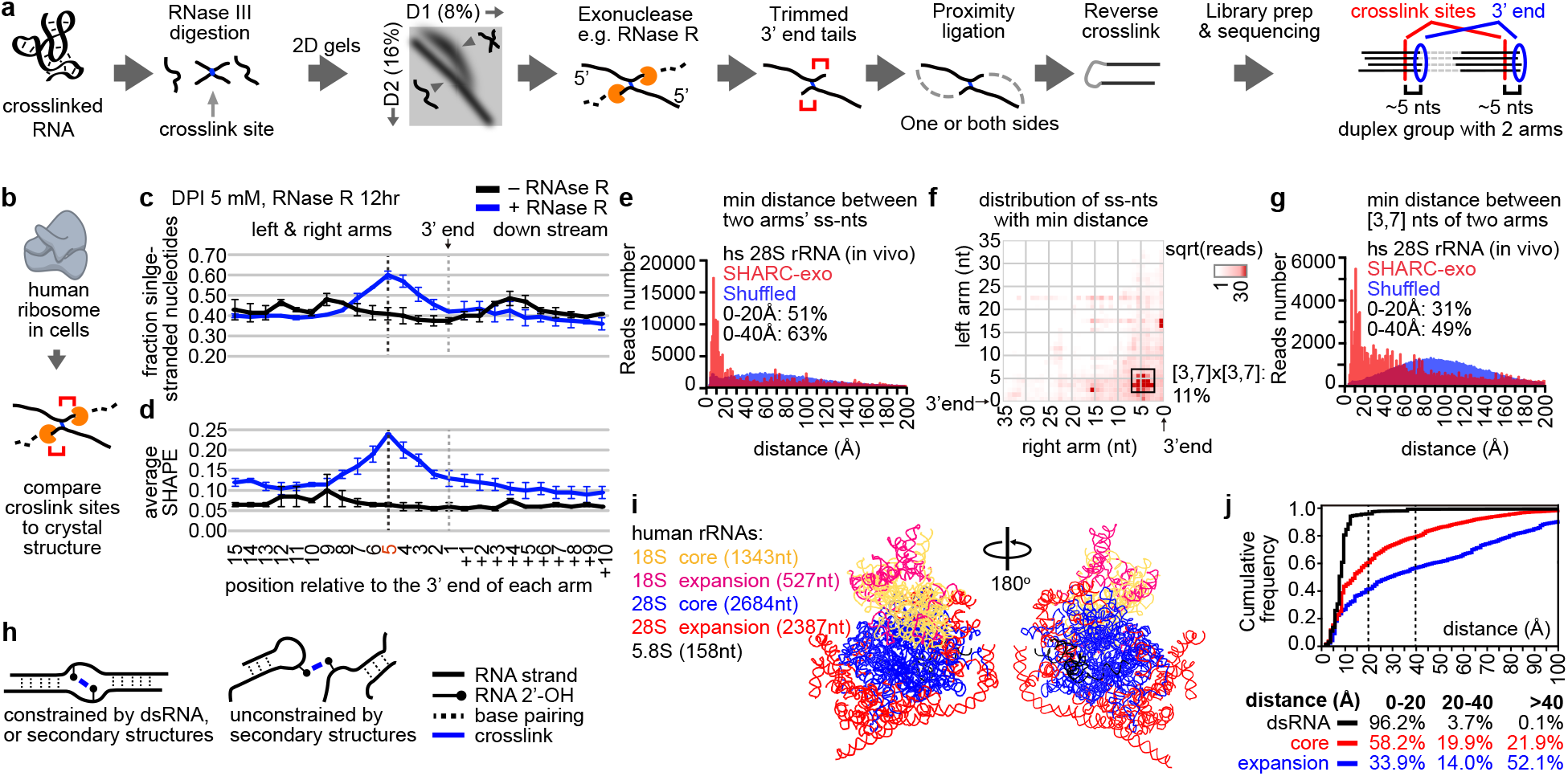
Exonuclease trimming measures spatial distance at nucleotide resolution in cells. **a**, RNA samples, either in vitro or in cells, are crosslinked, digested to short fragments using RNase III, and the crosslinked fragments isolated using a DD2D gel. Isolated crosslinked fragments are then trimmed with exonucleases, proximally ligated, decrosslinked, ligated with adapters for cDNA library preparation. The putative crosslinking sites are determined based on the trimmed 3’ ends. **b**, Benchmarking SHARC-exo against the human ribosome in cells. **c-d**, Identification of SHARC crosslinking sites in the human ribosome from crosslinked cells. SHARC-exo condition: 5mM DPI and 12 hour RNase R trimming. Error bars represent standard deviations from 2 biological replicates. **c**, Fraction of single stranded nucleotides were calculated along each arm of the gapped reads. **d**, Average icSHAPE signal was calculated along each arm of the gapped reads. **e**, Distribution of minimal distances between single-stranded nucleotides (ss-nts) in the two arms of each gapped read for the human 28S ribosomal RNA. The blue histogram is the distribution for randomly shuffled reads. **f**, Positions of single-stranded nucleotides that are closest to each other on the two arms of each read. Square root of reads were plotted. All reads were aligned relative to the 3’ end. Positions 3 to 7 for each arm were boxed, which represent 11% of total coverage. **g**, Distribution of minimal distances between nucleotides in the 3 to 7 range on both arms. The blue histogram is the distribution for randomly shuffled reads. **h**, Two types of SHARC crosslinked spatial proximity. **i**, Illustration of the core and expansion segments (ESs) in 3 human cytoplasmic rRNAs. **j**, Cumulative distribution of the minimal distances between the two arms of each read for reads mapped to different types of structures. Only non-dsRNA, or tertiary contact, reads are used for the core and expansion segments. Note, the color coding in panels i and j have different meanings.

To validate the exo trimming approach, we first applied it to PARIS experiments, where the well-established crosslinking preference of psoralen enables rigorous testing of the trimming efficiency. After RNase R treatment, the reads are significantly shorter (**Supplementary Fig. 3a**). Counting from the 3’ end of each arm of the gapped reads, we observed a strong enrichment of uridines at the 3^rd^ to 6^th^ position, peaking at the 5^th^ nucleotide, suggesting that the psoralen crosslinking to uridine blocked RNase R trimming, leaving ∼5 nts at the 3’ end (**Supplementary Fig. 3b**). In contrast, no enrichment of uridine was observed at the exact location without trimming. Therefore, the exo trimming strategy allows us to pinpoint the crosslinking sites with high precision. As an example, we showed the identification of crosslinking sites in helical regions of the 28S rRNA (**Supplementary Fig. 3c-e**).

### SHARC-exo accurately measures static and dynamic spatial distances in RNA in cells

To test SHARC-exo, we first crosslinked HEK293 cells with DPI, extracted RNA using proteinase K (PK) and the TNA method, fragmented RNA with RNase III, and isolated the crosslinked fragments using the DD2D gel method. We recovered 1.01%, 1.31% and 1.89% RNA fragments as crosslinked using 5, 12.5, or 25 mM DPI, respectively (**Supplementary Fig. 3f-h**, comparable to psoralens used in the PARIS method) ^24^. PK digests proteins to <6 amino acids long ^35^, and efficiently removed the vast majority of proteins (**Supplementary Fig. 3i**, TNA method), therefore the detected interactions are unlikely to be mediated by proteins. We sequenced the SHARC-exo libraries and observed 3.3-14.5% of the reads are gapped, similar to PARIS (**Supplementary Table 1**) ^20,21^. Crosslinked reads are highly reproducible at different DPI concentrations and RNase R trimming conditions (**Supplementary Fig. 3j**). The two arms of each gapped read span a wide range of distances, for example, up to the entire length of rRNAs (1869 and 5070 nucleotides, respectively, **Supplementary Fig. 3k**). The crosslinked Together, these results demonstrated efficient and robust SHARC crosslinking of RNA in cells.

To test the ability of SHARC-exo in measuring spatial distances, we focused on the ribosome due to its high abundance, complex structures, and intermolecular interactions (**Fig. 2b**) ^36,37^. Given that RNA homodimers are rare due to the transient nature of most intermolecular interactions ^38^, all subsequent analysis of structures in individual RNA species were performed under the assumption that the gapped reads were derived from the same RNA, instead of two identical RNA molecules. We first calculated the fraction of single-stranded nucleotides close to the 3’ end of each arm of the gapped reads, based on the ribosome cryo-EM structure ^36^ (**Fig. 2c-d**, and **Supplementary Fig. 4a-d**). Counting from the 3’ end, trimmed samples exhibited a dramatic increase in the fraction of single-stranded nucleotides between the 1^st^ and 8^th^ nucleotides, with a peak at the 5^th^ (**Fig. 2c**, ∼1.3-fold over non-trimmed). We observed a similar trend when using experimentally determined icSHAPE reactivities for the ribosome ^21^ (**Fig. 2d**). The stronger enrichment of icSHAPE signal (**Fig. 2d**, ∼ 3.7-fold) compared to the counts of single-stranded nucleotides further confirmed the selective crosslinking of unconstrained nucleotides by DPI and the efficient trimming. A/U nucleotides are slightly enriched near the crosslink sites, likely reflecting their lower base-pairing potential (**Supplementary Fig. 4e**)

To determine the range and precision of distance measurements by SHARC-exo, we calculated spatial distances between the two arms of each gapped read in the ribosome cryo-EM model ^36,37^. The minimal distance has a narrow distribution with a long tail, where 51% are within 20 Å, with a mode of ∼8 Å, close to the physical length of the crosslinker (∼7 Å) (**Fig. 2e, Supplementary Fig. 2a**). In contrast, the distances for randomly shuffled reads have a much broader distribution (Wilcoxon rank sum (WRS) test, p<10^−300^). To determine whether trimming precisely reveals the crosslinked nucleotides, we searched for nucleotides along each arm closest between the two arms (**Fig. 2f**). Not surprisingly, the nearest point is the 5^th^ nt, consistent with the highest SHAPE reactivity (not seen on the non-trimmed SHARC data, **Supplementary Fig. 5**). The distance between 5^th^ +/- 2 nts on the two arms follow a narrow distribution, with a mode distance of 8 Å, and 31% reads less than 20 Å, and 49% less than 40 Å (WRS test p<10^−300^, compared to shuffled reads, **Fig. 2g**).

The ribosome is a highly dynamic and flexible macromolecular machine. SHARC-exo captures spatial distances of the ribosome in its entire life cycle in cells that include both intra-ribosome dynamics and inter-ribosome contacts. To understand the long tails in distributions (in **Fig. 2e, g**), we separated distances into ones constrained by extensive base pairs, which are more stable, and those simply in spatial proximity (tertiary motifs), which are more dynamic (**Fig. 2h**) ^39^. We further split tertiary contacts to the stable core and the more flexible Expansion Segments (ESs), many of which are not resolved with cryo-EM (**Fig. 2i**). As expected, distances constrained by secondary structures are predominantly within 20 Å (dsRNA, 96.2%, **Fig. 2j**), whereas the core and ES tertiary distances have increasingly broader distributions (58.2% and 33.9% within 20 Å, respectively). Together, this analysis further demonstrated SHARC-exo’s high accuracy and the ability to capture heterogeneous conformations in cells.

To test the robustness of SHARC-exo, we compared multiple DPI concentrations and trimming conditions. Regardless of DPI concentration, SHARC-exo produced consistent enrichment of single-stranded nucleotides near the 5^th^ nucleotide (**Supplementary Fig. 4a-d**). The minimum distances between the two arms are primarily within 20 Å (**Supplementary Fig. 5a**). However, higher DPI concentrations reduced trimming efficiency, which can potentially be explained due to disruption of the endogenous structure or monoadducts that block trimming. The monoadducts may reduce the resolution in distance measurements. At the same DPI concentration, heavier trimming increased the resolution of spatially proximal nucleotides (**Supplementary Fig. 5b-d**).

### SHARC-exo analysis of RNA structures and interactions in vivo

To test the ability of SHARC-exo in capturing known structures, we extracted spatial distances within 20 Å in the ribosome (**Fig. 3a-b**, left panels). SHARC-exo measurements (upper right triangles) are highly consistent with distances between icSHAPE-reactive nucleotides in the cryo-EM model for both the 18S and 28S rRNAs (lower left triangles). Shuffling of SHARC-exo reads resulted in random distributions (**Fig. 3a-b**, right panels). The zoom-in views of the two regions showed both highly consistent distance measurements and ones missed by SHARC-exo (blue and red boxes in areas 1-2, **Fig. 3c-d**). The missed spatial proximities likely represent tight ribosome regions inaccessible to DPI or nucleotides with steric hindrance.

**Figure 3.**
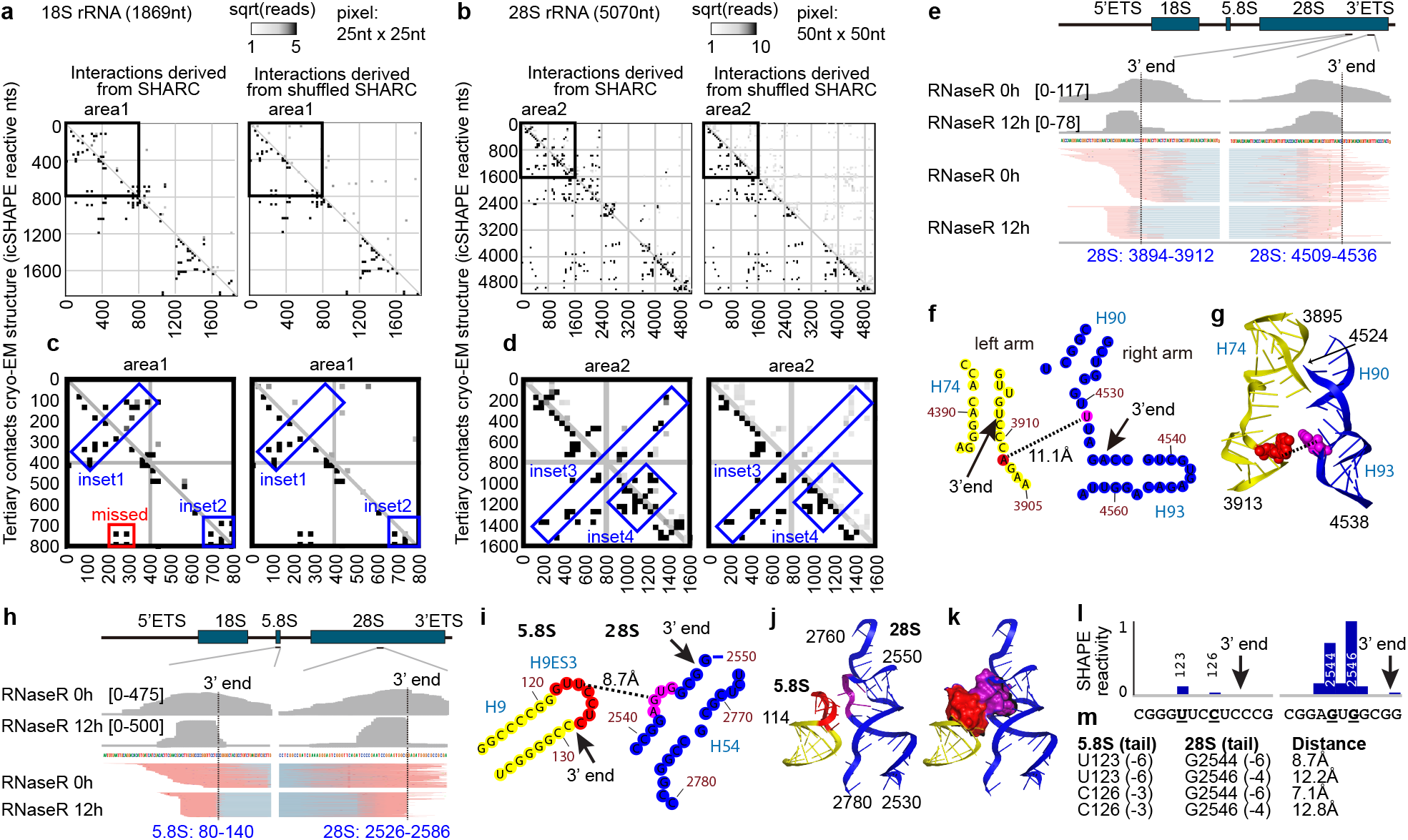
SHARC-exo captures spatial distances within the human ribosome. **a-b**, Comparing SHARC-exo captured spatial proximities to a cryo-EM structure model of the human ribosome in 25nt x 25nt (**a**) or 50nt x 50nt (**b**) windows (PDB:4V6X). icSHAPE-measured reactive nucleotides within 20 Å of each other in the 18S (**a**) and 28S (**b**) rRNAs are plotted on the lower left corner of each square. SHARC-exo gapped reads with 2 arms within 20 Å of each other are plotted on the upper right corner, and the read numbers are square-root scaled. Positions of SHARC-exo reads are randomly shuffled as a control (right panels). **c-d**, Zoom-in views of two areas in the 18S and 28S rRNAs. Blue rectangles highlight consistent distance measurements between SHARC-exo and cryo-EM. Red rectangle highlight regions missed by SHARC-exo. **e**, An example tertiary proximity in the 28S rRNA captured by SHARC-exo. **f**, Secondary structure model of the two regions, showing the consensus 3’ ends of the gapped reads, expected crosslink sites and distance. **g**, 3D structure model of the crosslinked sites. **h-m**, SHARC-exo captured an interaction between 5.8S and 28S rRNAs. **h**, Gapped reads for the interactions between 5.8S and 28S rRNA. **i**, The 3’ ends, putative crosslinking sites and distance mapped onto the secondary structure model of rRNAs. **j-k**, 3D model of the interaction, where the two loops involved in interactions are shown in red and purple. Interhelical stacking is shown in spheres (k). **l-m**, icSHAPE measurement of nucleotide flexibility around the crosslinking sites (l) and all possible distances between the two interacting loops (m). **m**, Distances between 2’OH groups at the nucleotides with high SHAPE reactivity. Tail length in the parentheses indicate the distance between the 3’ ends and the reactive nucleotides that are potentially crosslinked.

SHARC-exo captured spatial distances both constrained by secondary structures or simply in spatial proximity (**Fig. 3e-m**, see more examples in **Supplementary Fig. 6**). In one instance in the 28S rRNA, RNase R trimming resulted in significantly shortened 3’ ends for both arms of the DG (**Fig. 3e**). Tracing back to the 5^th^ nucleotides from the 3’ ends, where the crosslinks are expected, we obtained a pair of nucleotides with a spatial distance of 11.1 Å between the 2’ oxygens in the ribosome cryo-EM model (**Fig. 3f-g**). SHARC-exo also captured intermolecular interactions. For example, SHARC-exo precisely mapped a tertiary contact between the loop on 5.8S helix 9 ES 3 (H9ES3) and an internal bulge on 28S H54 (**Fig. 3h**). This interaction is stabilized by base stacking between 5.8S U126 and 28S G2544 (**Fig. 3i-k**). These two stretches of single-stranded nucleotides have significantly higher icSHAPE reactivity than the surrounding helical regions (**Fig. 3l**). The spatial distances among the most reactive two nucleotides on each side range from 7.1 to 12.8 Å. Their distances to the SHARC-exo determined 3’ ends range from 3 to 7 nucleotides, consistent with the global average for SHARC-exo (**Fig. 3m**). Together, these results demonstrate that SHARC-exo can capture static and alternative RNA-RNA interactions at near nucleotide resolution.

In addition to the ribosome, SHARC-exo also captured spatial distances in other non-coding RNAs, including the RPPH1 RNA in RNase P, the 7SL RNA in signal recognition particle (SRP), and U4/U6 snRNAs in the spliceosome (**Supplementary Figs. 7-9**). RNase P is a ribozyme that cleaves off the 5’ leader of tRNA precursors ^40,41^. SHARC-exo captured five proximal nucleotide pairs in the range of 17-36 Å (compare to ∼190 Å -- the overall length of RPPH1 structure. **Supplementary Fig. 7**). In the 7SL RNA, all SHARC-exo measured distances are in the range of 9-26 Å, except one at 77.5 Å, which is likely due to an alternative conformation previously predicted as a precursor in the SRP assembly ^42^ (**Supplementary Fig. 8**). U4 and U6 snRNAs form a stable complex in the spliceosome, and two DGs connecting U4 to U6 were detected (**Supplementary Fig. 9a**) ^43^. Crosslinking sites were mapped to two regions in spatial proximity, including a 3-way junction (DG1, **Supplementary Fig. 9b-c**) and single-stranded regions near an intermolecular helix (DG2, **Supplementary Fig. 9b**,**d**). In both structures, exo trimming pinpointed the nucleotides in spatial proximity. Together these results demonstrated that SHARC-exo could measure spatial distances in a wide variety of RNAs in cells.

### SHARC-exo distance measurements improve Rosetta-based RNA 3D modeling

Having demonstrated accurate distance measurements by SHARC-exo, next, we investigated whether these constraints can improve 3D structure prediction. For example, we focused on a specific region, h22-h24, in the 18S rRNA (**Fig. 4a, Supplementary Fig. 10a**). SHARC-exo captured two major spatially proximal pairs of nucleotides at 7.8 and 21.0 Å (**Fig. 4b, Supplementary Fig. 10b**). Using these two distances as constraints and a linear pseudo-energy function, we modeled the 3D structure of this 18S segment (see methods in Supplementary Information). The addition of the constraints significantly reduced the RMSD distribution for all models and the top 200 models (**Fig. 4c-d, Supplementary Fig. 10c**). Clustering showed that the SHARC-exo constrained top model displays high topological similarity to the cryo-EM structure, while the de novo model deviates substantially (**Fig. 4e-f, Supplementary Fig. 10d**). In the native ribosome, the h22-h24 region is stabilized by interactions with other RNA and protein components. Despite using only two constraints in this unfavorable case, the resolution of the 3D model increased significantly. The availability of deeper sequencing coverage and denser constraints will likely further improve the resolution of 3D modeling.

**Figure 4.**
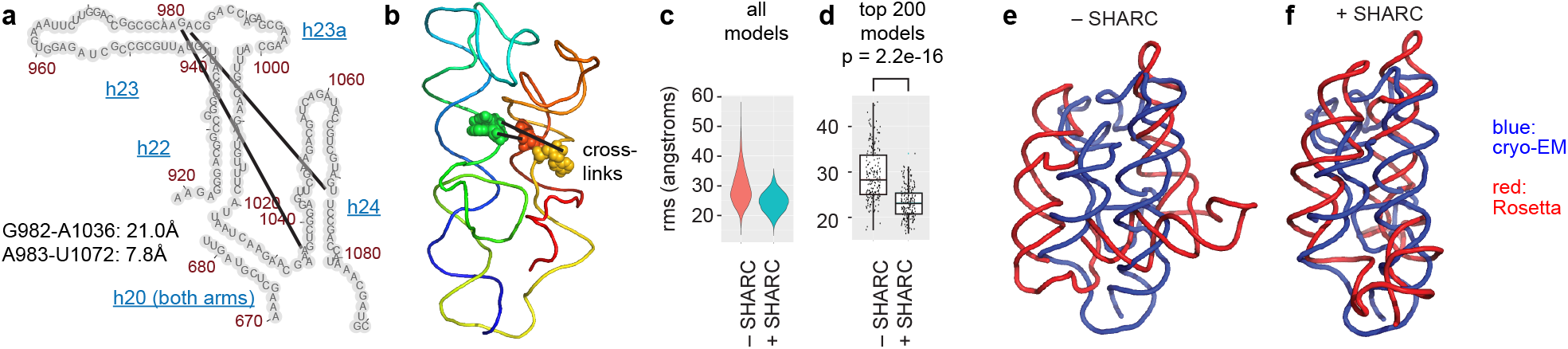
Spatial distances captured by SHARC-exo improve 3D modeling. **a**, Secondary structure model of h22-h24, showing the two crosslinks and their distances. Only the right side of h20 is included for modeling. **b**, 3D model of h22-h24, showing the two crosslinks (PDB: 4V6X). **c**, Distribution of the RMSD values for all models with (n=12,363) or without (n=20,394) SHARC-exo constraints. **d**, Top 200 models are shown in box plots. p-value was calculated using the Wilcoxon ranksum test. **e**, Comparing the top model from the de novo Rosetta run (red) with the cryo-EM model (blue). **f**, Same as panel e, except that the Rosetta model was constrained with the two SHARC-exo distances measurements.

To further benchmark the SHARC-exo method, we performed in vitro crosslinking of a well characterized model RNA, the P4-P6 domain of the tetrahymena ribozyme ^6^. SHARC-exo revealed 17 DGs with highly variable abundance, where the low abundance ones are likely artifacts from RNA mis-folding or crosslinking (**Supplementary Fig. 10f**). We focused on 5 abundant DGs where each contain >3% of total reads. The RNase R trimming shortened the 3’ ends and improved the measurement of spatial distances by 11-23 Å, based on the analysis of the crystal structure ^6^ (**Supplementary Fig. 10g-h**). The RNase R refined distances, in the range of 19-41 Å, were still longer than the minimal crosslinkable distances between the two arms (e.g. **Supplementary Fig. 10i-j**). These results demonstrate that the crosslinking and RNase R trimming successfully captured spatial distances, but there is still space for further improvement. The addition of the SHARC and SHARC-exo derived distances significantly reduced overall RMSD distribution for complete set of all models and the top 100 (**Supplementary Fig. 10k)**. Models constrained by SHARC-exo distances are much more compact than non-constrained or SHARC-constrained ones (**Supplementary Fig. 10l-o)**. This result further confirmed the usefulness of distance measurements in 3D modeling.

### SHARC-exo captures alternative RNA conformations

Many flexible regions in the ribosome, especially the ES, play essential roles in translation ^44,45^. However, they are often at low resolution or not resolvable with crystallography or cryo-EM due to their dynamic nature ^36^. In SHARC-exo data, reads with two arms that span >40 Å are predominantly located in the ES (96.52%, vs. 3.41% for core tertiary, and 0.08% for dsRNA, **Fig. 5a, Supplementary Fig. 11a-c**). Consistent with this, the ESs, even if visible by cryo-EM, have a higher B-factor, indicating higher flexibility, while the core segments are considerably better resolved (average resolution 5.4 Å, ranges between 2.2 and 21 Å, **Supplementary Fig. 11d-e**) ^36^. Two ESs in the 28S rRNA, 78ES30 and 79ES31, have the highest read coverage with between-arm distances >40 Å (hub1 and hub2, **Fig. 5b-c**). Hub1 makes extensive contacts on many regions on the ribosome, most of which are other flexible ESs (**Fig. 5d, Supplementary Fig. 11f**). For example, the top 6 DGs connecting hub1 are all located on the surface of the ribosome, among which the top-ranked is hub2 (**Fig. 5d, Supplementary Fig. 11g**). The flexibility of both hub1 (78ES30) and its partners make it possible for them to reach each other. Using Rosetta and a single distance constraint between hub1 and hub2, we found that the two regions can be modeled in spatial proximity (from 128 to 16 Å, **Fig. 5e-f**) without any clashes with other parts of the ribosome surface (**Supplementary Fig. 11h-i**).

**Figure 5.**
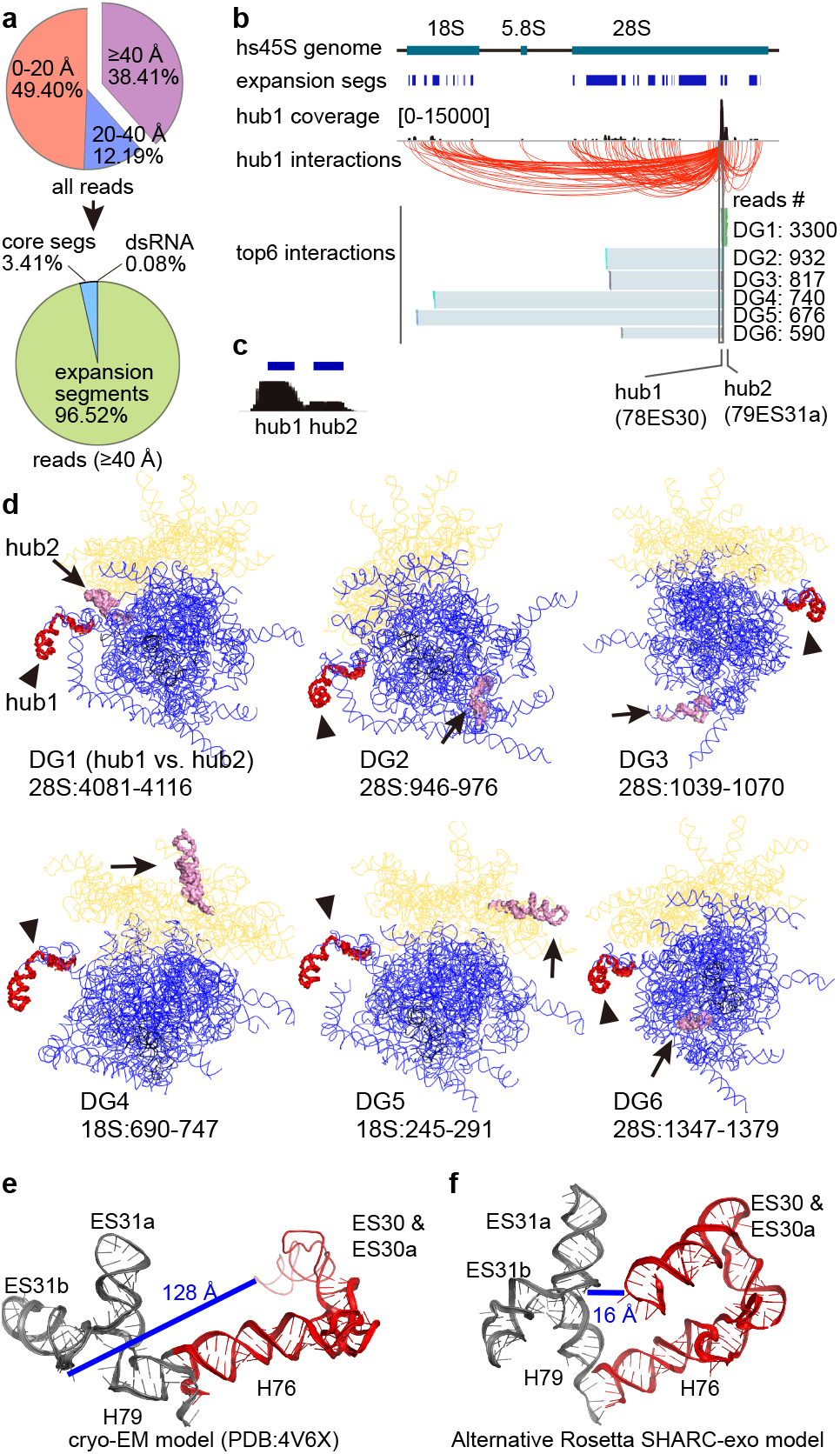
SHARC-exo reveals dynamic conformations of expansion segments in the ribosome. **a**, Distribution of reads based on their distances between the two arms. For reads with minmal distance >40 Å, the vast majority are mapped to the expansion segments (lower panel). **b**, All hub1 interactions are shown by red arcs. Among top ranked DGs connecting hub1, the highest abundance expansion segment (78ES30), 4 of them are within the 28S (DGs 1-3 and 6), and 2 of them with the 18S (DGs 4 and 5). **c**, Zoom-in view of hub1 and hub2, the two most highly connected dynamic regions in the rRNAs. **d**, Locations of the hub1 (red colored, indicated by arrowheads) and its targets (gray, indicated by arrows). Blue line: 28S rRNA. Yellow line: 18S rRNA. Black line: 5.8S rRNA. **e-f**, Cryo-EM (**e**) and a representative Rosetta (**f**) model of the hub1-hub2 region (28S:3936-4175) in the ribosome. Distances are between the 2’OH groups at nucleotides 4000 and 4123. **f**, The Rosetta model was constructed with a single constraint between nucletoides 4000 and 4123.

Next, we examined more complex RNA-RNA interactions between the 5.8S and 28S and between 18S and 28S rRNAs (**Supplementary Fig. 12-13**). We discovered both spatial proximal nucleotide pairs and distant ones that likely represent intermediates during ribosome assembly. Two significant regions in the 5.8S interact extensively with the 28S (**Supplementary Fig. 12a**). Among the top 6 DGs connecting 5.8S and 28S, DGs 1, 4, and 5 captures direct contacts, DGs 3 and 6 are likely due to the alternative conformations of ESs on the 28S that allow the formation of intermolecular contacts, which were not captured by cryo-EM, underlining the power of the SHARC method (**Supplementary Fig. 12b**) ^36^. The remaining DG2 connects two regions that cannot reach each other in the mature ribosome but are supported by extremely high sequencing coverage (**Supplementary Fig. 12a**, close to DG1). This is likely explained due to spatial proximity during the assembly of the ribosome. The interactions that we captured between 18S and 28S expansion segments suggest a highly dynamic nature of the translation machine (**Supplementary Fig. 13**). Together with the alternative conformations in 7SL (Supplementary Fig. 8), these results suggest that SHARC-exo captures static and dynamic structures in cells.

### SHARC-exo reveals compact folding of the 7SK RNA

The noncoding RNA 7SK plays an essential role in transcriptional regulation ^46,47^. Still, the structural basis of its function is largely unknown, except for a few small regions that were solved by crystallography and NMR ^48-53^. For the full-length 7SK, 331 nt in humans, both secondary and tertiary structures remain uncertain. Wassarman and Steitz proposed the first secondary structure model with four major helices, a “linear model” based on chemical probing (**Supplementary Fig. 14a-b**) ^54^. Deep phylogenetic analysis together with manual adjustments revealed a consistent global secondary structure model across metazoans (Marz model, or “circular model”), featuring eight helical regions, among which a terminal helix (M1) circularizes 7SK ^55^. More recent work using the evolutionary coupling method that detects spatial interactions failed to identify the M1 terminal helix (**Supplementary Fig. 14c-e**) ^56^. In vivo icSHAPE ^21^, a measurement of 1D nucleotide flexibility, only provided consistent but not conclusive evidence for the overall validity of helical regions in the Marz model (**Supplementary Fig. 14f-g**). Here, using SHARC-exo in combination with low-resolution methods PARIS and CLIP, we conclusively demonstrate the existence of the circular model and extensive tertiary contacts within this RNA that suggest compact 3D folding.

Using SHARC-exo, we discovered extensive secondary and tertiary contacts among the helices and single-stranded regions (**Fig. 6a, Supplementary Fig. 15**). These contacts suggest tight folding of the 7SK RNA in cells. In particular, the two most extended helices, M3 and M7, are packed together (a subset of the contacts shown in **Fig. 6b**). To validate the compact folding of 7SK, we reanalyzed our recent PARIS and previously published eCLIP data ^21,57^ (**Fig. 6c-d, Supplementary Fig. 16-17**). PARIS validated the local structures in the Marz model in both human and mouse cells (M3, M4/M5, and M7), especially the terminal helix M1 (DG1 in **Supplementary Fig. 16a-c**). In addition, PARIS revealed proximity between distant regions (**Supplementary Fig. 16c-e**, DGs 2-3). These long-range contacts suggest direct contacts between M3 and M7 since psoralen crosslinking requires stable structures, where at least two base pairs are needed to sandwich a psoralen molecule ^58^. In addition to 2-segment (1-gap) reads that represent RNA duplexes, PARIS also captures more complex structures in the form of multi-segment reads, where two structures that form together in one molecule are crosslinked ligated and sequenced (**Fig. 6e, Supplementary Fig. 16f**) ^34,38^. Therefore, multi-segment reads provide direct evidence that three or more segments are close to each other in space in the same RNA molecule. The multi-segment reads connect the 5’ end M3 to the 3’ end M7 and their surrounding sequences. Together, these PARIS data suggest compact folding of the 7SK RNA.

**Figure 6.**
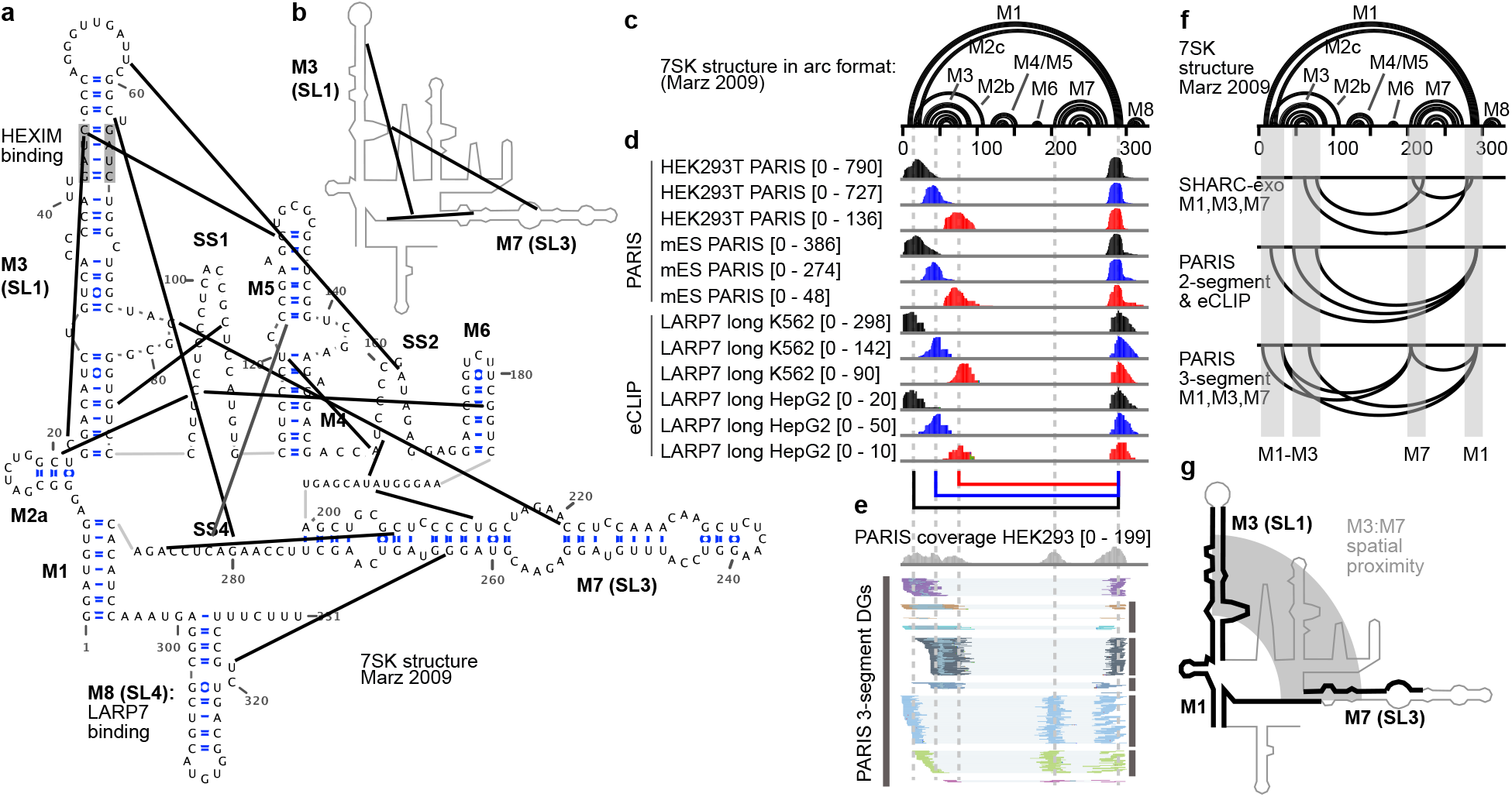
SHARC-exo, PARIS and eCLIP reveal compact folding of the 7SK RNA. **a**, Secondary structure model of the human 7SK RNA (Marz 2009, helices M1-M8 and single stranded regions SS1-4) and 15 SHARC-exo derived spatial constraints (thick black lines). HEXIM and LARP7 binding sites are labeled. **b**, SHARC-exo derived tertiary proximities between M3 and M7. **c**, Marz 2009 secondary structure model of 7SK in arc format. **d**, eCLIP and PARIS captures long-range contacts among M1, M3 and M7. Each track shows the coverage of one DG connecting two regions. **e**, PARIS two-gap (3-segment) reads that support long-range contacts in the compact 7SK RNA. The 5 vertical dash lines align the major peaks interacting with each other. **f**, Comparison of long-range contacts derived from SHARC-exo, PARIS 2-segment, eCLIP 2-segment and PARIS 3-segment reads. **g**, Model of spatial proximity between M3 and M7 as determined by all 3 methods.

CLIP experiments occasionally crosslink a protein molecule to more than one RNA fragment in spatial proximity ^59^. Proximity ligation can join these fragments in one sequencing read (**Fig. 6d, Supplementary Fig. 17**). We reanalyzed the extensive collection of eCLIP datasets and found that LARP7, an integral component of the 7SK complex, is strongly crosslinked to multiple locations, including the M1, M3, and M7-M8 (**Supplementary Fig. 17a-b**). The LARP7 eCLIP gapped reads confirmed the local 7SK secondary structures (M3, M6, and M7), the terminal helix M1. They revealed long-range structures that bring M3 and M7 to spatial proximity, similar to PARIS (**Fig. 6d, Supplementary Fig. 17c-d**, DGs 1-3). Together, our integration of 3 orthogonal approaches, SHARC-exo, PARIS, and eCLIP, provided strong support for the circular model (Marz model) of the 7SK secondary structure and suggest a compact folding of the 7SK helices, in particular the direct contacts between M3 and M7 (**Fig. 6f-g**).

## DISCUSSION

This study reports a series of new reversible crosslinkers, SHARC, that can capture spatial proximity in RNA with high efficiency. We develop a new exo trimming strategy that improved resolution in both SHARC and PARIS to near single nucleotides (∼3-7 nucleotides from the 3’ end), therefore generally applicable to various types of crosslinkers. The high throughput SHARC-exo method measures spatial distances between nucleotides either within an RNA or between different RNA molecules in living cells with high efficiency. We show that SHARC-exo distance information can be used to constrain Rosetta-based 3D RNA modeling, therefore opening the possibility of understanding the 3D structures of the entire transcriptome in vivo. Using the ribosome as an example, we demonstrate that SHARC-exo also reveals highly heterogeneous conformations of expansion segments in cells, challenging to characterize using conventional physical methods. Finally, we integrated SHARC-exo with two other methods, PARIS and CLIP, to conclusively determine a secondary structure model for the 7SK RNA and reveal a compact folding of the multiple helices. These results highlight significant advancements compared to previous methods for RNA 3D structure analysis.

Future improvements and extension of the SHARC-exo principle will further enhance its versatility and reliability and broaden its applications. First, despite our careful in vitro analysis of the SHARC reagents, the kinetics of the two-step acylation and hydrolysis reactions are difficult to characterize experimentally and theoretically because they are not necessarily decoupled or orthogonal. These reactions are likely different in vitro and in vivo, making it challenging to develop a simplified model to study them in detail. Nevertheless, better understanding of these reactions is important for further improvement of the crosslinking chemistry. The current RNase R trimming is not 100% efficient, likely due to the presence of SHARC monoadducts on RNA. Even though our computational identification of trimming stop sites uses the 3’ end medians, which is robust against outliers, further improvement of this experimental step will increase the resolution and accuracy. Most cellular RNAs are associated with proteins. Incorporating RNA-protein interactions and protein structure information will enable 3D modeling of RNP complexes in cells. Current acylation-based crosslinkers apply to all four nucleotides yet are limited to flexible ones. In some highly structured RNAs, the number of flexible and, therefore, cross-linkable nucleotides might be moderate (**Supplementary Fig. 18**). Critical spatially proximal nucleotides may be non-reactive, making it potentially challenging to capture such constraints. In the future, the development of chemical crosslinkers that react with other functional groups in RNA with reduced bias will further improve the efficiency, resolution, and dynamic range of the distance measurements. Current modeling methods that can use experimental constraints, such as Rosetta, are extremely computationally expensive. With the ability to measure spatial distances in high throughput, new computational tools are urgently needed further to exploit the rich structural information in the SHARC-exo data and enable more rapid 3D modeling for larger RNAs and deconvolution of structural ensembles on a transcriptome-wide scale. Targeted enrichment coupled with SHARC-exo can be applied to many low-abundance RNAs to study their structures ^24^. We anticipate that direct high throughput analysis of RNA 3D structures in vivo will reveal new principles of RNA structure formation and function. Given the critical roles of RNA in human genetic and infectious diseases, in vivo, 3D structural information is invaluable for developing RNA-based and RNA-targeted therapeutics.

## METHODS

Methods, including statements of data availability and any associated accession codes and references, are available in the online version of the paper.

## Acknowledgments

We thank members of the Lu and Velema labs for their help with this work. The Lu lab is supported by startup funds from the University of Southern California, the Pathway to Independence Award from NHGRI (R00HG009662), NIGMS (R35GM143068), USC Research Center for Liver Disease (P30DK48522), Illumina and USC Keck Genomics Platform (KGP) Core Lab Partnership Program, the Norris Comprehensive Cancer Center (P30CA014089) and USC Center for Advanced Research Computing. The Velema lab is supported by startup funds from Radboud University and an NWO ENW XS grant from the Dutch Research Council (OCENW.XS4.215). The Yesselman lab is supported by startup funds from the University of Nebraska-Lincoln, and Tobacco Settlement Fund (21-5734-0010)

## Author contributions

Z.L. and W.A.V. conceived the project. W.A.V. performed the SHARC synthesis and crosslink characterizations. R.V.D. and Z.L. developed the SHARC-exo strategy. K.L. and J.B. performed the cellular RNA structure experiments. Z.L., M.Z., K.L., J.B. and W.H.L performed the sequencing data analysis. R.V.D. and J.D.Y. performed Rosetta modeling. Z.L., W.A.V. and J.D.Y. wrote the manuscript with input from all coauthors.

## Additional information

Any supplementary information, chemical compound information, and source data are available in the paper’s online version. Reprints and permissions information is available online. Correspondence and requests for materials should be addressed to Z.L. and W.A.V.

