## Supplementary material for "Chemical Reversible Crosslinking Enables Measurement of RNA 3D Distances and Alternative Conformations in Cells": SI

**Synthesis of activated dicarboxylic acids.** 1,1'-Oxalyldiimidazole was purchased from Tokyo Chemical Industries. All other activated dicarboxylic acids were synthesized (Scheme S1). The dicarboxylic acid (0.20 mmol) was dissolved in 0.1 mL of DMSO. To this was added a solution of CDI (0.40 mmol) in DMSO (0.1 mL) and the resulting mixture was kept under nitrogen at room temperature for 1 hour. Heavy bubbling was observed in all cases, which stopped after ~10 minutes. The resulting 1.0 M solution of activated dicarboxylic acids was used immediately in crosslinking experiments, without further purification. Successful activation was confirmed for all compounds and full analysis ($^1$H NMR, $^{13}$C NMR and MS) was obtained. Note that imidazole is formed as a byproduct in the reaction and is present in all spectra.

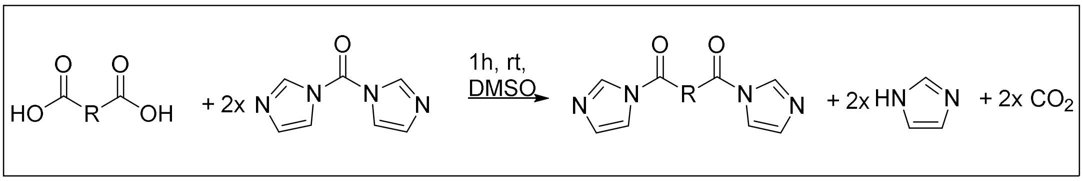


*Scheme S1*: Synthesis of crosslinkers from dicarboxylic acids and CDI.

Oxalic acid imidazolide: *1,2-di(1H-imidazol-1-yl)ethane-1,2-dione.* $^1$H NMR (400 MHz, DMSO) δ 8.23 (d, *J* = 1.1 Hz, 1H), 7.58 (t, *J* = 1.4 Hz, 1H), 7.05 – 7.00 (m, 1H). $^{13}$C NMR (101 MHz, DMSO) δ 162.39, 138.16, 130.47, 116.87. ESI-MS: [M+H] calculated: 191.1 ; Found: 191.8

Succinic acid imidazolide: *1,4-di(1H-imidazol-1-yl)butane-1,4-dione.* $^1$H NMR (400 MHz, DMSO) δ 8.52 (t, *J* = 1.1 Hz, 2H), 7.77 (t, *J* = 1.5 Hz, 2H), 7.12 (dd, *J* = 1.7, 0.8 Hz, 2H), 3.49 (s, 4H). $^{13}$C NMR (101 MHz, DMSO) δ 169.98, 137.61, 130.91, 116.99, 29.65. ESI-MS: [M+H] calculated: 219.1 ; Found: 219.2

Glutaric acid imidazolide: *1,5-di(1H-imidazol-1-yl)pentane-1,5-dione.* $^1$H NMR (400 MHz, DMSO) δ 8.42 (t, *J* = 1.1 Hz, 2H), 7.71 (t, *J* = 1.5 Hz, 2H), 7.08 (dd, *J* = 1.7, 0.8 Hz, 2H), 3.16 (t, *J* = 7.0 Hz, 4H), 2.06 (p, *J* = 7.0 Hz, 2H). $^{13}$C NMR (101 MHz, DMSO) δ 170.58, 137.39, 130.69, 116.91, 33.80, 18.27. ESI-MS: [M+H] calculated: 233.1 ; Found: 233.2

Diglycolic acid imidazolide: *2,2'-oxybis(1-(1H-imidazol-1-yl)ethan-1-one).* $^1$H NMR (400 MHz, DMSO) δ 8.41 (t, *J* = 1.1 Hz, 2H), 7.71 (t, *J* = 1.5 Hz, 2H), 7.10 (dd, *J* = 1.7, 0.8 Hz, 2H), 5.04 (s, 4H). $^{13}$C NMR (101 MHz, DMSO) δ 167.93, 137.10, 130.66, 116.55, 69.65. ESI-MS: [M+H] calculated: 235.1 ; Found: 235.1

Terephthalic acid imidazolide: *1,4-phenylenebis((1H-imidazol-1-yl)methanone).* $^1$H NMR (400 MHz, DMSO) δ 8.28 (t, *J* = 1.1 Hz, 2H), 8.02 (s, 4H), 7.76 (t, *J* = 1.5 Hz, 2H), 7.25 – 7.16 (m, 2H). $^{13}$C NMR (101 MHz, DMSO) δ 165.97, 139.25, 136.03, 131.20, 130.37, 118.73. ESI-MS: [M+H] calculated: 267.1 ; Found: 267.0

Isocinchomeronic acid imidazolide: *pyridine-2,5-diylbis((1H-imidazol-1-yl)methanone).* $^1$H NMR (400 MHz, DMSO) δ 9.16 (dd, *J* = 2.2, 0.8 Hz, 1H), 8.71 (t, *J* = 1.0 Hz, 1H), 8.54 (dd, *J* = 8.1, 2.2 Hz, 1H), 8.37 – 8.31 (m, 2H), 7.97 (dd, *J* = 1.7, 1.2 Hz, 1H), 7.81 (t, *J* = 1.5 Hz, 1H), 7.23 (dd, *J* = 1.7, 0.8 Hz, 1H), 7.19 (dd, *J* = 1.7, 0.8 Hz, 1H). $^{13}$C NMR (101 MHz, DMSO) δ 164.53, 163.36, 152.20, 149.66, 140.30, 139.77, 139.32, 131.74, 131.38, 130.69, 126.34, 124.68, 118.68, 118.61. ESI-MS: [M+H] calculated: 268.1 ; Found: 268.2

6,6'-binicotinic acid imidazolide: *[2,2'-bipyridine]-5,5'-diylbis((1H-imidazol-1-yl)methanone).* $^1$H NMR (400 MHz, DMSO) δ 9.16 (dd, *J* = 2.2, 0.8 Hz, 2H), 8.69 (dd, *J* = 8.2, 0.9 Hz, 2H), 8.46 (dd, *J* = 8.3, 2.3 Hz, 2H), 8.35 (d, *J* = 1.1 Hz, 2H), 7.81 (t, *J* = 1.5 Hz, 2H), 7.22 (d, *J* = 1.6 Hz, 2H). $^{13}$C NMR (101 MHz, DMSO) δ 167.89, 157.57, 150.81, 139.70, 139.09, 131.21, 129.46, 121.58, 118.77. ESI-MS: [M+H] calculated: 345.1 ; Found: 345.2

Dipicolinic acid imidazolide (DPI for short): *pyridine-2,6-diylbis((1H-imidazol-1-yl)methanone).* $^1$H NMR (400 MHz, dmso) δ 8.67 (dd, *J* = 1.3, 0.8 Hz, 2H), 8.51 – 8.37 (m, 3H), 7.94 (dd, *J* = 1.7, 1.2 Hz, 2H), 7.18 – 7.11 (m, 2H). $^{13}$C NMR (101 MHz, DMSO) δ 163.1, 148.8, 140.5, 140.1, 130.6, 130.2, 118.8. ESI-MS: [M+H] calculated: 268.1 ; Found: 268.3.

**In vitro crosslinking of model RNA.** The model RNA **1** was purchased from Integrated DNA Technologies. Nine µL of 10 µM RNA **1** in 0.06 M MOPS, pH 7.5; 0.1 M KCl; 2.5 mM $MgCl_2$, was heated to 95 °C for 2 min and then slowly cooled to room temperature. One µL of 1 M activated dicarboxylic acid stock solution in DMSO was added and the mixture was incubated for 4 hours at room temperature. Reactions were quenched by addition of 9 volumes of precipitation solution (0.33M NaOAc, pH 5.2, glycogen 0.2 mg/mL) and 30 volumes of absolute ethanol. RNA was precipitated for 1 hour at -20˚C, and then centrifuged (21000 RCF) for 40 min at 4˚C. The pellet was washed with 70% ethanol, air dried, and resuspended in 10 µL RNase-free water. Precipitated RNA was analyzed using 20% PAGE and imaged using Sybr Gold and a Bio-Rad Gel Documentation System and safeVIEW-MINI2 Imaging System. The distribution between unreacted RNA 1and crosslinked RNA was determined by quantifying the band intensity with ImageJ. All experiments were performed in triplicate.

**Reversal of in vitro crosslinked RNA.** Five µL of 10 µM crosslinked RNA in water was diluted with 45 µL 100 mM borate buffer pH 10.0 and incubated for 2 hours at 37 °C. Reactions were quenched by addition of 50 uL of precipitation solution (0.33M NaOAc, pH 5.2, glycogen 0.2 mg/mL) and 300 µL of absolute ethanol. RNA was precipitated for 1 hour at -20˚C, and then centrifuged (21000 RCF) for 40 min at 4˚C. The pellet was washed with 70% ethanol, air dried, and resuspended in 10 µL RNase-free water. RNA was analyzed using 20% PAGE and imaged using Sybr Gold and a Bio-Rad Gel Documentation System and safeVIEW-MINI2 Imaging

System. The distribution between unreacted RNA 1and crosslinked RNA was determined by quantifying the band intensity with ImageJ. All experiments were performed in triplicate.

**Cell culture.** HeLa and HEK293 cells were purchased from ATCC and maintained in Dulbecco's modified Eagle's medium (DMEM, Gibco) + 10% fetal bovine serum (FBS, Gibco) + Pen/Strep antibiotic, in 37°C incubator with 5% CO2. All cell culture were handled according to protocols approved by the University of Southern California.

**SHARC crosslinker preparation for crosslinking.** SHARC reagents were made by dissolving 1 part SHARC reactant in 200µl anhydrous DMSO (Sigma, 276855) and 2 parts CDI Sigma, 115533) in 250µl DMSO. Dissolved SHARC reactant was pipetted into the tube containing CDI. After briefly vortex and spinning down, a needle was inserted into the top of the 1.5mL centrifuge tube to allow $CO_2$ product to escape. Mixed solutions were left on the room temperature to react for 30-60 minutes before crosslinking.

**In vivo cross-linking.** Hela and HEK293 cells with 80% confluency in a 10 cm dish were washed twice with 1x PBS. Then cells were collected, resuspended in 1x PBS and transferred into a 1.5ml tube with final volume of 900µl. For each tube of cells, added 100µl of SHARC crosslinker to make the final concentration of 0, 5, 12.5 and 25mM. Cells were incubated in a rotator at room temperature for 30 minutes. After cross-linking, cross-linking solution was removed and cells were washed twice with 1x PBS.

**Extraction of cross-linked RNA (TNA method,** adapted from [1]**).** For each 10cm dish cells, added 100µl of 6M GuSCN (Sigma, 368975) and lysed cells with vigorous manual shaking for 1min. Then, cell lysate was added 12µl of 500mM EDTA (Invitrogen™, 15575020), 60µl of 10x PBS(Invitrogen™, AM9625), and water to final volume of 600µl. Each sample was passed through a 25G or 26G needle about 20 times to further break the insoluble material. Proteinase K (PK) (Thermo Scientific™, EO0492) was added to final concentration of 1mg/ml, and PK treatment was performed at 37ºC for 1 hour on a shaker at 600-900RPM. After PK digestion, 60µl of 3M sodium acetate (pH 5.3) (Invitrogen™, AM9740), 600µl of water-saturated phenol (pH 6.6) (Invitrogen™, AM9712), and 1 volume pure isopropanol were added to precipitate total nucleic acids by spinning at 15000 rpm for 20 minutes at 4ºC. After twice washing using 70% ethanol, total nucleic acids were resuspend in 300µl of nuclease-free water. For 100µg of TNA samples, 50units of TURBO™ DNase (Invitrogen™, AM2239) were added to remove DNA at 37ºC for 20 minutes. Then added 20µl of 3M sodium acetate, equal volume of water-saturated phenol, two volume of pure isopropanol to precipitate RNA sample by spinning 20mins at 12,000 x g at 4ºC.

**RNA fragmentation.** 10µg of cross-linked RNA was fragmented using 10µl of RNase III (NEB, M0245) with 50mM $MnCl_2$ and 1x supplied shortcut buffer at 37ºC for 5mins. After incubation, equal volume of phenol was immediately added to stop the reaction. Then one tenth volume of 3M sodium acetate (pH 5.3), 3µl of GlycoBlue (Invitrogen™, AM9516), three volume of pure ethanol were added to precipitate RNA. Fragmented RNA was resuspended in RNase-free water.

**DD2D purification of cross-linked RNA.** <u>First dimension gel.</u> Prepare 8% 1.5mm thick denatured first dimension gel using the UreaGel system (National Diagnostics, EC-833) with MOPS buffer (Fisher, BP2900500). Briefly, 3.2ml UreaGel concentrate, 5.8ml UreaGel diluent, 1ml 10x MOPS buffer, 80 µl 10% of APS and 4ul TEMED (Thermo Scientific™, 17919) were mixed to make 8% first dimension gel. Loading dsRNA ladder (NEB, N0363S) as molecular weight marker. Run the first dimension gel at 30W for 7~8 minutes in 1x MOPS buffer. After electrophoresis was finished, staining the gel with SYBR Gold (Invitrogen™, S11494) in 1x MOPS buffer and excising each lane between 50nt to topside from the first dimension gel. The second dimension gel can usually accommodate three gel splices. <u>Second dimension gel.</u> Prepare the 16% 1.5mm thick urea denatured second dimension gel using the UreaGel system with MOPS buffer. Briefly, 6.4ml UreaGel concentrate, 2.6ml UreaGel diluent, 1ml 10x MOPS buffer, 80 µl 10% of APS and 4ul TEMED were mixed to make 16% first dimension gel. Using prewarmed 1x MOPS buffer to fill the electrophoresis chamber to facilitate denaturation of the cross-linked RNA. Run the second dimension at 30W for 50 minutes to maintain high temperature and promote denaturation. Gel containing the cross-linked RNA above the diagonal from the 2D gel was excised and crushed for RNA extraction.

**RNase R treatment.** RNase R is a 3' -> 5' exonuclease that is capable of unwinding and digesting doublestranded RNA with a 3' overhang. Purified crosslinked RNAs from DD2D gel were treated with 20 units of RNase R (Biovision, M1228) in 1x RNase R digestion buffer with 5mM ATP at 45ºC for 2, 12 and 24 h, respectively. Control RNA was without RNase R treatment. After RNase R treatment, one tenth volume of 3M sodium acetate (pH 5.3), 3µl of GlycoBlue, three volume of pure ethanol were added to precipitate RNA.

**Proximity Ligation.** Purified RNA fragments were proximity ligated by T4 RNA Ligase1 (NEB, M0437M). Briefly, 2µl of 10x ligation buffer, 5µl of T4 RNA Ligase, 1µl of SuperaseIn (Invitrogen™, AM2696) and 1µl of 0.1mM ATP were added to 10µl of purified dsRNA fragments [2]. Ligation mixture was incubated at room temperature overnight. After ligation, the samples were boiled for 2 minutes to stop the reaction. After heat denaturation, samples were centrifuged to remove the precipitate and then precipitated by ethanol.

**Reverse crosslinking.** To proximity ligated RNA fragments, 5x decrosslinking buffer (500mM Boric acid, pH 11) was added, and nuclease free water was added to bring decrosslinking buffer to 1x. Samples were incubated for 2 hours at 45℃ to guarantee reversal (this is higher than the temperatures used in the in vitro experiments). After reverse crosslinking, RNA was purified with three volume of ethanol and 1µl of GlycoBlue.

**Adapter Ligation.** Reverse crosslinked RNAs were heated at 80ºC for 90s, then snapped cooling on ice. To each sample, 3µl of 10µM ddc adapter /5rApp/AGATCGGAAGAGCGGTTCAG/3ddC/, 1µl of T4 RNA ligase 1, 2µl of DMSO, 5µl of PEG8000, 1µl of 0.1M DTT, 1µl of SuperaseIn and 2µl of 10x T4 RNA ligase buffer were added to perform adapter libation at room temperature for 3 hours. After adapter ligation, following reagents were added to remove free adapters: 3µl of 10x RecJf buffer (NEBuffer 2, B7002S), 2µl of RecJf (NEB, M0264S), 1µl of 5'Deadenylase (NEB, M0331S), 1µl of SuperaseIn, Reaction was incubated at 37ºC for 1 hour. Then 20µl of water was added to each sample to make total volume of 50µl and Zymo RNA clean and Concentrator-5 (Zymo Reasearch, R1013) was used to purify RNA.

**Reverse Transcription.** SuperScript IV (SSIV) (Invitrogen™, 18090010) was used to performing reverse transcription. The reaction buffer was optimized $Mn^{2+}$ buffer (1x): 50mM Tris-HCl (PH 8.3), 75mM $CH_3COOK$ and 1.5mM $MnCl_2$. Briefly, 1pmol of barcoded RT primer and 1µl of 10 mM dNTP were added to RNA samples and heated at 65ºC for 5 minutes in a PCR block, chill the samples one ice rapidly. Then 4µl of 5x $Mn^{2+}$ buffer, 2µl of 0.1 M DTT, 1µl of SuperaseIn and 1µl of SSIV were added to each sample. Mixed sample was incubated at 25ºC for 15 minutes, 42ºC for 10 hours, 80ºC for 10 minutes; hold at 10ºC. After reverse transcription, 1µl RNase H and RNase A/T1 mix were added and incubated at 37ºC for 30 minutes at 1000 rpm in a thermomixer to remove RNA. Synthesized cDNA were purified using Zymo DNA clean and Concentrator-5.

**cDNA circularization and library generation.** 1µl of CircLigase™ II ssDNA Ligase (Lucigen, CL9021K), 1µl of 50mM $MnCl_2$ and 10x CircLigaseII™ buffer were added to cDNA sample and performed circularization at 60ºC for 100 minutes. 80ºC treatment for 10 minutes was followed to stop the reaction. The circularised cDNA products were directly used to library PCR. Library PCR preparation was done as described in ref. [3]. PCR products were run on 6% native TBE gel. Gel containing DNA products from 175bp and topside (corresponding to > 40bp insert) was excised and crushed for DNA extraction.

**In vitro SHARC-exo analysis of the P4-P6 RNA.** The P4-P6 (PDB: 1HR2) DNA with T7 promoter (TAATACGACTCACTATAG) was purchased from twist bioscience. After PCR amplification, the DNA was cleaned up using the Qiagen PCR Purification Kit and purified using a 8% native polyacrylamide gel. The P4-P6 (1HR2) RNA was transcribed using the MEGAscript T7 Transcription Kit from Thermo Fisher (AM1334) from 136ng of DNA template, and purified on denatured polyacrylamide gels. 10ug of P4-P6 RNA, 10uL of refolding buffer, and water was added to a final volume of 44uL per sample. The RNA was then denatured by incubating at 90C° for 5 minutes followed by snap cooling on ice. 1uL of 500mM $MgCl_2$ was then added to each sample while cold and then mixed. Samples were then allowed to come to room temperature over several minutes to refold. After refolding, either 5uL of DMSO for controls or 5uL 50mM DPI was added to each sample. Samples were then incubated at room temperature for 30 minutes. After incubation, samples were purified using ethanol precipitation. The crosslinked RNA was then converted into cDNA libraries as described above. In particular, we divided the crosslinked RNA fragments from the DD2D gels into 2 fractions, where one was treated with RNase R at 37C° 2 hours, while the other was not treated. The cDNA library was sequenced on a MiSeq machine.

**SHARC-Seq analysis**
Mapping. The 3'end adapters of sequencing data were removed using Trimmomatic. PCR duplicates were removed using readCollapse script from the icSHAPE pipeline. After removing 5' header, reads were mapped to manually curated hg38 genome using STAR program[4]. The parameters used are as follows: STAR --runThreadN 8 --runMode alignReads --genomeDir OuputPath --readFilesIn SampleFastq --outFileNamePrefix Outprefix --genomeLoad NoSharedMemory outReadsUnmapped Fastx --outFilterMultimapNmax 10 --outFilterScoreMinOverLread 0 --outSAMattributes All --outSAMtype BAM Unsorted SortedByCoordinate --alignIntronMin 1 --scoreGap 0 --scoreGapNoncan 0 --scoreGapGCAG 0 --scoreGapATAC 0 --scoreGenomicLengthLog2scale -1 --chimOutType WithinBAM HardClip --chimSegmentMin 5 --chimJunctionOverhangMin 5 --chimScoreJunctionNonGTAG 0 --chimScoreDropMax 80 --chimNonchimScoreDropMin 20.

Classify alignments. The primary mapping alignments were extracted from SampleAligned.sortedByCoord.out.bam, and classified into six different types using gaptypes.py (https://github.com/zhipenglu/CRSSANT) [5]. cont.sam, continuous alignments; gap1.sam, non-continuous alignments with one gap; gapm.sam, non-continuous alignments with more than one gaps; trans.sam, non-continuous alignments with the two arms on different strands or chromosomes; homo.sam, non-continuous alignments with the two arms overlapping each other; bad.sam, non-continuous alignments with complex combinations of indels and gaps. Gap1. and gapm alignments containing splicing junctions and short 1-2 nt gaps were filtered out using gapfilter.py (https://github.com/zhipenglu/CRSSANT). Then filtered gap1.sam, filtered gapm.sam and trans.sam were used to analyze RNA structures and interactions.

Cluster alignments to groups. Filtering alignments were assembled to DGs and NGs using the crssant.py script (https://github.com/zhipenglu/CRSSANT). After DG clustering, crssant.py verifies that the DGs do not contain any non-overlapping reads, i.e. any reads where the start position of its left arm is greater than or equal to the stop position of the right arm of any other read in the DG. If the DGs do not contain any non-overlapping reads, then the following output files ending in the following are written:
Sample.sam: SAM file containing alignments that were successfully assigned to DGs, plus DG and NG annotations;
dg.bedpe: bedpe file listing all duplex groups.

**Visualization of SHARC-seq data in Intergrative Genomic Viewer.** Assembled alignments with DGs tag were displayed using intergrative Genomic Viewer (IGV)[6] visualization tool (V.2.8.13). The bed output file (from crssant.py script) can be visualized in IGV, where the two arms of each DG can be visualized as two 'exons', or as an arc the connects far ends of the DG.

**Structure analysis of rRNAs.** To analyze the RNase R trimming efficiency (e.g. Fig. 2f), we examined gapped alignments against the ribosome cryo-EM structure. For each alignment, we calculated the minimal physical distance between the two arms in the ribosome. Then the nucleotides involved in the minimal distances were recorded (counting from the 3' end of each arm). In a hypothetical example, we found that the minimal distance between the two arms in one read was between the $10^{th}$ nt from the right arm and $15^{th}$ nt from the left arm (both counting from the 3' ends). Then this tuple (10, 15) is considered one point on the heatmap (Fig. 2f and Supplementary Fig. 5b). After all the minimal distance nucleotides are calculated, their frequencies are plotted in the heatmap in square root scale.

The SHARC-seq reads aligned to 45S pre-rRNA (NR_046235.3) were collected and used to construct the interaction matrix. To build the physical interaction map of 28S rRNA and 18S rRNA, the cryo-EM model of the 28S rRNA and 18S rRNA was downloaded from RCSB Protein Data Bank (PDB) (ID: 4V6X). Watson-Crick and non-Watson-Crick base pairs were analyzed using the DSSR software [7]. The 3D structures of ribosome were visualized by the PyMOL system (Educational version, https://pymol.org/2/). Spatial distances in the cryo-EM model were extracted directly for use. The resolution of the human ribosome cryo-EM model is highly variable across the entire complex (PDB: 4V6X, Supplementary Fig. 2c from [8]). Although the average resolution is 5.4Å, the lowest goes to 21Å. The ribosome structure analysis and conclusions are based on longer distance intervals, e.g. 0-20, 20-40Å, etc. The modeling runs also used 0-20 and >20Å as the intervals for penalty calculations. In addition, the low-resolution regions are confined to the expansion segments, and do not affect the analysis of the stable core regions (as we showed in the B-factor plot, Supplemental Fig. 11d-e). In our analysis of the expansion segments, the distances that we focused on are much longer than 20A (Fig. 5e-f, Supplementary Fig. 11h-i). Therefore, the limited resolution of the ribosome model does not affect the analysis.

**Structure analysis of representative RNAs.** In order to accurately and easily analyze SHARC-seq data, pseudogenes and multicopy genes from gencode, refGene and Dfam were masked from hg38 genome. And then single copy of them was added back as a separated “chromosome”. For example, multicopy of snRNAs were masked from the basic hg38 assembly genome, and 9 snRNAs (U1, U2, U4, U5, U6, U11, U12, U4atac and U6atac) were concatenated into one reference, separated by 100nt “N”s, was added back. The curated hg38 genome contained 25 reference sequences, or “chromosomes”, masked the multicopy genes and added back single copies. This reference is best suited for the PARIS analysis. SHARC-seq reads were mapped to representative RNAs were collected and used for IGV visualization.

**Cross-linking distance analysis.** The ribose 2’OH in every flexible nucleotide (single stranded or icSHAPE activated) was used to calculate the cross-linking distance. The minimum distance between two arms’ flexible nucleotide to used to analyze the mininum distance distribution. The distance between No.3 to No.7 flexible nucleotides from 3’end of each arm was used for 3-7 distance distribution analysis.

**rRNA dynamic structure analysis.** The core and expansion segment boundaries of rRNA were derived from Chandramouli et al. (2008) [9] and Wakeman and Maden (1989) [10]. The SHARC-seq reads with ⩾40Å between two arms were collected and separated to core and expansion alignments. The dynamic reads were selected based on the rules that one arm mapped to the same region of rRNA, other arm mapped to different regions. The selected dynamic alignments were load to IGV for visualization.

**Computational Modeling of h22-24 region in the 18S.** Rosetta software (version 61146) was used to model RNAs for this study [11,12]. Helices of secondary structure regions were pre-built with the example command below to save computational expense: rna_helix.py -seq cag cug -resum 5-7 27-29 -o example_helix_1.pdb. The 920-1080 nucleotide region of the human 18S RNA was modeled with and without SHARC determined constraints. For the model set without SHARC constraints no cst file or flag was used. For the model set with SHARC constraints, the following linear energy function example command was used to assign constraints for 2’OH atoms that participated in the crosslinking reaction so that between 0A-20A there is no energy penalty and to apply a linearly scaling energy penalty if the atoms are >20A apart.

AtomPair O2' 63 O2' 117 LINEAR_PENALTY 10.0 0 10 1.0. Here 10.0 is the ideal distance between the atoms in Å, 0 is the energy penalty assigned to the range, 10 is the tolerance for the energy trough and 1.0 is the slope constraint.

Models were built with the command shown below using fasta, secondary structure and constraint files (for modeling set containing the SHARC data, otherwise no constraint file). For the native file used to get an rms for these files, the 920-1080 region of the 18S RNA was cut out using pymol and renumbered using renumber_pdb_in_place.py.

rna_denovo.static.linuxgccrelease -nstruct 1000 -fasta ../18s_920_1080.fasta -s ../18s_920_1080_helix_1.pdb ../18s_920_1080_helix_2.pdb-secstruct_file ../18s_920_1080.secstruct -cst_file ../18s_920_1080.cst -native ../18s_920_1080_renumbered.pdb -minimize_rna true -out:file:silent 18s_920_1080_tert.out

After modeling runs were finished, models were extracted using easy_cat.py. Example: easy_cat.py directory. To extract the top 1% scoring models from the bulk of the models for each run condition the following command was used (in this example 200 models are extracted): silent_file_sort_and_select.py [example_file.out] -select 1-200 -o [example_file_cluster.out]. The lowest 1% energy models were then clustered from each run condition to inspect the different pose topologies that existed within the lowest energy scoring models. Clustering was done with the command shown below (in this example 10 clusters are made using a cluster radius of 5Å: rna_cluster.static.linuxgccrelease -in:file:silent [example_file.out] -out:nstruct 10 -cluster:radius 5 -out:file:silent [example_file_cluster.out]. Clusters were extracted with the following command. The -no_replace_names flag here is used to prevent clusters from being renamed: python extract_lowscore_decoys.py [example_cluster_file.out] -no_replace_names

**Computational 3D modeling of the P4-P6 RNA.** The P4-P6 secondary structure was determined from PDB:1HR2 and is as follows: ....((((.(....((((((....(((..(((((((..(((((((((....)))))))))................)))....).))).)))...))))))...).))))((...((((...((((((((.....))))))))..))))...))..
Models were generated using the following sample command line: rna_denovo.static.linuxgccrelease -nstruct 1000 -fasta ../1hr2.fasta -secstruct_file ../1hr2.secstruct -s ../1hr2_helix* -cst_file ../1hr2_trim_tert.cst -native ../1hr2_chain_a_native.pdb -out:file:silent 1hr2_20A.out -minimize_rna true. Models generated without constraints had the –cst_file flag and cst file omitted from the command. The top 5 DGs by number of reads in SHARC-exo data, each with >3% of total reads, were used to constrain modeling with the equivalent DG being used for SHARC-constrained models. Linear atom pair constraint was set so that distances within 20Å carried no penalty and distances greater then 20Å were penalized with a slope of 1. RMSD values for models against the 1HR2 crystal structure was calculated. The top 100 scoring models from each group were clustered into 5 groups with a cluster radius of 5Å. Wilcox Ranked Sum Tests were performed between each two groups of top 100 models with the following R command wilcox.test(rmsd ~ group, data = XXX, exact = FALSE, alternative = 'greater').

**Analysis of hub1-hub2 alternative conformations**. First, the minimal 28S segment that contained both regions are limited to residues 3935 – 4175. Secondary structure was determined by running x3dna on the extracted segment [7]: x3dna-dssr -i=input.pdb -o=dssr.out. As the structure did not contain bases for all models so we went by hand to assign addition bases pairs base on geometry. Secondary structure is as follows: ((..(((((((.((((.((((..........(((((...((((.........(.((......................)).)................)))).............))))).)))).)).)).)))))))...(((((.....((((..((.((((((....))))))))).............
.((((((((((.....))))))))))...))))..)))))).....))

We generated 15317 models using FARFAR by command: rna_denovo.default.macosclangrelease -s stem_1.pdb stem_2.pdb helix_1.pdb helix_2.pdb -nstruct 1000 -fasta test.fasta -secstruct_file test.secstruct -minimize_rna false -cst_file test.cst, where the pdbs contained the original static structures of helices and stems not included in the contact. Linear atom pair constraint was set such that distances within 20 Å carries no penalty, while distance above 20 Å is penalized at a slope of 1. For each of the models we checked for steric clashes of the rest of the RNA and local proteins by comparing the distance between each phosphate of each modeled RNA nucleotide to the phosphate of each remaining RNA and the c-alpha of atom of each amino acid. A clash was defined as a distance of less than 5 angstroms.

**Analysis of 7SK structures using PARIS.** PARIS data from human and mouse cells were used to generate DGs for 7SK [13]. To analyze the secondary structures of 7SK, we clustered HEK293 PARIS non-continuous alignments on 7SK using CRSSANT [3].

**Analysis of 7SK structures using LARP7 eCLIP.** LARP7 eCLIP data in HepG2 and K562 cells were downloaded from ENCODE [14] and analyzed as follows. First reads mapped to 7SK were extracted from the mapped bam files (chr6:52995620-52995951 in hg38 coordinates). Reads with CIGAR gap flags D and N are extracted. All reads with D flags are converted to N for consistency. Then all reads with "N" were divided to three groups based on read start using the script readspan7SK.py and short-span reads were used to construct local structures.

**Data availability.** The raw and processed SHARC sequencing data was deposited to NCBI GEO with accession number GSE167812 (https://www.ncbi.nlm.nih.gov/geo/query/acc.cgi?acc=GSE167812) with the accession code ‘ezqnuoagzpedlah’.

**Code availability.** Custom codes used for data analysis in this paper can be found at https://github.com/zhipenglu/CRSSANT and https://github.com/minjiezhang-usc/SHARC-seq.

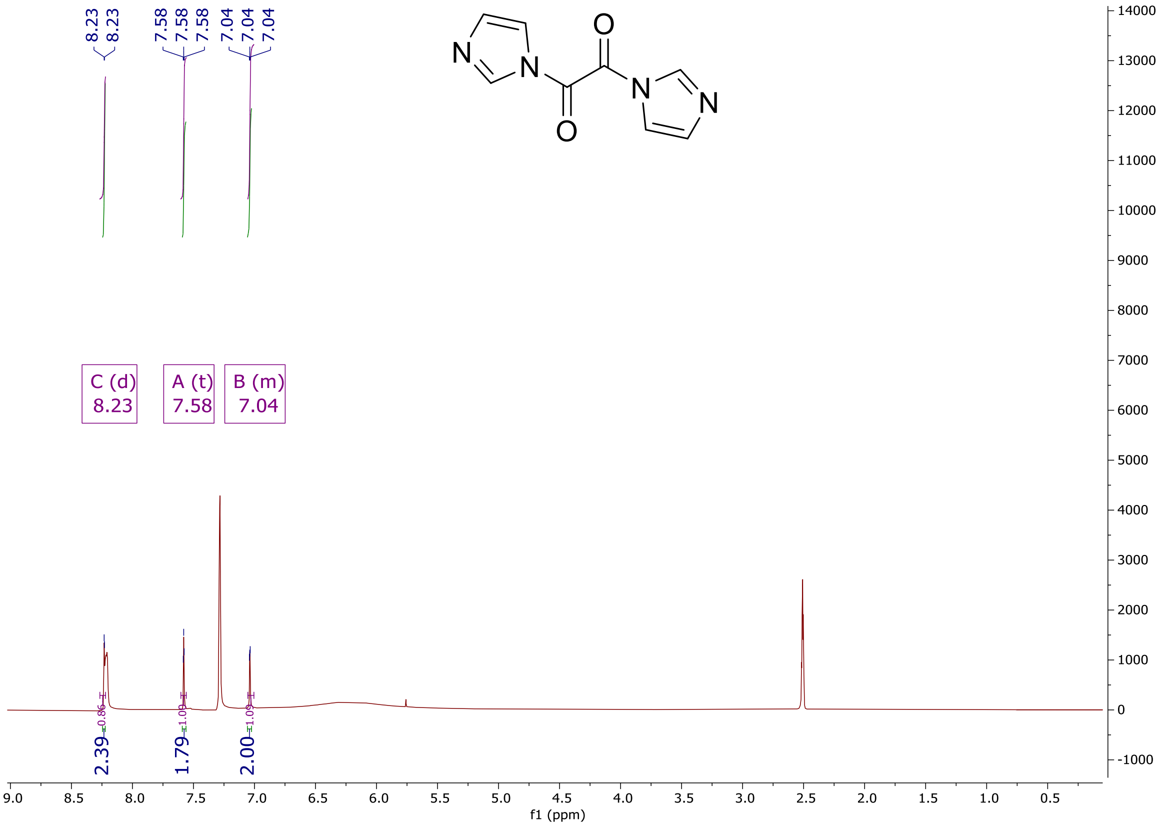


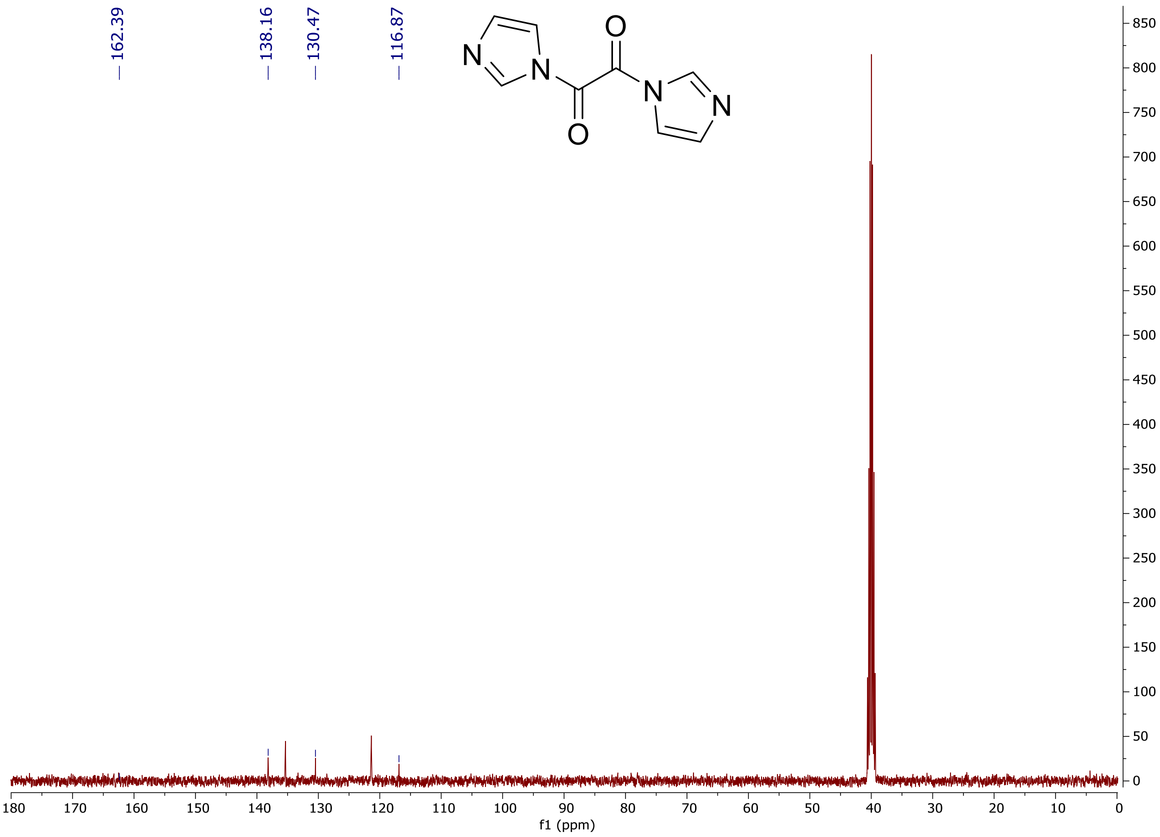


**Supplementary Figure 1. Mass spectrometry and NMR characterization of SHARC reagents.**

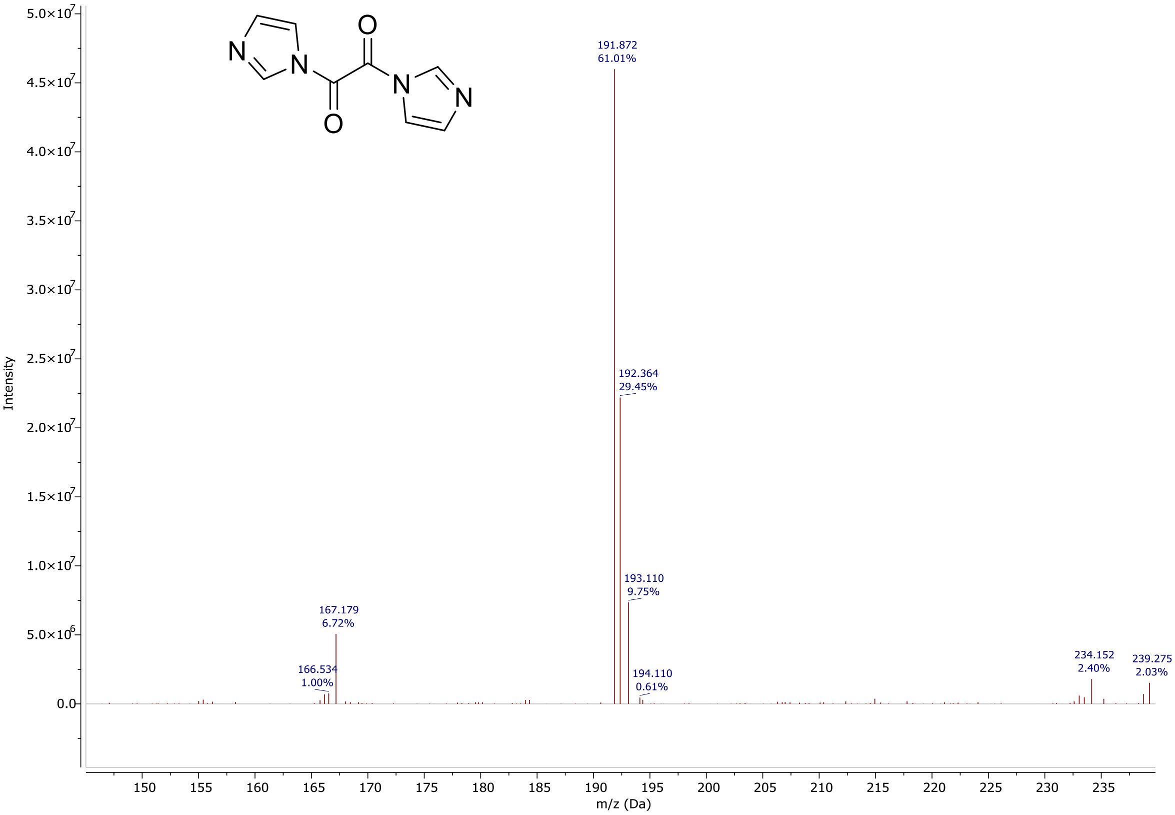

191.872
61.01%
192.364
29.45%
193.110
9.75%
194.110
0.61%
167.179
6.72%
166.534
1.00%
234.152
2.40%
239.275
2.03%
Intensity
m/z (Da)


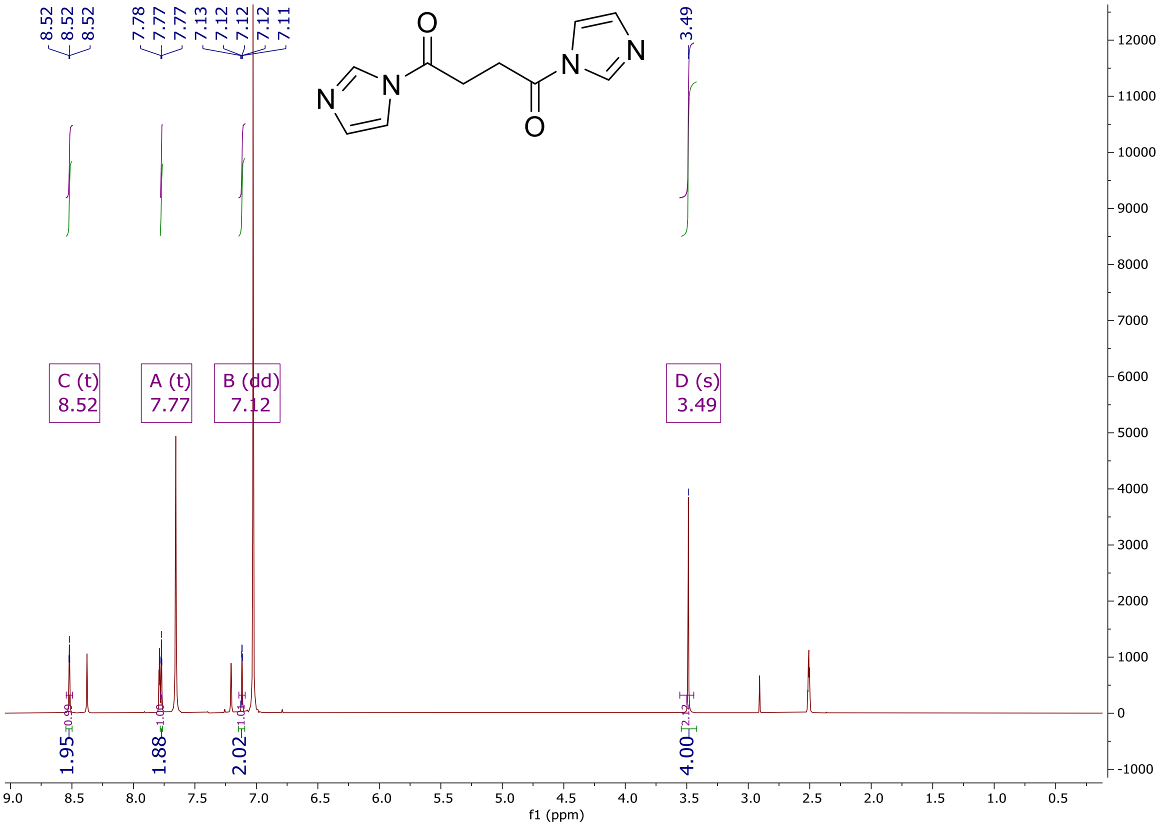

8.52
8.52
8.52
7.78
7.77
7.77
7.13
7.12
7.12
7.12
7.11
3.49
C (t)
8.52
A (t)
7.77
B (dd)
7.12
D (s)
3.49
1.95
1.88
2.02
4.00
f1 (ppm)

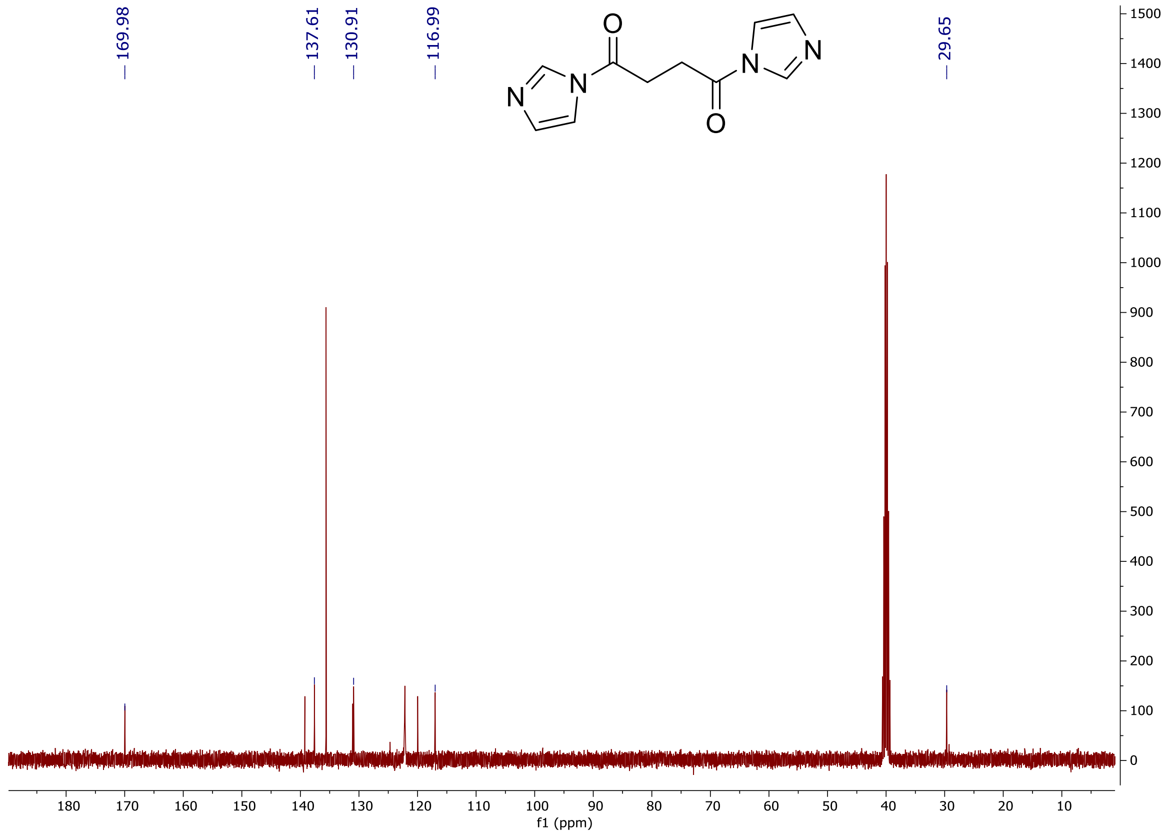
169.98
137.61
130.91
116.99
29.65
f1 (ppm)

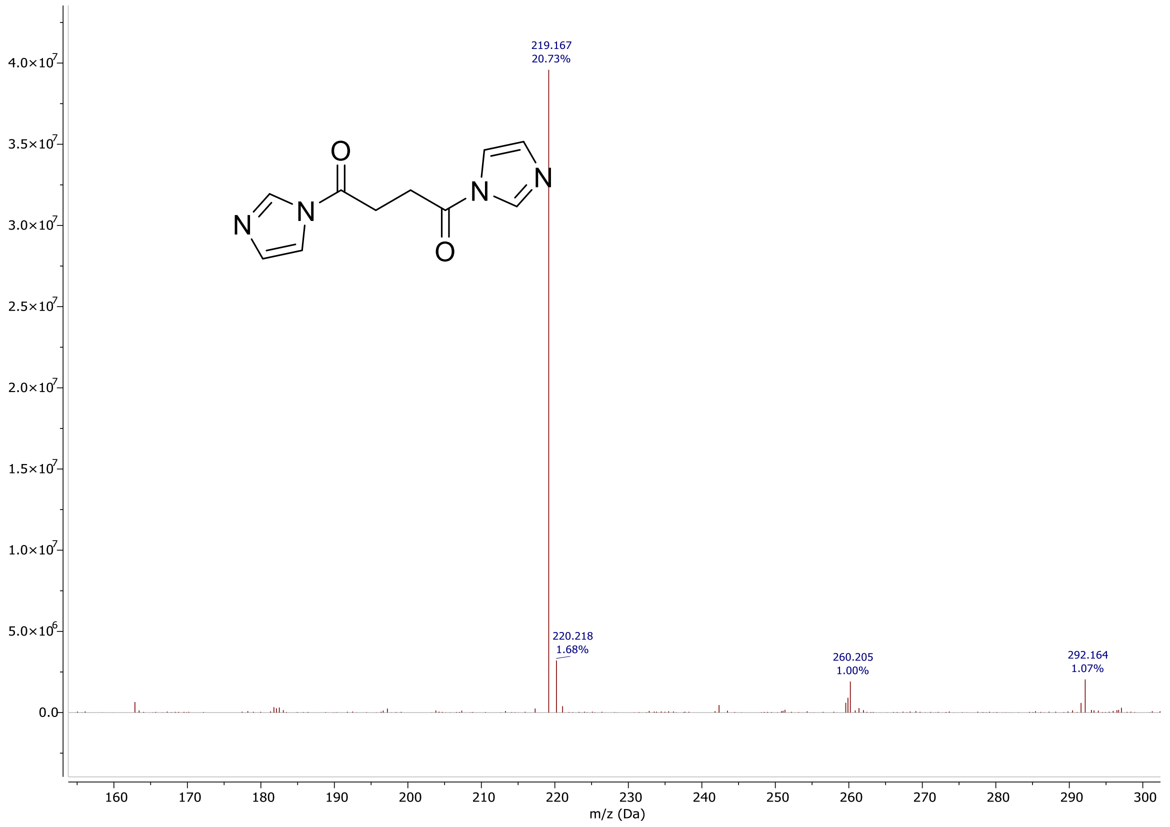
219.167
20.73%
220.218
1.68%
260.205
1.00%
292.164
1.07%
m/z (Da)

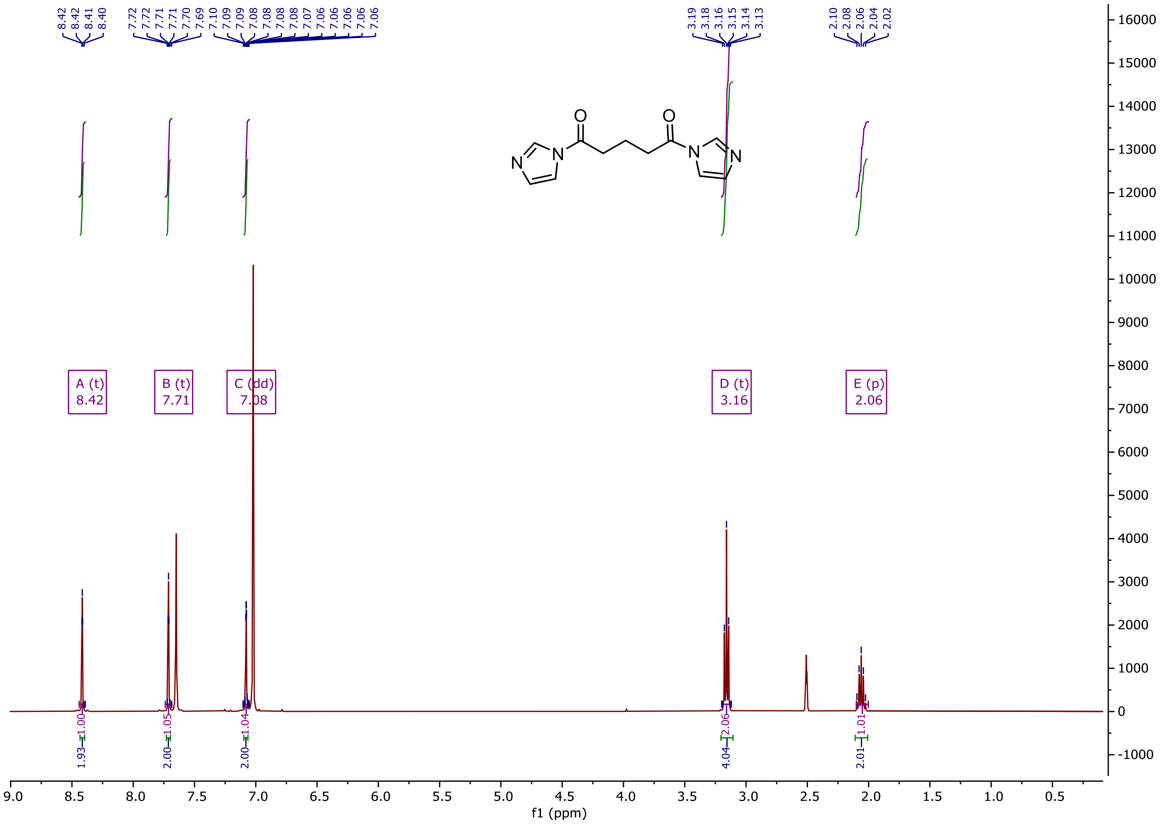

A (t)
8.42
B (t)
7.71
C (dd)
7.08
D (t)
3.16
E (p)
2.06
f1 (ppm)


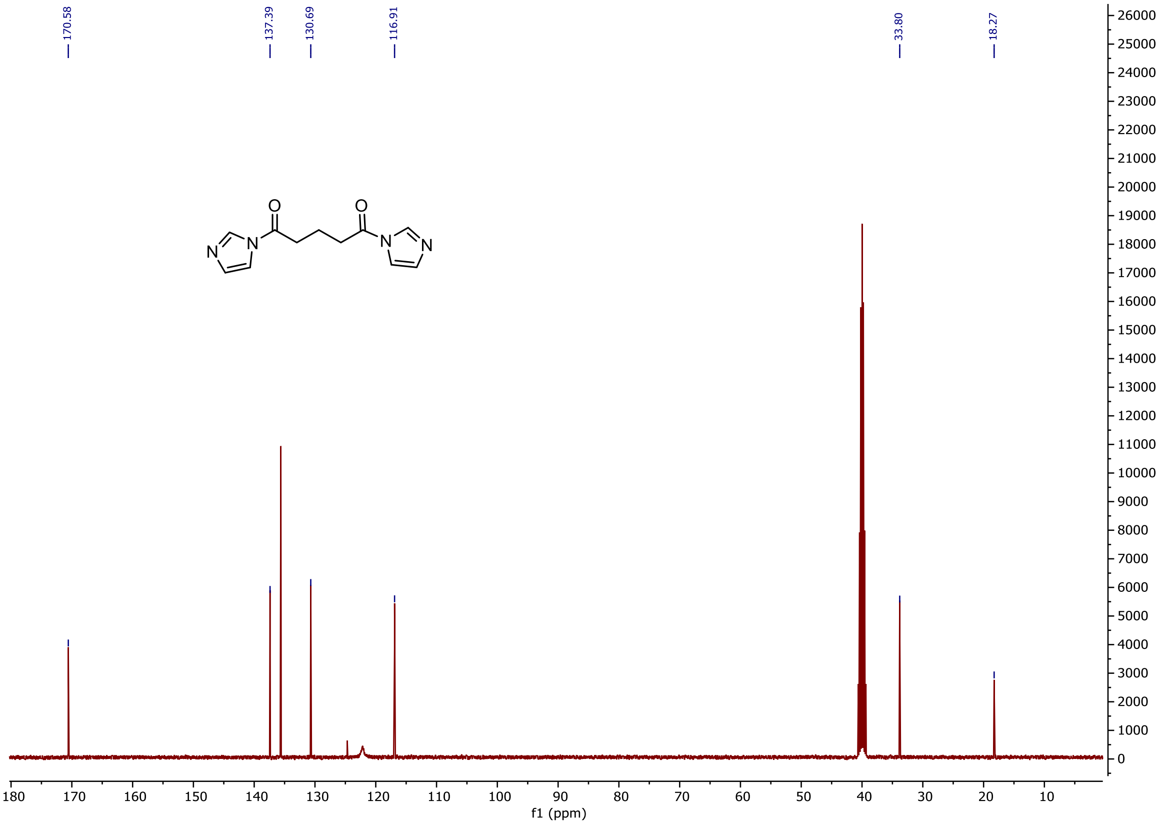

170.58
137.39
130.69
116.91
33.80
18.27
f1 (ppm)

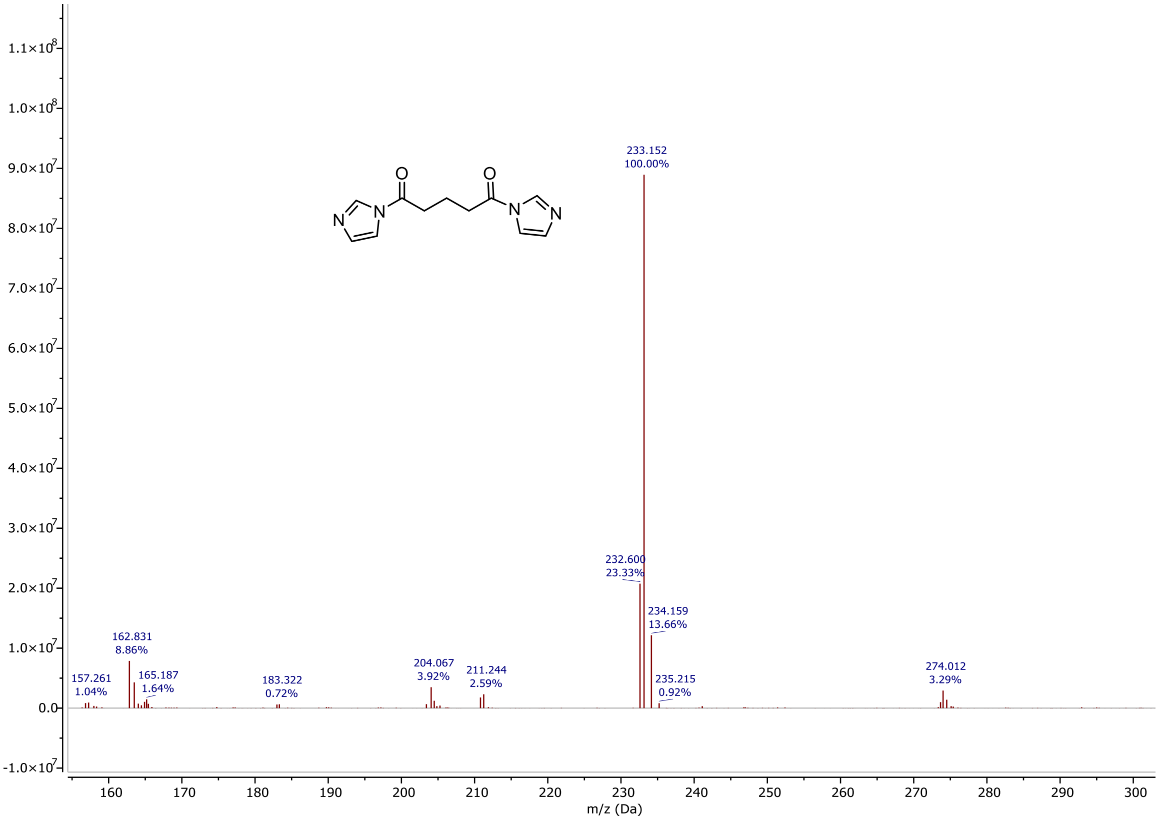
233.152
100.00%
232.600
23.33%
234.159
13.66%
162.831
8.86%
157.261
1.04%
165.187
1.64%
183.322
0.72%
204.067
3.92%
211.244
2.59%
235.215
0.92%
274.012
3.29%
m/z (Da)

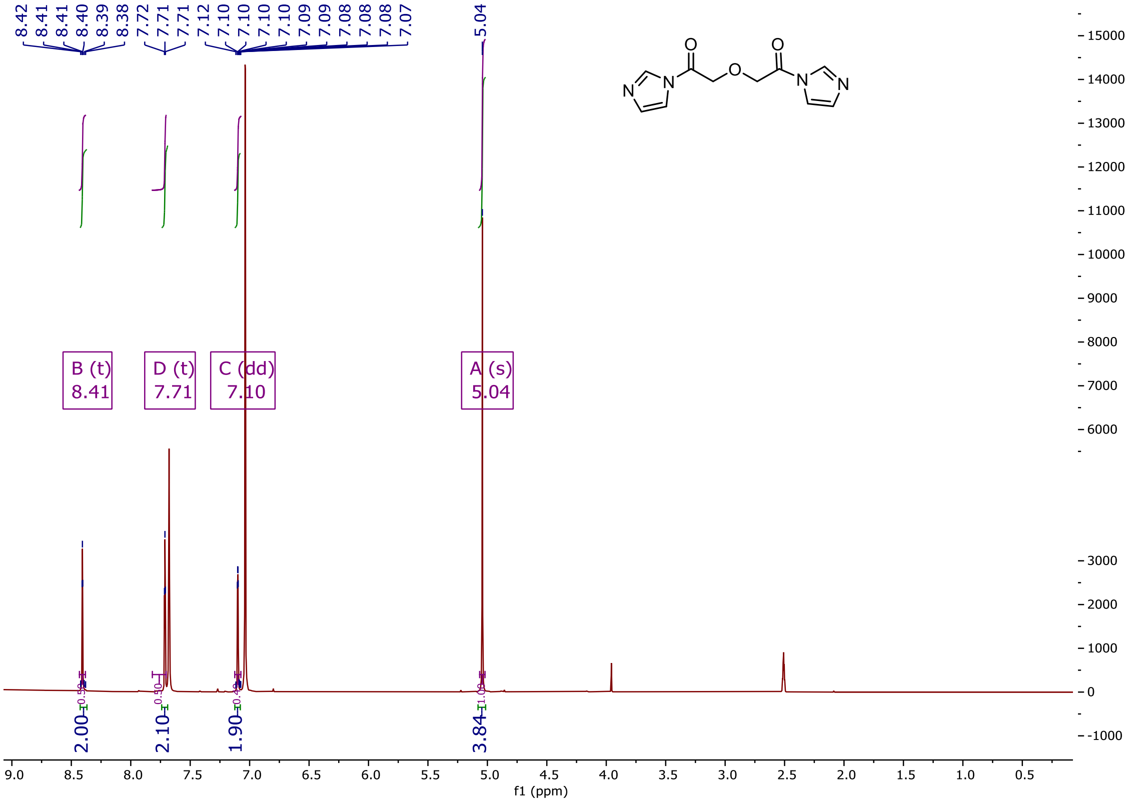
8.42
8.41
8.41
8.40
8.39
8.38
7.72
7.71
7.71
7.12
7.10
7.10
7.10
7.10
7.09
7.09
7.08
7.08
7.07
5.04
B (t)
8.41
D (t)
7.71
C (dd)
7.10
A (s)
5.04
2.00
2.10
1.90
3.84
f1 (ppm)

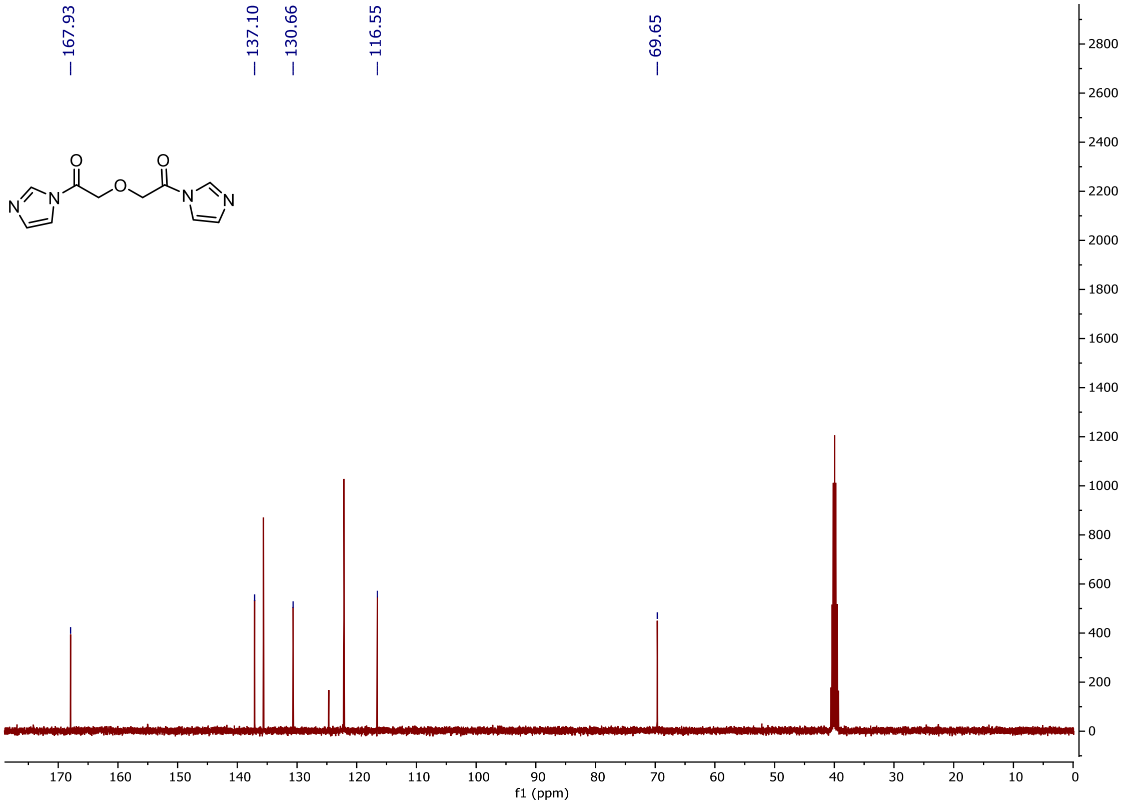

167.93
137.10
130.66
116.55
69.65
O
O
N
N
O
N
N
f1 (ppm)


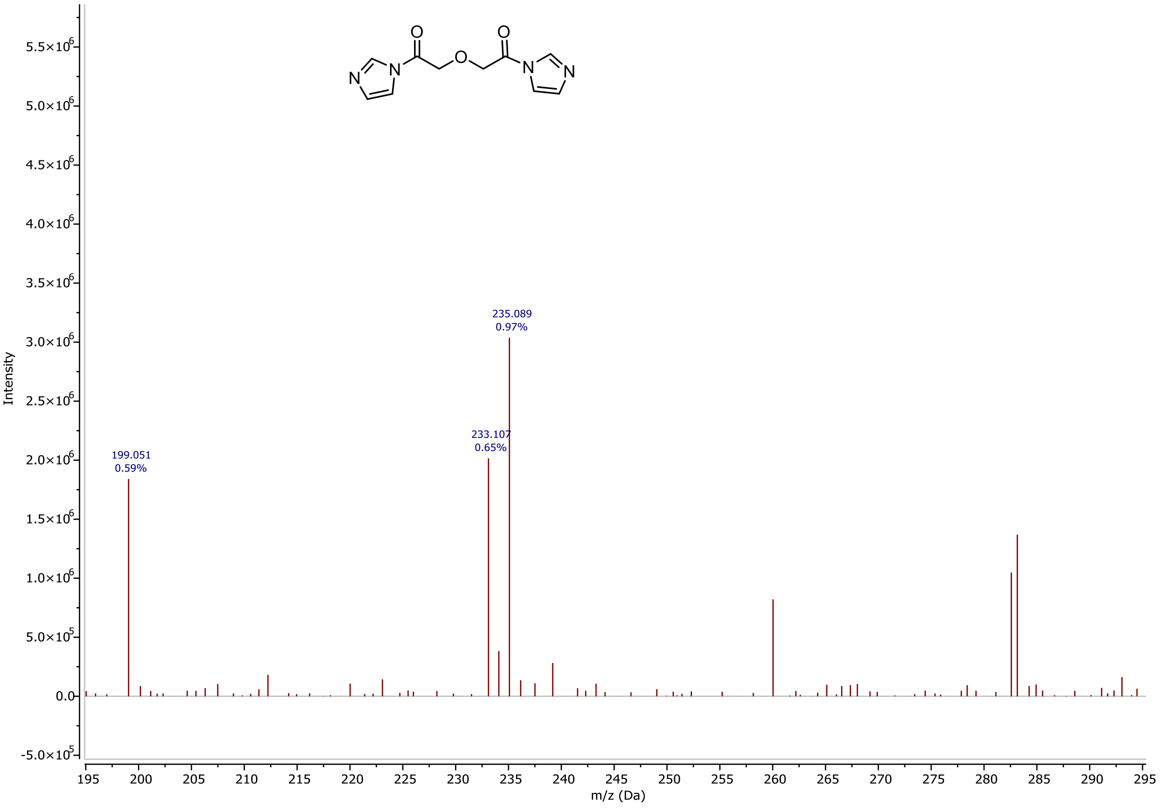

O
O
N
N
O
N
N
235.089
0.97%
233.107
0.65%
199.051
0.59%
Intensity
m/z (Da)

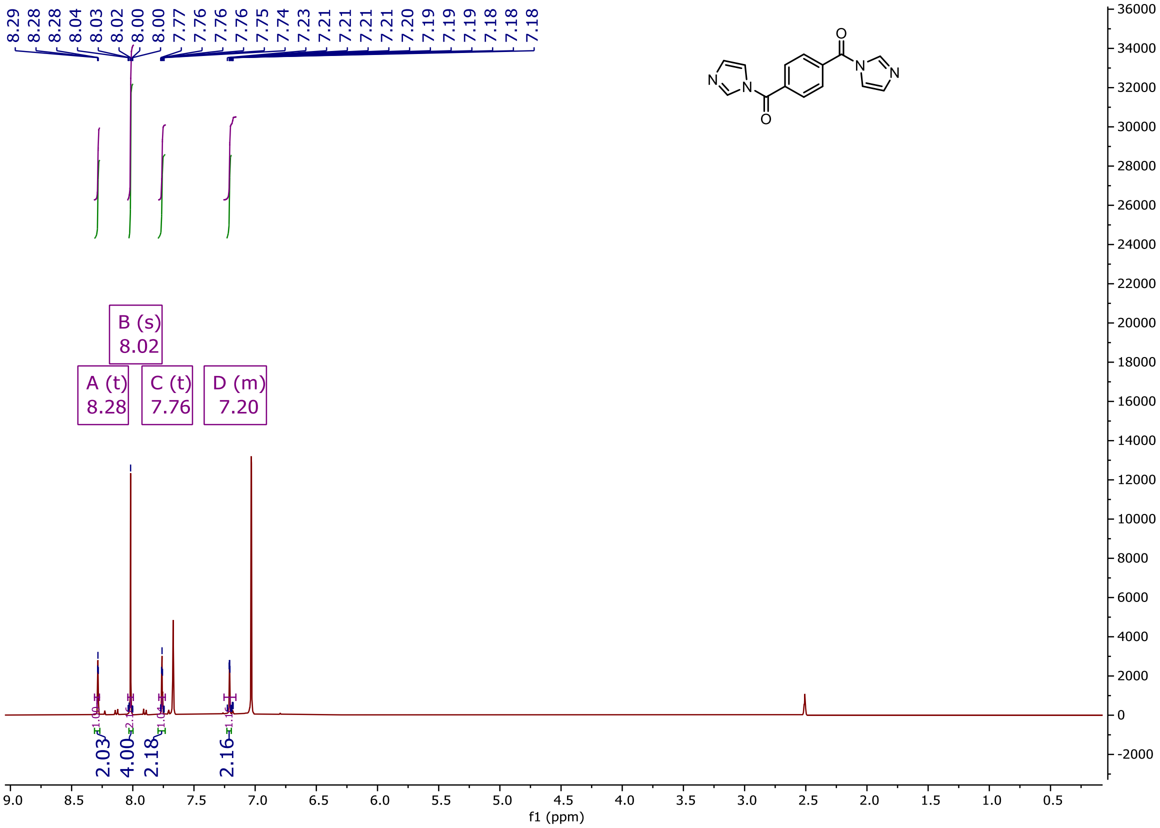

8.29
8.28
8.28
8.04
8.03
8.02
8.00
8.00
7.77
7.76
7.76
7.76
7.75
7.74
7.23
7.21
7.21
7.21
7.21
7.20
7.19
7.19
7.19
7.18
7.18
7.18
N
N
O
O
N
N
B (s)
8.02
A (t)
8.28
C (t)
7.76
D (m)
7.20
2.03
4.00
2.18
2.16
f1 (ppm)


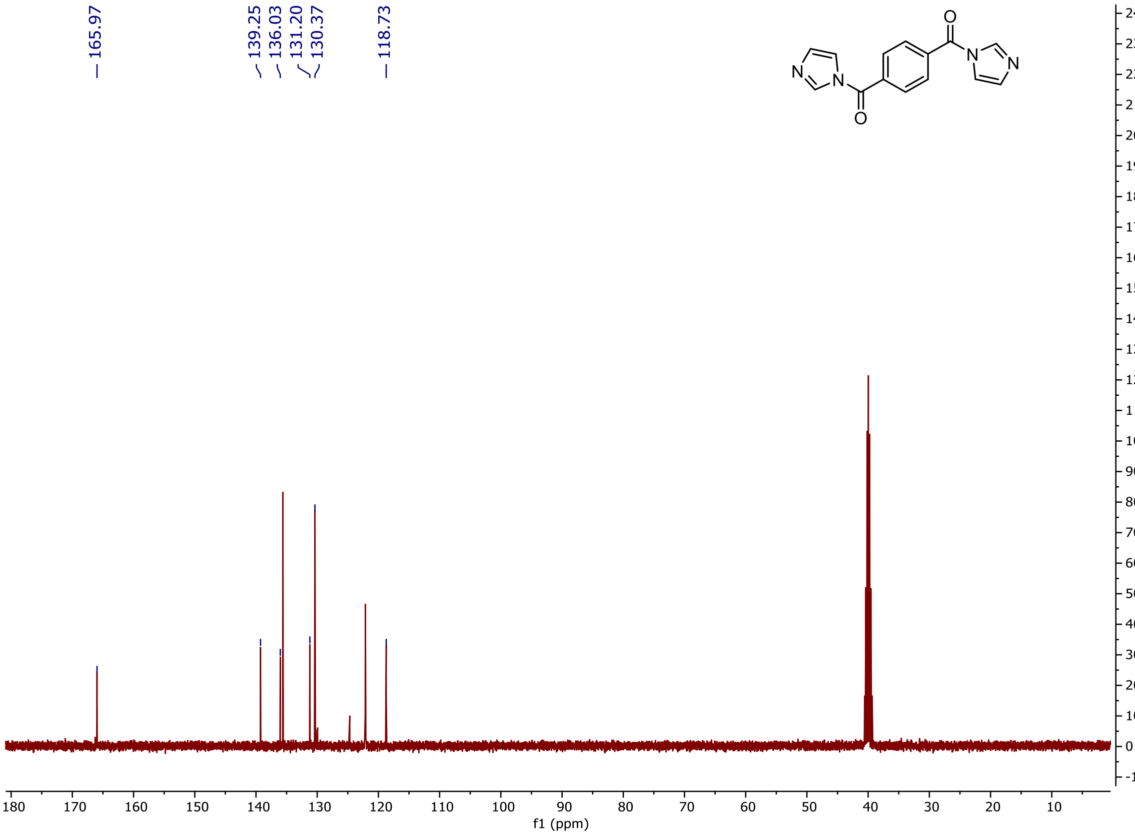

165.97
139.25
136.03
131.20
130.37
118.73
N
N
O
O
N
N
f1 (ppm)

! !

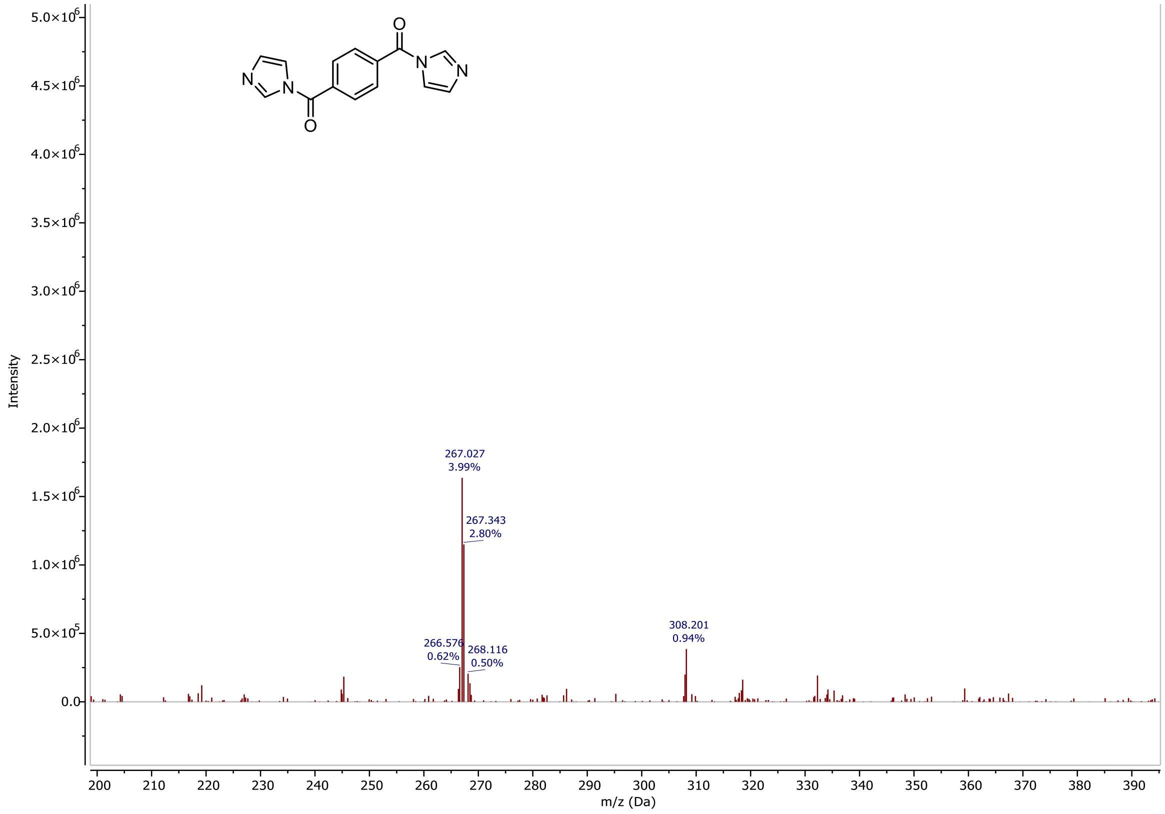

Intensity
m/z (Da)
267.027
3.99%
267.343
2.80%
266.576
0.62%
268.116
0.50%
308.201
0.94%


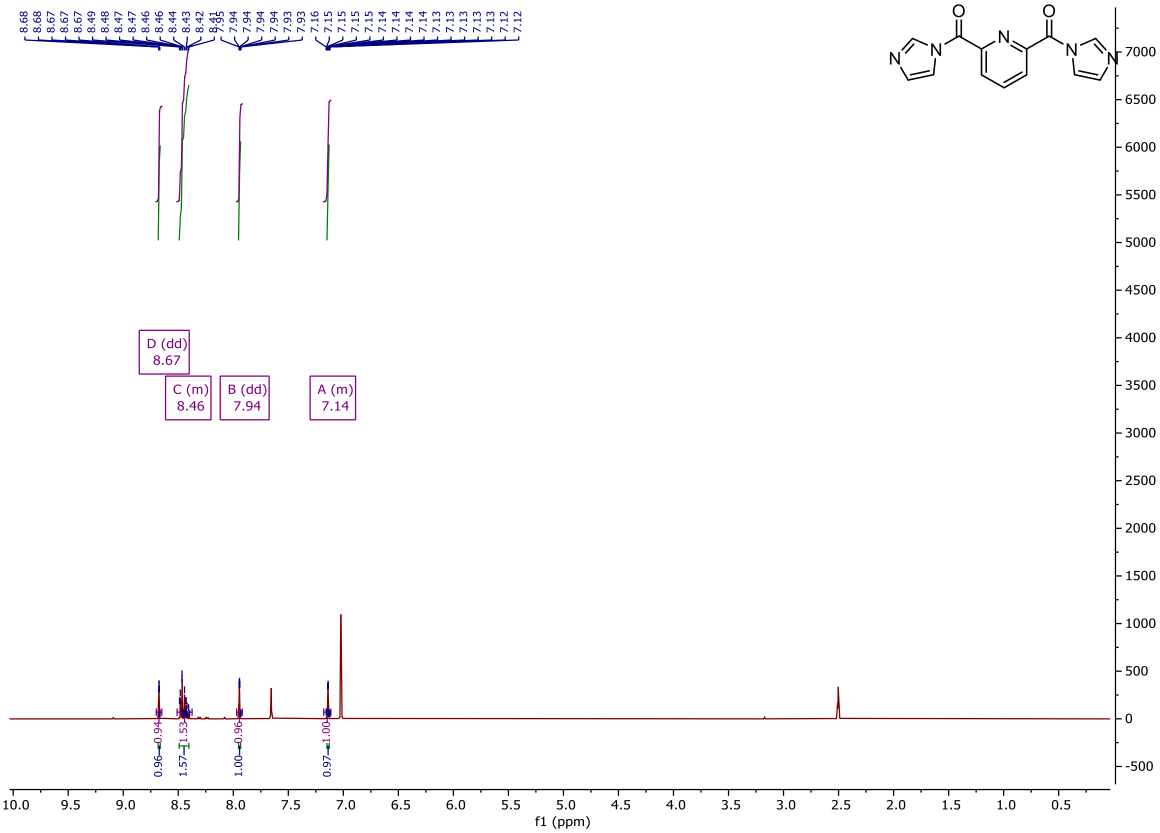

D (dd)
8.67
C (m)
8.46
B (dd)
7.94
A (m)
7.14
f1 (ppm)

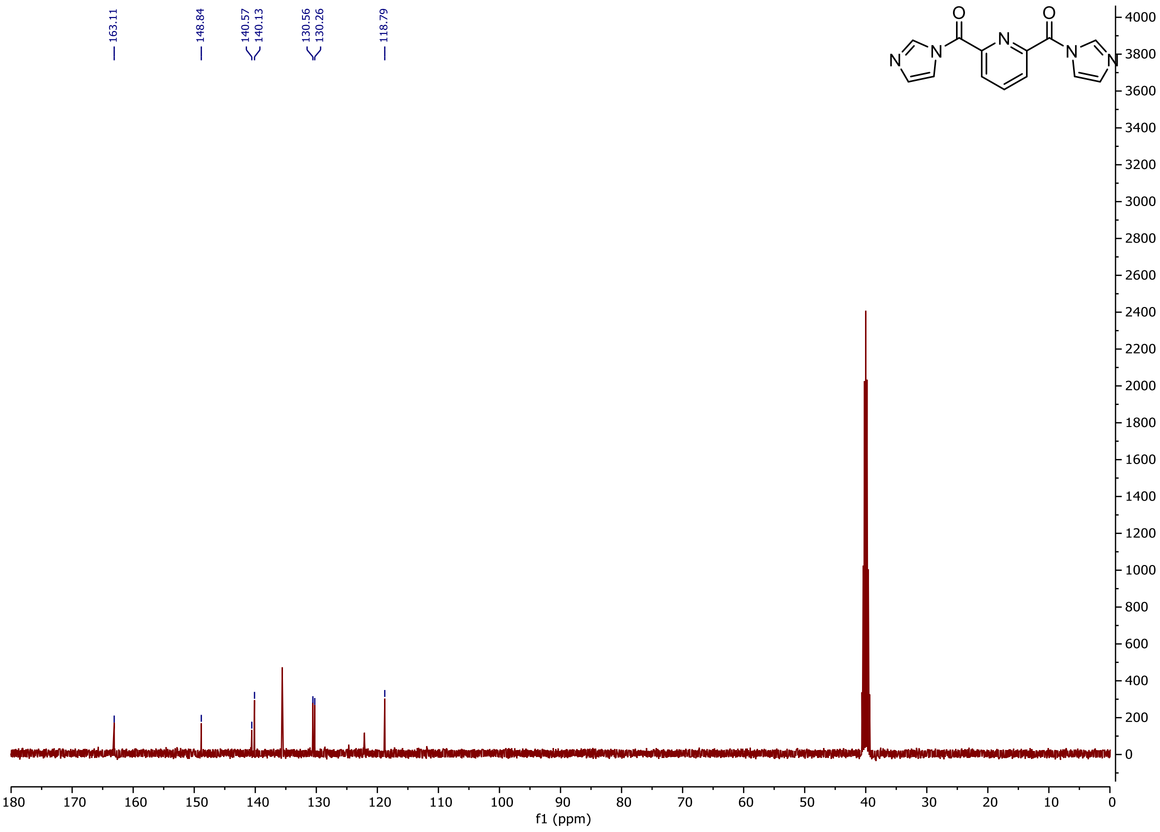

163.11
148.84
140.57
140.13
130.56
130.26
118.79
f1 (ppm)


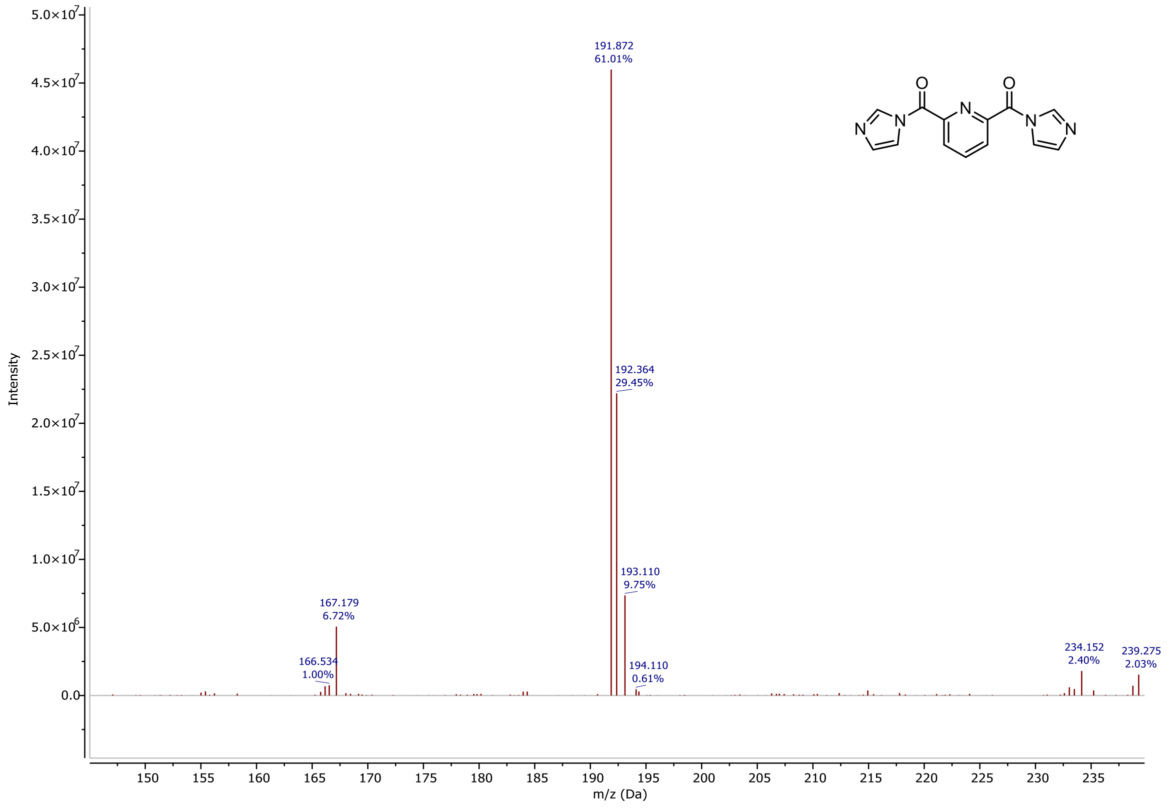

191.872
61.01%
192.364
29.45%
193.110
9.75%
167.179
6.72%
166.534
1.00%
194.110
0.61%
234.152
2.40%
239.275
2.03%
Intensity
m/z (Da)

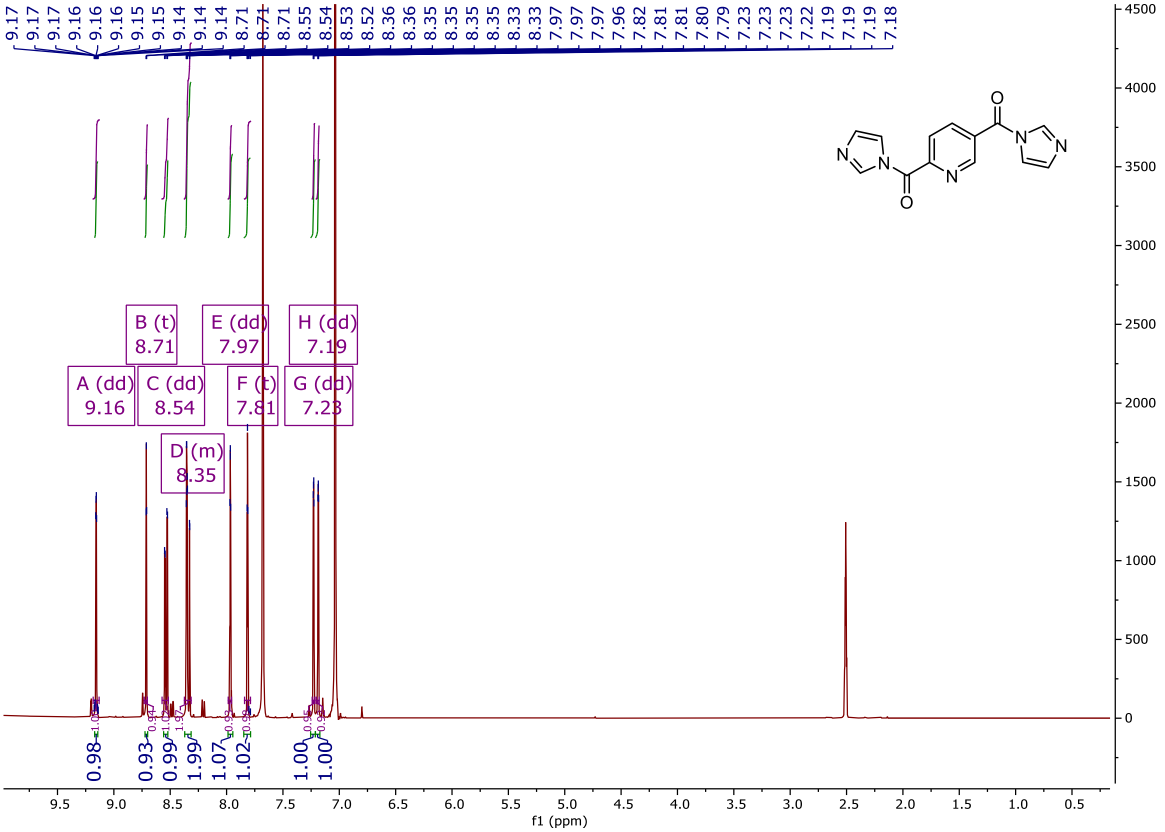
9.17 9.17 9.17 9.16 9.16 9.16 9.15 9.15 9.14 9.14 9.14 8.71 8.71 8.71 8.55 8.54 8.53 8.52 8.36 8.36 8.35 8.35 8.35 8.33 8.33 7.97 7.97 7.96 7.82 7.81 7.81 7.80 7.79 7.23 7.23 7.23 7.22 7.19 7.19 7.19 7.18
A (dd) 9.16
B (t) 8.71
C (dd) 8.54
D (m) 8.35
E (dd) 7.97
F (t) 7.81
G (dd) 7.23
H (dd) 7.19
0.98 0.93 0.99 1.99 1.07 1.02 1.00 1.00
f1 (ppm)

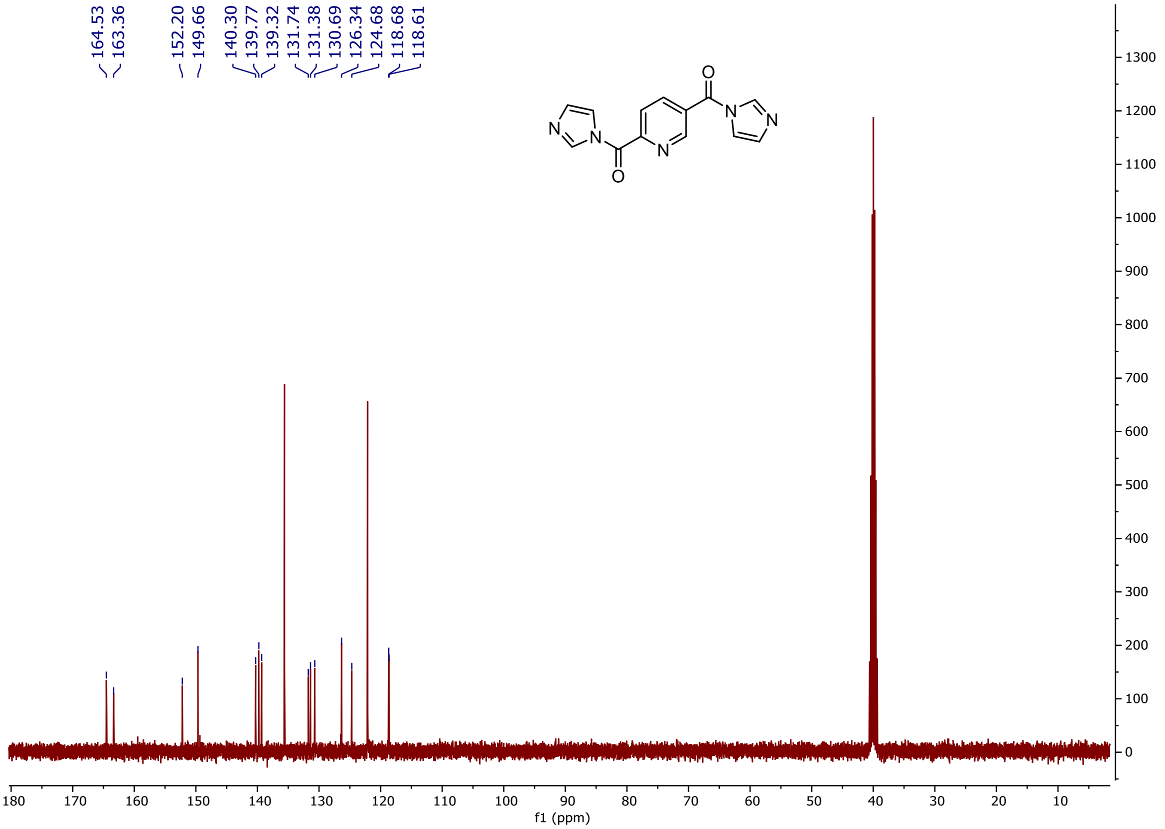
164.53 163.36 152.20 149.66 140.30 139.77 139.32 131.74 131.38 130.69 126.34 124.68 118.68 118.61
f1 (ppm)

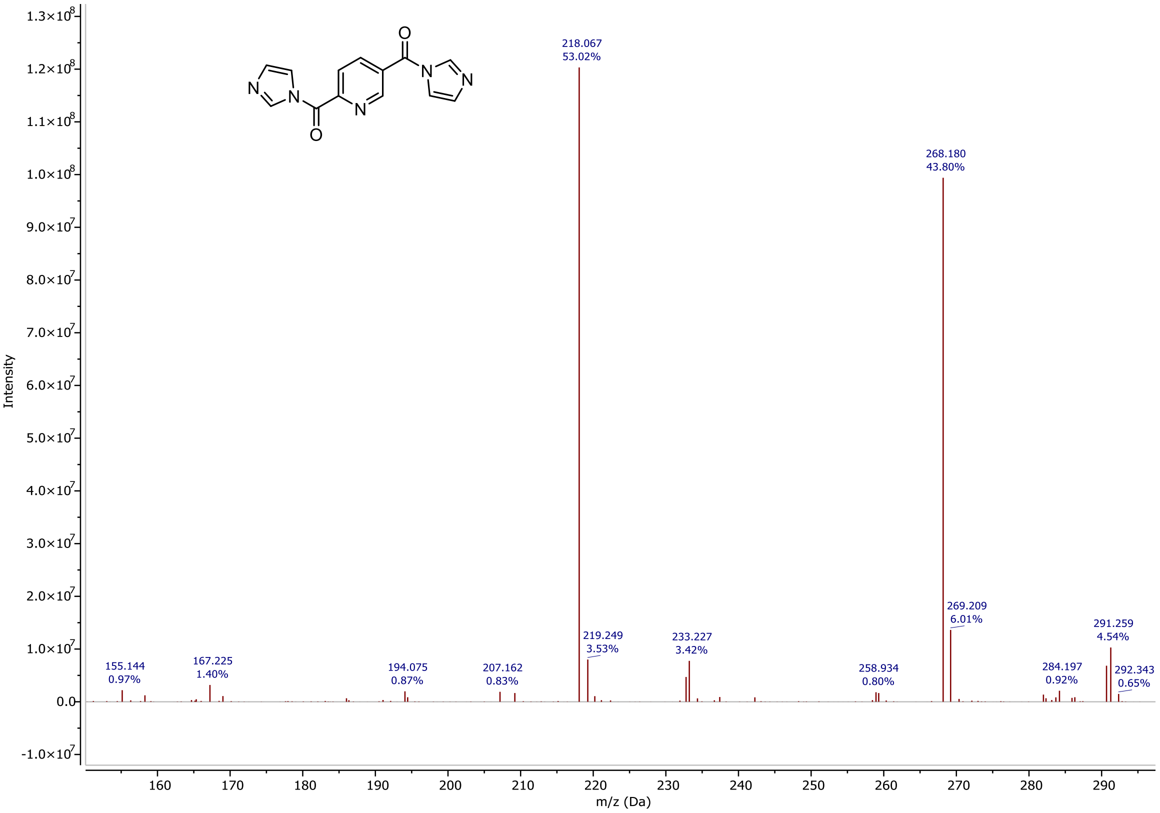

218.067
53.02%
268.180
43.80%
155.144
0.97%
167.225
1.40%
194.075
0.87%
207.162
0.83%
219.249
3.53%
233.227
3.42%
258.934
0.80%
269.209
6.01%
284.197
0.92%
291.259
4.54%
292.343
0.65%
Intensity
m/z (Da)


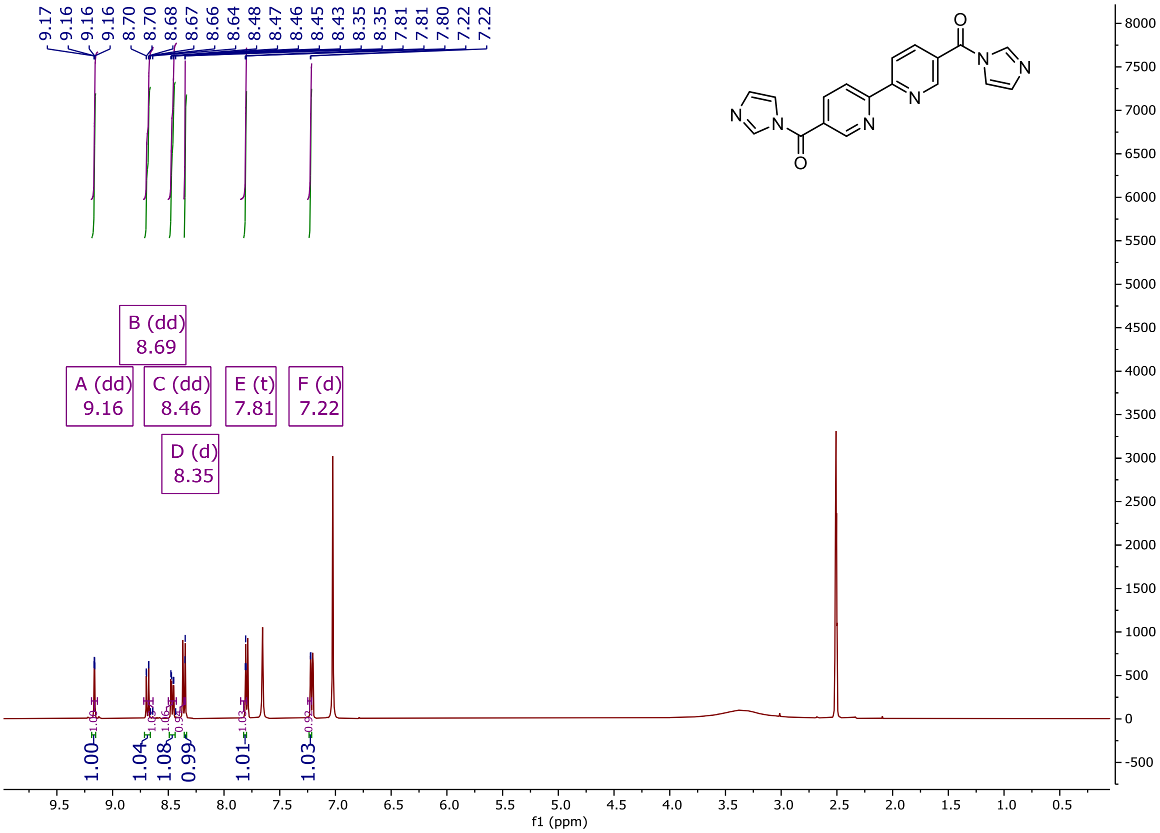

9.17
9.16
9.16
9.16
8.70
8.70
8.68
8.67
8.66
8.64
8.48
8.47
8.46
8.45
8.43
8.35
8.35
7.81
7.81
7.80
7.22
7.22
B (dd)
8.69
A (dd)
9.16
C (dd)
8.46
E (t)
7.81
F (d)
7.22
D (d)
8.35
1.00
1.04
1.08
0.99
1.01
1.03
f1 (ppm)

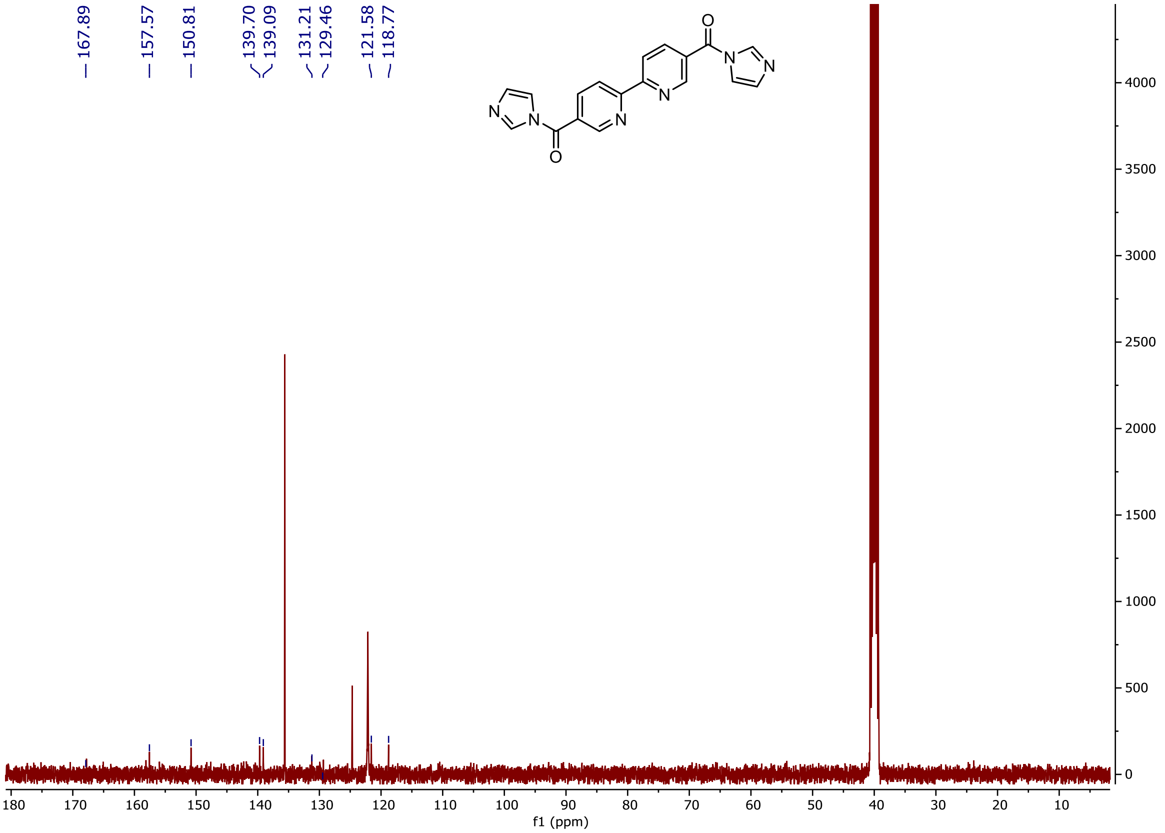

167.89
157.57
150.81
139.70
139.09
131.21
129.46
121.58
118.77
N
O
f1 (ppm)


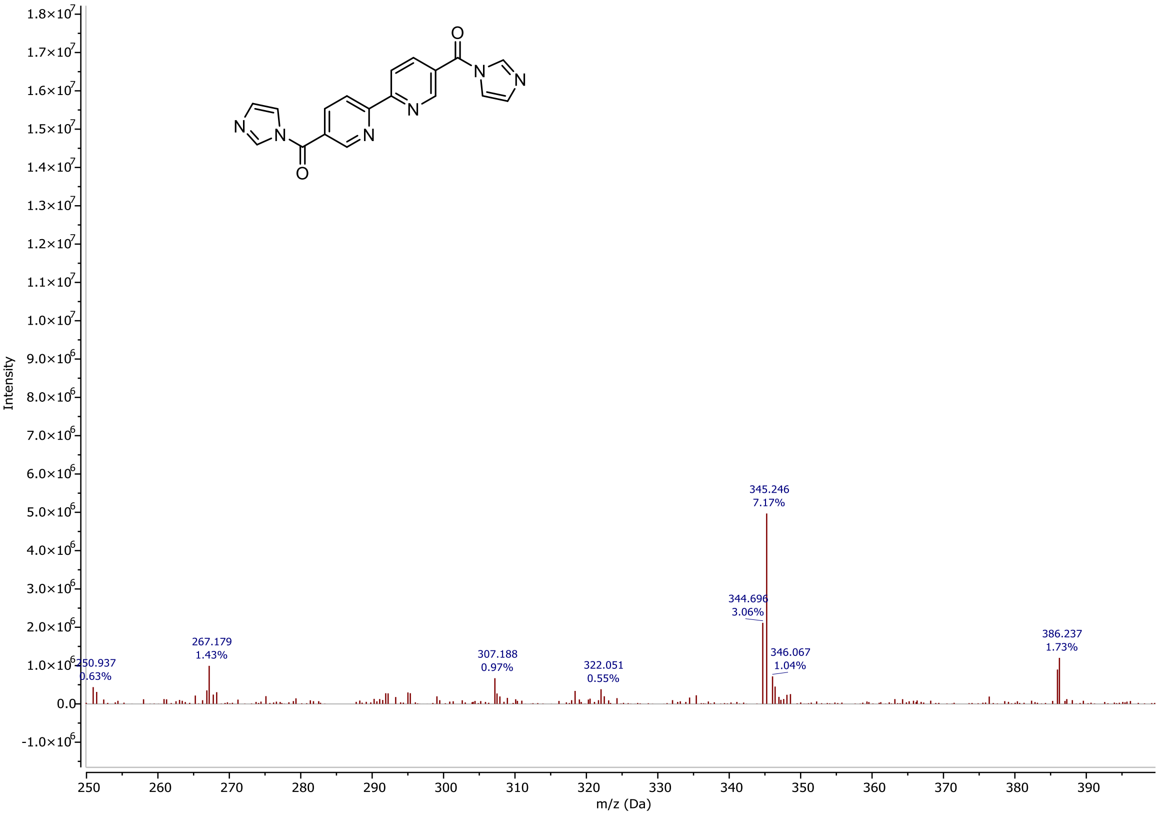

Intensity
m/z (Da)
345.246
7.17%
344.696
3.06%
346.067
1.04%
386.237
1.73%
307.188
0.97%
322.051
0.55%
267.179
1.43%
250.937
0.63%

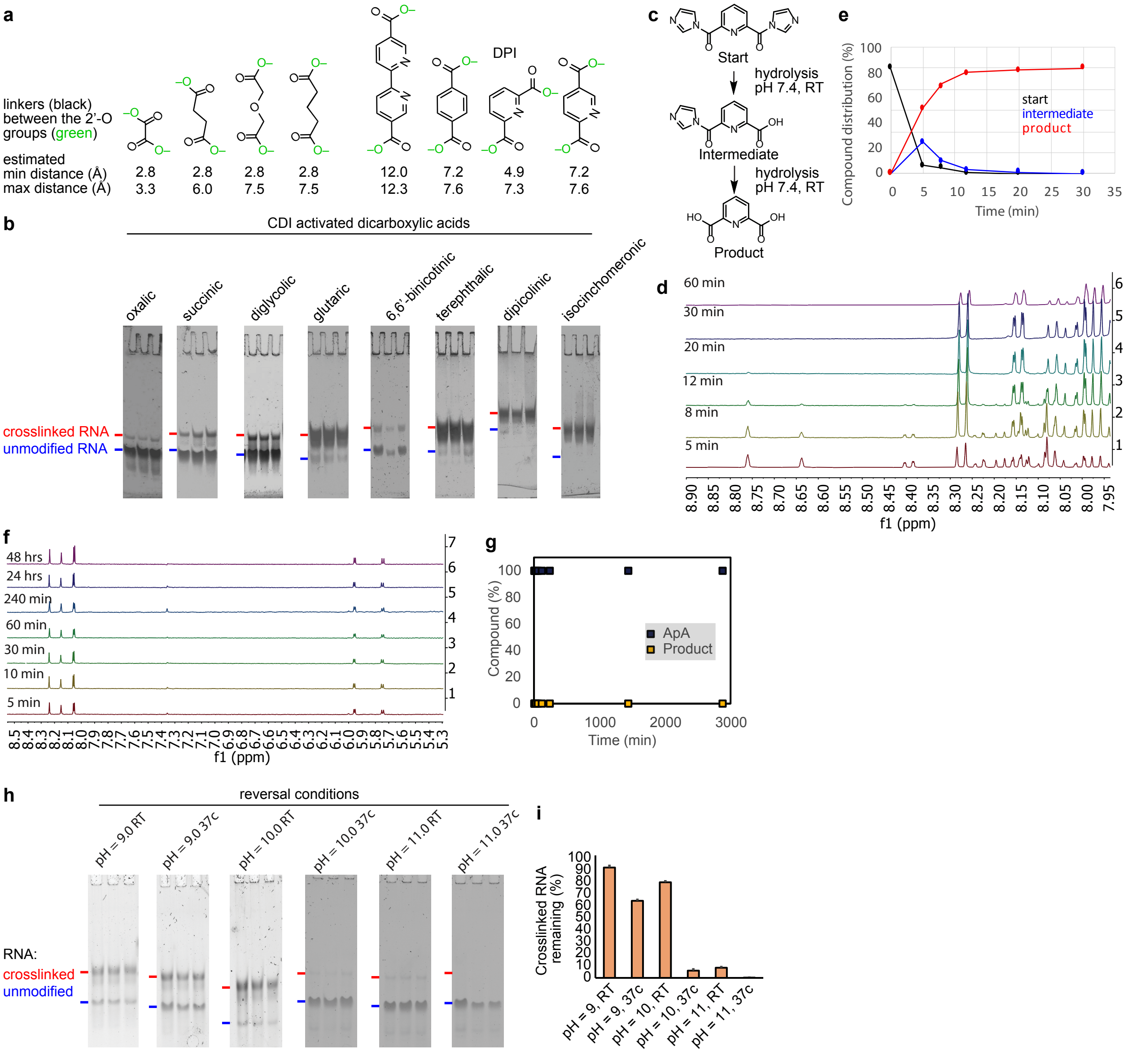


**Supplementary Figure 2. Characterization of SHARC chemistry. a**, Minimal and maximal possible lengths between the crosslinked 2' oxygen groups in RNA. The 3D chemical structures were either extracted from PDB (3U8G for oxalic acid, 5GIB for succinic acid, 6WTX for terephthalic acid and isocinchomeronic acid, 4IH3 for dipicolinic acid) or modeled using ChemDoodle (glutaric acid and 6,6'-binicotinic acid), and then the spatial distances were estimated in PyMOL. Hydroxyl groups are represented by the green oxygen atoms. The minimal distances were set at 2.8, approximately the distance between two oxygens between hydrogen-bonded water molecules. **b**, PAGE analysis of crosslinking efficiency with the different activated dicarboxylic acids (100 mM), with 10 μM model RNA 1 in 0.06 M MOPS, pH 7.5; 0.1 M KCl; 2.5 mM MgCl2 at room temperature for 4 hours. The three lanes represent triplicate experiments. **c**, Reaction scheme of hydrolysis reaction of DPI. **d**, Hydrolysis of DPI in phosphate buffer pH 7.4 analysed by NMR spectroscopy over time. At time t=5 min, the hydrolysis has started and a mixture of starting material, mono-hydrolyzed and di-hydrolyzed DPI is present. At t=60 min the hydrolysis is complete and only product is present (dipicolinic acid and imidazole). Because the spectra are obtained in deuterated buffer t=0 is omitted: the reaction has started before the first spectra can be obtained. The spectrum of unreacted DPI in DMSO-d6 can be found in the experimental section. **e**, Formation of hydrolyzed DPI products over time. After ~30 min all DPI has been hydrolyzed. **f**, Hydrolysis of ApA in 100 mM borate buffer pH 10.0 analysed by NMR spectroscopy over time. **g**, Formation of hydrolyzed ApA products over time, based on quantification of panel f. Even after 48 hours, no hydrolysis products are observed. **h**, PAGE analysis of crosslink reversal efficiency at different alkaline and temperature conditions (37 C or room temperature RT). The three lanes represent triplicate experiments. **i**, An increase in crosslink reversal (= decrease in crosslinked RNA) is observed at increased pH and temperature, based on quantification of gel pictures in panel h. Near complete reversal was achieved at pH 10-11 without obvious damage.

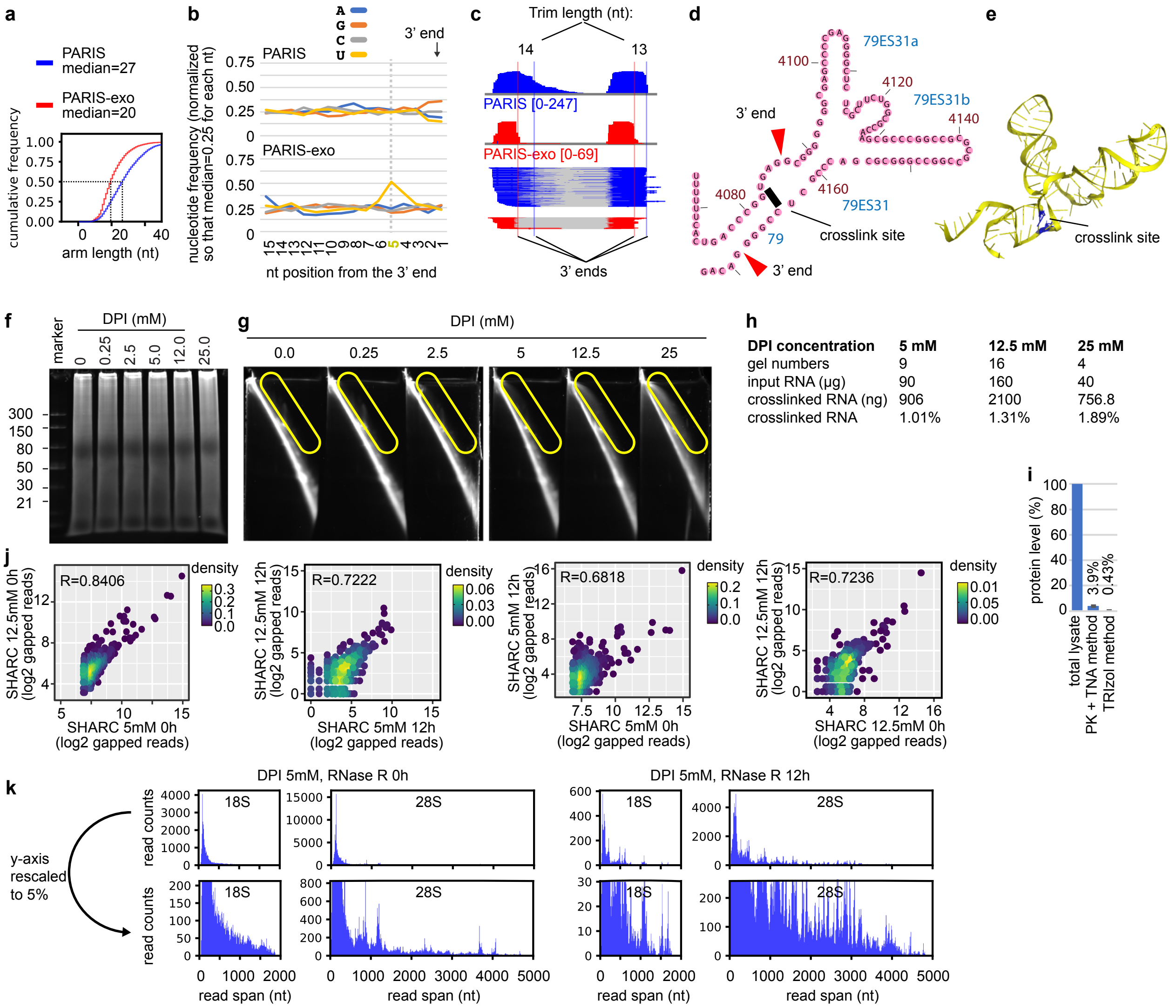


**Supplementary Figure 3. Establishing the exo method for PARIS (a-e) and SHARC (f-j). a,** RNase R reduces arm length in PARIS-exo. **b,** RNase R trimming leads to enrichment of U at the 5th nucleotide from the 3' end. **c-e,** An example of precise crosslink site identification using PARIS-exo. **c**, Blue and red vertical lines indicate the 3' ends (median) from PARIS and PARIS-exo reads. **d,** The PARIS-exo derived 3' ends and crosslinking sites are mapped to the H79 and ES31 of human 28S secondary structure. **e,** cryo-EM structure of H79 and ES31 (PDB: 4V6X). **f**, HEK293 cells are crosslinked with DPI at different concentrations. Total RNA crosslinked cells are fragmented by RNase III and separated on a 8% denatured urea-TBE gel. **g**, RNA fragments from the first dimension (panel f) were electrophoresed again on a second dimension of 16% urea-TBE gel. The smear above the diagonal represents crosslinked RNA. **h**, Quantification of the recovery of crosslinked RNA fragments from the DD2D gel system (replicates n = 9, 16 and 4 for the 3 conditions, respectively). The increase in yield is not linear in response to higher crosslinker concentration, because most accessible crosslinking sites have reacted at lower concentrations, yielding an increase in concentration ineffective. **i,** To measure protein content in crosslinked and purified RNA samples, we first crosslinked cells with 5mM DPI. We prepared, in triplicates, (1) total cell lysate in RIPA buffer, (2) RNA extracted using the PK and TNA method, and (3) RNA extracted using standard TRIzol method. All samples were measured for protein concentration using the BCA method, and values normalized against the total lysate. Relative protein concentrations for the PK+TNA method: 4.66%, 3.58%, 3.41%. Relative protein concentrations for the TRIzol method: 0.28%, 0.66%, 0.34%. **j**, Scatter plot of gapped reads in each duplex group (DG) among experiments with different conditions. **k**, The span of SHARC (5mM DPI, no RNase R trimming) and SHARC-exo (5mM DPI and 12 hour RNase R) gapped reads mapped to the ribosomal RNAs 18S (1869nt) and 28S (5070nt). The lower panels are the same data as upper, with the y-axis rescaled to 5% to show the longer-distance reads.

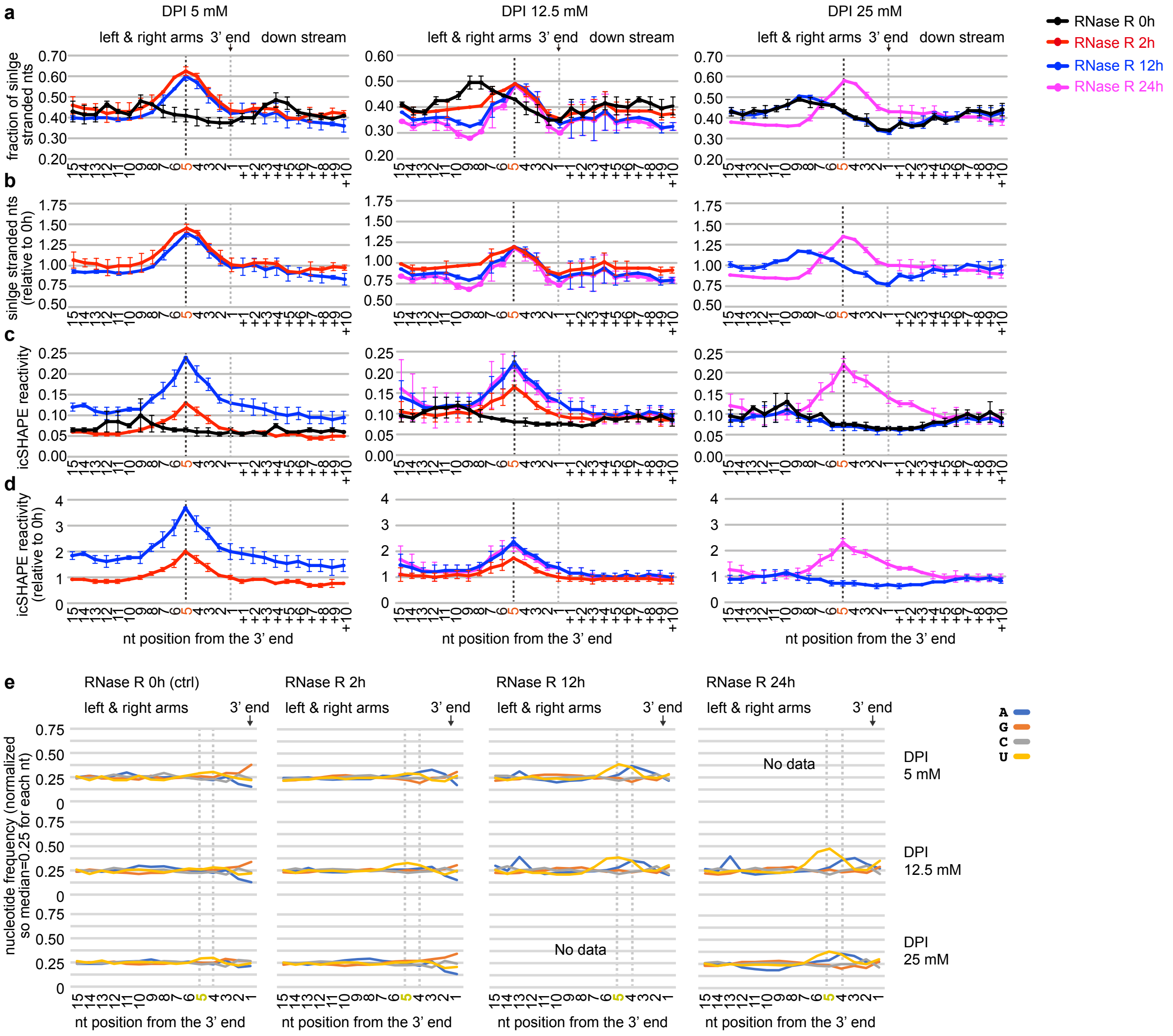


**Supplementary Figure 4. SHARC crosslinking and RNase R trimming reveal crosslink sites with high resolution.** The 28S rRNA was analyzed for all SHARC experiments. **a,** Fraction of single-stranded nucleotides around the 3' end of the left and right arms. Single stranded nucleotides were defined based on the human ribosome cryo-EM structure (PDB: 4V6X). RNase R trimming leads to a dramatic enrichment of single-stranded nucleotides (ss-nts) around the 5th nucleotide, marked by the black vertical dashed line. **b,** Quantification of differences between RNase R trimmed samples vs. non-trimmed PARIS or SHARC data shown in panel **a**, at the 5th nucleotide position upstream of the 3' end. **c,** Average icSHAPE reactivity around the 3' end of the left and right arms, showing better enrichment of accessible nucleotides at the 5th position than panel **a**. **d,** Quantification of differences between RNase R trimmed samples vs. non-trimmed SHARC data shown in panel **c**, at the 5th nucleotide position upstream of the 3' end. In panels a and c, the higher signal for the non-trimmed SHARC data between 7 and 15 reflects the diffused higher probability single stranded regions. This higher signal collapsed around the 5th nucleotide upon RNase R trimming. **e**, Frequencies of the 4 nucleotides around the 3' end of the left and right arms. Stronger RNase R trimming revealed a slight enrichment of A and U near the 5th nucleotide, likely reflecting the weaker secondary structure constraints near the SHARC crosslinking sites.

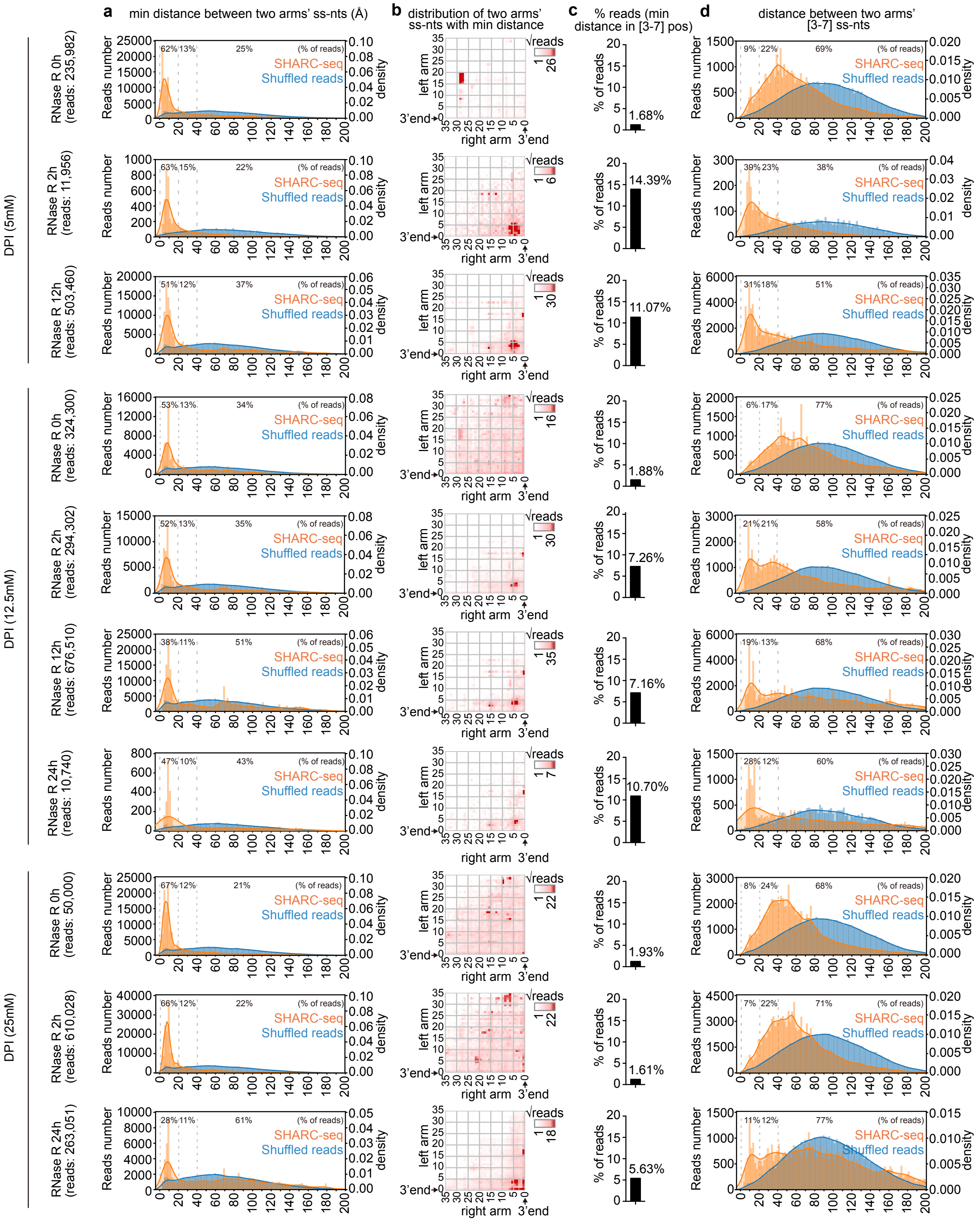

**Supplementary Figure 5. Characterization of SHARC crosslinking and RNase R trimming conditions. a**, The distribution of minimal distances between two arms' ss-nts (single-stranded nucleotides). **b**, Heatmap showing the positions of two arms' ss-nts at min distance; **c**, Percentage of reads, with minimal distance located in [3,7] x [3,7] positions. **d**, The distribution of distances between the two arms' 3rd to 7th ss-nts. In panels a and d, kernel distributions are represented by lines and labeled on the right; percentages of reads with distances in 0-20, 20-40 and >40 Angstroms.

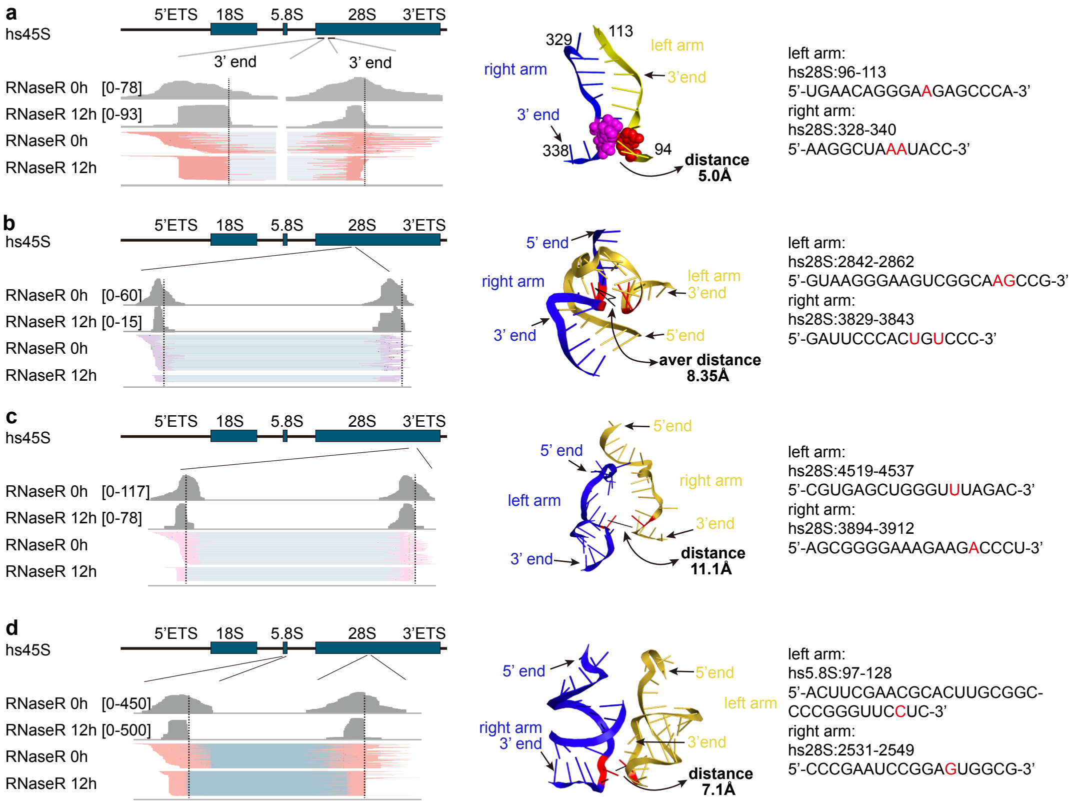


**Supplementary Figure 6. Example secondary and tertiary proximal nucleotides crosslinked by SHARC-exo in the ribosome.** SHARC sequencing data for the ribosome with or without RNase R treatment are compared (0 vs. 12 hour RNase R) on the left panels. Vertical dash lines represent the median 3' ends for the RNase R trimmed reads. In the middle, positions of 3' ends, crosslinking sites and distances are labeled on the cryo-EM model of the ribosome (PDB:4V6X). Sequences of the region are shown on the right. Panels **a-b** are examples of spatial proximity constrained by secondary structures. Panels **c-d** are examples of tertiary contacts either within (**c**) or between (**d**) RNA molecules.

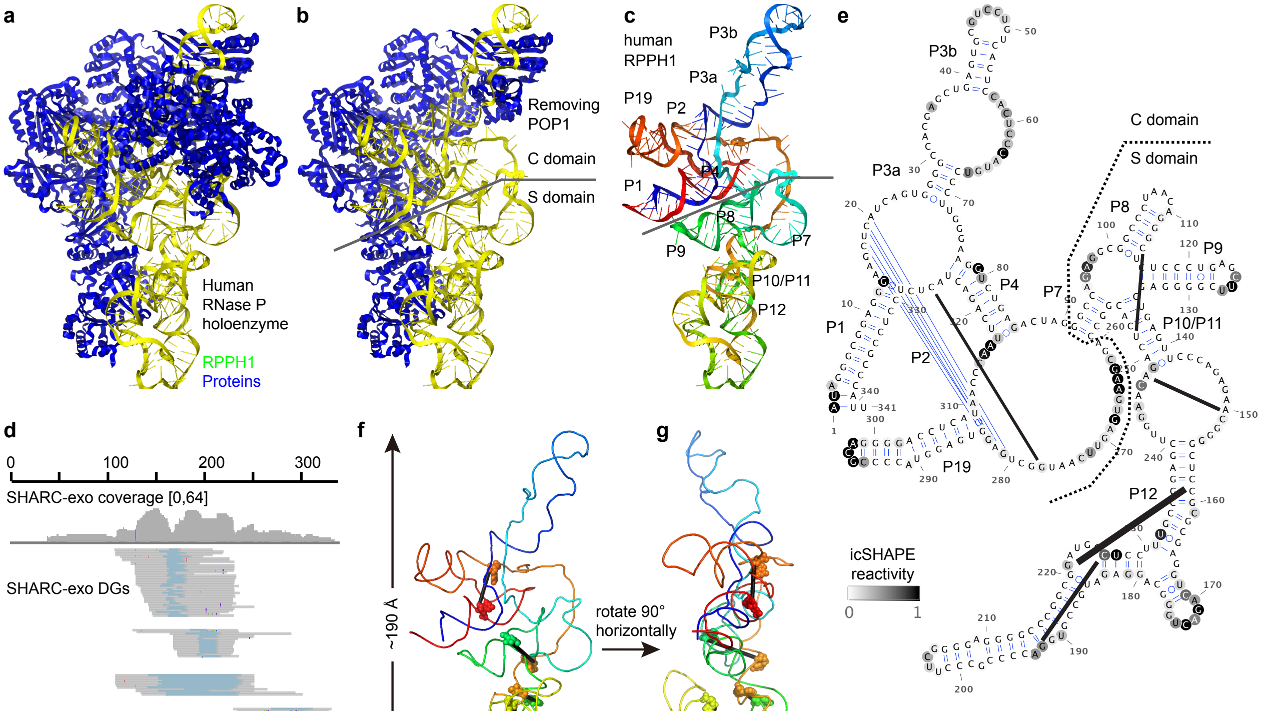


| reads | coord_1 | coord_2 | distance (Å) | domain |
|---|---|---|---|---|
| 39 | 159 | 221 | 26.2 | S |
| 16 | 192 | 226 | 35.1 | S |
| 12 | 150 | 248 | 17.3 | S |
| 9 | 114 | 252 | 35.4 | S |
| 7 | 277 | 326 | 18.5 | C |

**Supplementary Figure 7. SHARC-exo captures spatial distances in the RPPH1 RNA in human RNase P. a**, Cryo-EM structure of the RNase P holoenzyme, which contains 1 RNA and 10 protein partners (PDB: 6AHR). **b**, The holoenzyme contains an RNA core, the C domain, stabilized by extensive tertiary interactions, and the S domain, which is largely exposed and potentially dynamic (The removed protein POP1 is chain B in 6AHR). **c**, Helices in RPPH1: P1-P19, color-coded blue to red from the 5' end to the 3' end. **d**, SHARC-exo data showing all DGs with >5 reads each. **e**, icSHAPE and SHARC-exo (black lines) data overlapped onto the secondary structure of the RPPH1 RNA. icSHAPE data were extracted from our recent study (Lu et al. 2016 Cell, PMID: 27180905). Thickness of the black lines are scaled to the square root of the read numbers shown at the bottom. Coord_1 and coord_2 are the two crosslinked nucleotides in each DG. **f-g**, Crosslinking sites mapped to the 3D structure of RPPH1, in two views rotated horizontally. Crosslinked nucleotides are shown in spheres. A straight black line was drawn between each pair of nucleotides at the 2'O positions.

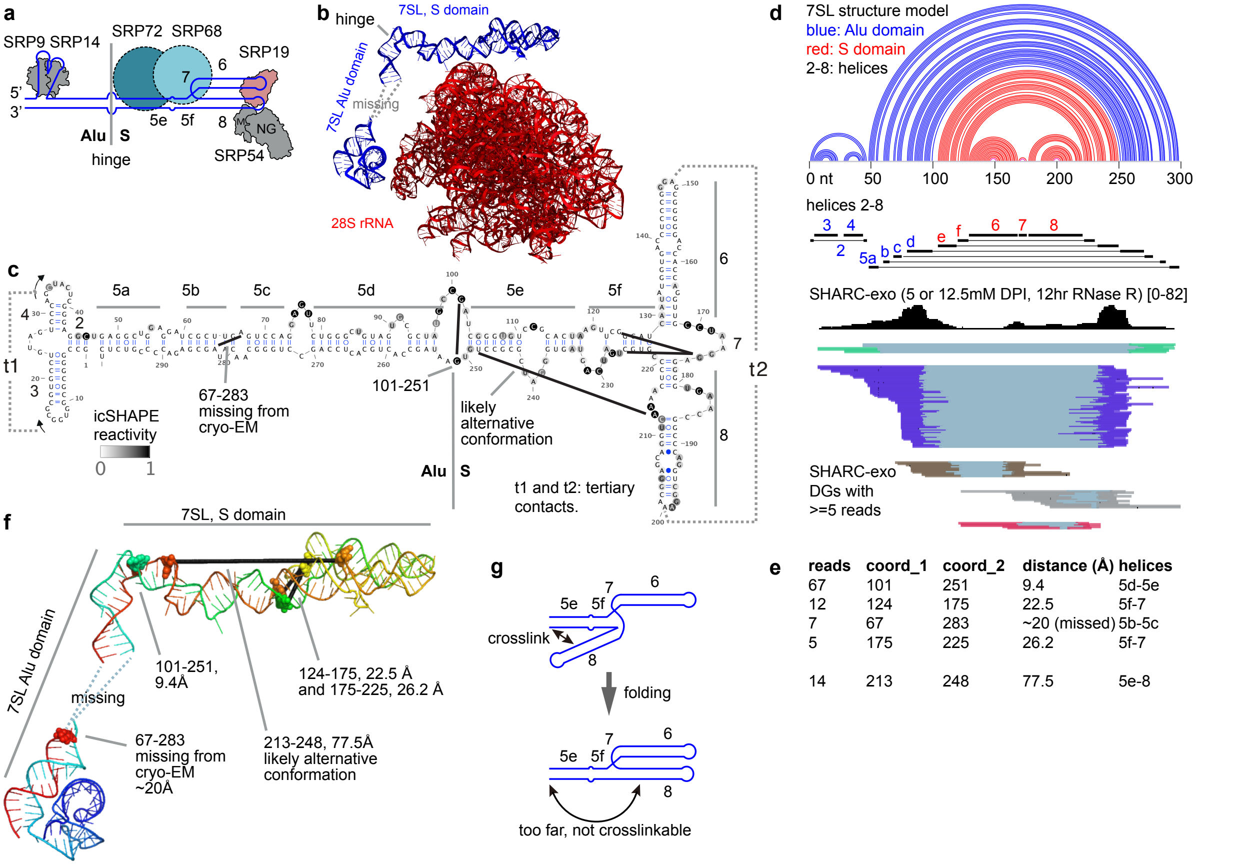


**Supplementary Figure 8. SHARC-exo measures spatial distances in the 7SL RNA**. **a**. Model of the SRP complex, which consists of the 7SL RNA and 6 proteins. These components can be organized into 2 major domains, Alu and S, separated by the hinge/elbow. The Alu domain contains helices 2, 3, 4, 5a, 5b, 5c, 5d, and proteins SRP9/14. The S domain contains helices 5e, 5f, 6, 7, 8, and proteins SRP19/54 and SRP68/72, where the SRP54 recognizes nascent peptides from ribosome. Redrawn from Grotwinkel et al. 2014 (PMID: 24700861). **b**. Cryo-EM structure of the 7SL RNA on the 28S rRNA (PDB: 6FRK). A section of the Alu domain is not visible based on the cryo-EM data and therefore marked missing. In PyMOL: set_view (-0.699150383, -0.621524274, 0.353370398, 0.497510254, -0.067985296, 0.864776254, -0.513455927, 0.780423343, 0.356752545, 0, 0, -445.966888428, 258.757995605, 294.736541748, 289.847534180, 271.441589355, 620.492187500, -20). **c**, Secondary structure model of the 7SL RNA based on the cryo-EM and icSHAPE data (extracted from Lu et al. 2016 Cell, PMID: 27180905). The 5 SHARC-exo crosslinking sites are marked by black lines. t1 and t2 are tertiary contacts in the cryo-EM model. **d**, SHARC-exo data supporting the spatial proximities. Top panel shows the secondary structure, where the Alu and S domains are color coded. A total of 5 DGs were identified with at least 5 reads in each DG. **e**, Numbers of reads, sequence coordinates, spatial distances and helices for the 5 DGs. The one on the bottom (213-248) likely represents an alternative conformation of the helix 8. **f**, Mapping the SHARC-exo derived crosslinking sites onto the cryo-EM structure of 7SL. Color blue to red, from the 5' end to the 3' end. The nucleotide 67 is missing from the cryo-EM, therefore, the distance is a rough estimate. Nucleotides 101 and 251 are right next to each other, and overlapped in this view. **g**, A model of the alternative conformations in 7SL S domain (Redrawn from Kuglstatter et al. 2002). The folded conformation is necessary for SRP19 binding, which then recruits SRP54. Unfolding of the packed helices 6 and 8 would allow helix 8 to bend backward to make contact with 5e. This alternative open conformation was previously suggested as an assembly precursor of the SRP complex (Kuglstatter et al. 2002, PMID: 12244299).

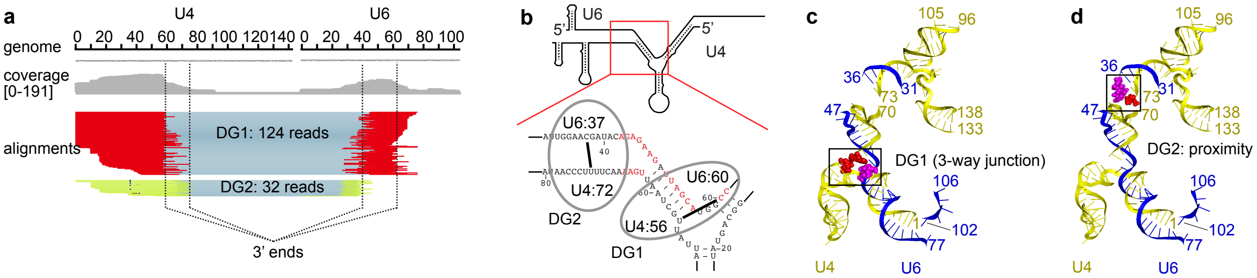


**Supplementary Figure 9. SHARC-exo captures intermolecular interactions in the spliceosomal RNAs. a**, SHARC-exo data for U4-U6 interactions. The numbers of reads and 3' ends of the two DGs are labeled. **b**, Secondary structure model of the U4-U6 dimer (redrawn from Patel and Steitz 2003, PMID: 14685174). SHARC-exo crosslinked sites are labeled. **c-d**, Physical locations of SHARC-exo crosslinking sites on the U5.U4/U6 tri-snRNP cryo-EM structure model (PDB:6QW6). The crosslinked sites of DG2 were missed in cryo-EM structure, but captured by SHARC-exo (U4:72 vs. U6:37). The diagram in panel d shows estimated locations.

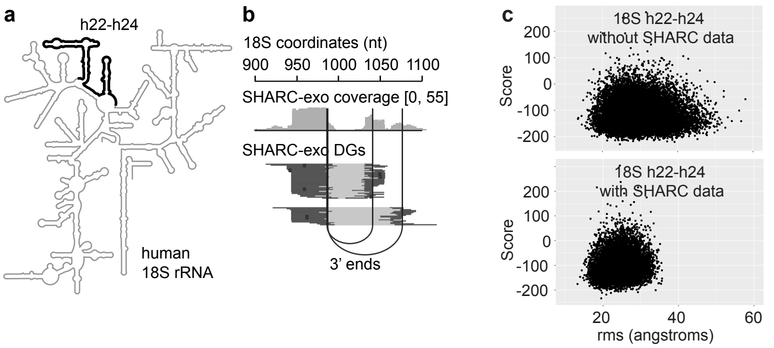


**d** rms values (angstroms)

| | –SHARC | +SHARC |
|---|---|---|
| cluster0 | 24.8 | 19.8 |
| cluster1 | 39.0 | 21.0 |
| cluster2 | 37.4 | 22.3 |
| cluster3 | 28.1 | 20.1 |
| cluster4 | 27.0 | 29.2 |

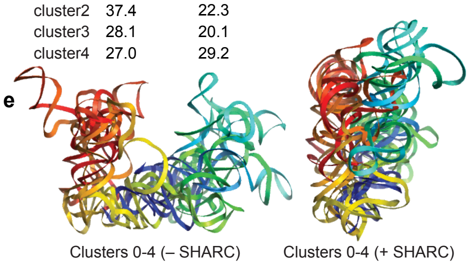


**f** (top 5, >3% reads: DG 0, DG 4, DG 3, DG 1, DG 2)

| | SHARC-exo | | | | SHARC | | | |
|---|---|---|---|---|---|---|---|---|
| | Left Arm | Right Arm | Reads | Distance (Å) | Left Arm | Right Arm | Reads | Distance (Å) |
| DG 0 | 9 | 152 | 43690 | 22.9 | 25 | 152 | 11640 | 34.6 |
| DG 4 | 66 | 152 | 8083 | 19.2 | 85 | 152 | 1791 | 31.9 |
| DG 3 | 12 | 67 | 5992 | 23.1 | 12 | 89 | 4740 | 38.2 |
| DG 1 | 15 | 113 | 4938 | 41.3 | 25 | 122 | 8981 | 64.8 |
| DG 2 | 114 | 152 | 3424 | 24.1 | 124 | 152 | 11303 | 40.9 |
| DG 5 | 66 | 129 | 1652 | 66.5 | 52 | 129 | 1555 | 35.3 |
| DG 6 | 113 | 129 | 1532 | 49.7 | 113 | 129 | 707 | 49.7 |
| DG 7 | 65 | 111 | 1370 | 20.7 | 67 | 123 | 996 | 46.4 |
| DG 9 | 15 | 46 | 1175 | 62.9 | 16 | 47 | 393 | 61.7 |
| DG 16 | 28 | 77 | 603 | 12.9 | 25 | 80 | 563 | 19.0 |
| DG 8 | 41 | 95 | 488 | 54.6 | 38 | 97 | 1250 | 44.0 |
| DG 11 | 67 | 121 | 395 | 37.5 | 69 | 123 | 664 | 45.1 |
| DG 14 | 5 | 27 | 381 | 40.5 | 3 | 27 | 61 | 45.9 |
| DG 10 | 17 | 125 | 274 | 78.9 | 25 | 125 | 1911 | 71.7 |
| DG 13 | 95 | 117 | 139 | 57.1 | 95 | 124 | 319 | 78.5 |
| DG 12 | 29 | 114 | 113 | 54.8 | 29 | 124 | 540 | 72.7 |
| DG 15 | 81 | 144 | 77 | 47.8 | 81 | 138 | 192 | 61.1 |
| DG 17 | 129 | 152 | 74 | 47.7 | 129 | 152 | 45 | 47.7 |

**g**

| DG | left trimmed | right trimmed | distance reduced by (Å) |
|---|---|---|---|
| DG0 | 16 | 0 | 11.7 |
| DG4 | 19 | 0 | 12.7 |
| DG3 | 0 | 22 | 15.1 |
| DG1 | 10 | 9 | 23.5 |
| DG2 | 10 | 0 | 16.8 |

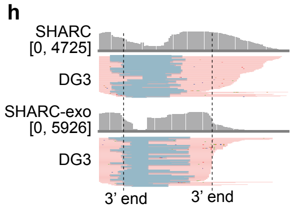


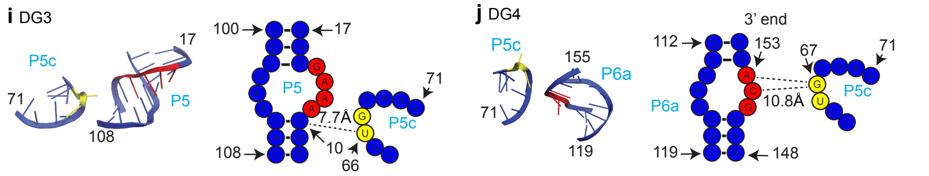


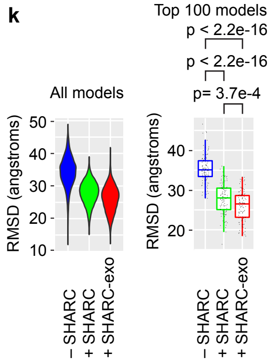


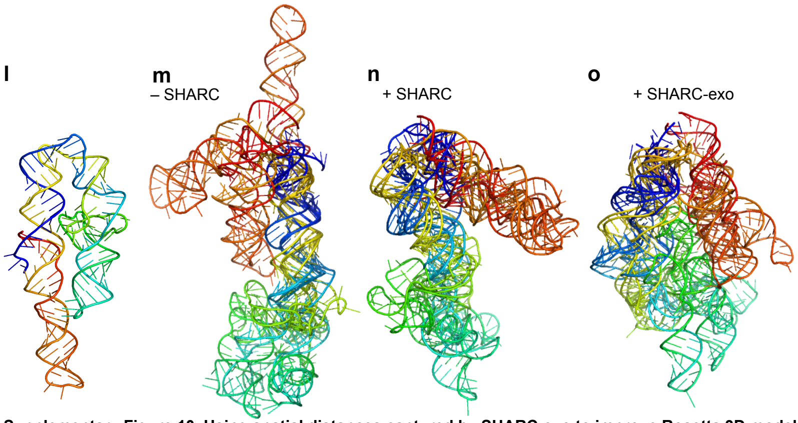


**Supplementary Figure 10. Using spatial distances captured by SHARC-exo to improve Rosetta 3D modeling. a,** Location of the helices h22-h24 (nucleotides 920-1090) in the human 18S rRNA. **b**, SHARC-exo data (5mM SHARC, 12h RNase R), showing DGs that captured two pairs of nucleotides in spatial proximity. Vertical lines indicate the 3' ends of the trimmed reads. **c**, Scatter plots of Rosetta rms (root-mean-square deviation) values vs. scores for the 18S h22-h24 region with or without SHARC-exo constraints. The two constraints removed the vast majority of high rms models (>35 angstroms). **d**, rms values of clusters 0-5 of the 18S h22-h24 segment with or without SHARC-exo constraints. **e**, Alignment of center models for the top 5 clusters (0-4). **f,** Table detailing the number of reads and indicated crosslinking site for each DG for the in vitro P4-P6 SHARC-exo data. DGs with reads more than 3% of the SHARC-exo library are selected for modeling analysis resulting in 5 DGs that account for 89% of SHARC-exo reads and 81% of SHARC reads. **g,** Numbers of nucleotides trimmed from the left and right arms after RNase R treatment (SHARC-exo vs. SHARC), and the improvement of distance measurements (column: distanced reduced by (Å)). **h**, Gapped reads for SHARC (top) and SHARC-exo (bottom) in DG3 (coverage listed in brackets). The vertical lines show the 3' ends (medians) under the SHARC-exo condition. **i-j**, Shortest distances between the two arms (measured on O2' atoms) in DGs 3 and 4. These distances are shorter than the distances measured at the 5th nucleotide from the 3' end, as shown in panel **f**. Un-paired residues most likely to be involved in the crosslinking are depicted in red and yellow. **k**, Violin plot of overall models RMSD as well as boxplot of the top 100 models for each modeling condition (using top 5 DGs from the SHARC and SHARC-exo conditions). Total numbers of models generated by Rosetta are as follows. –SHARC: 13132, +SHARC: 11925, +SHARC-exo: 12651. The –SHARC condition was run without tertiary contact constraints. The statistical significance was calculated using the Wilcox rank sum test. **l,** Crystal structure of the P4-P6 RNA (PDB: 1HR2). **m-o**, Overlap of the top 100 models as 5 clusters, for –SHARC (m), +SHARC (n), and +SHARC-exo (o) conditions.

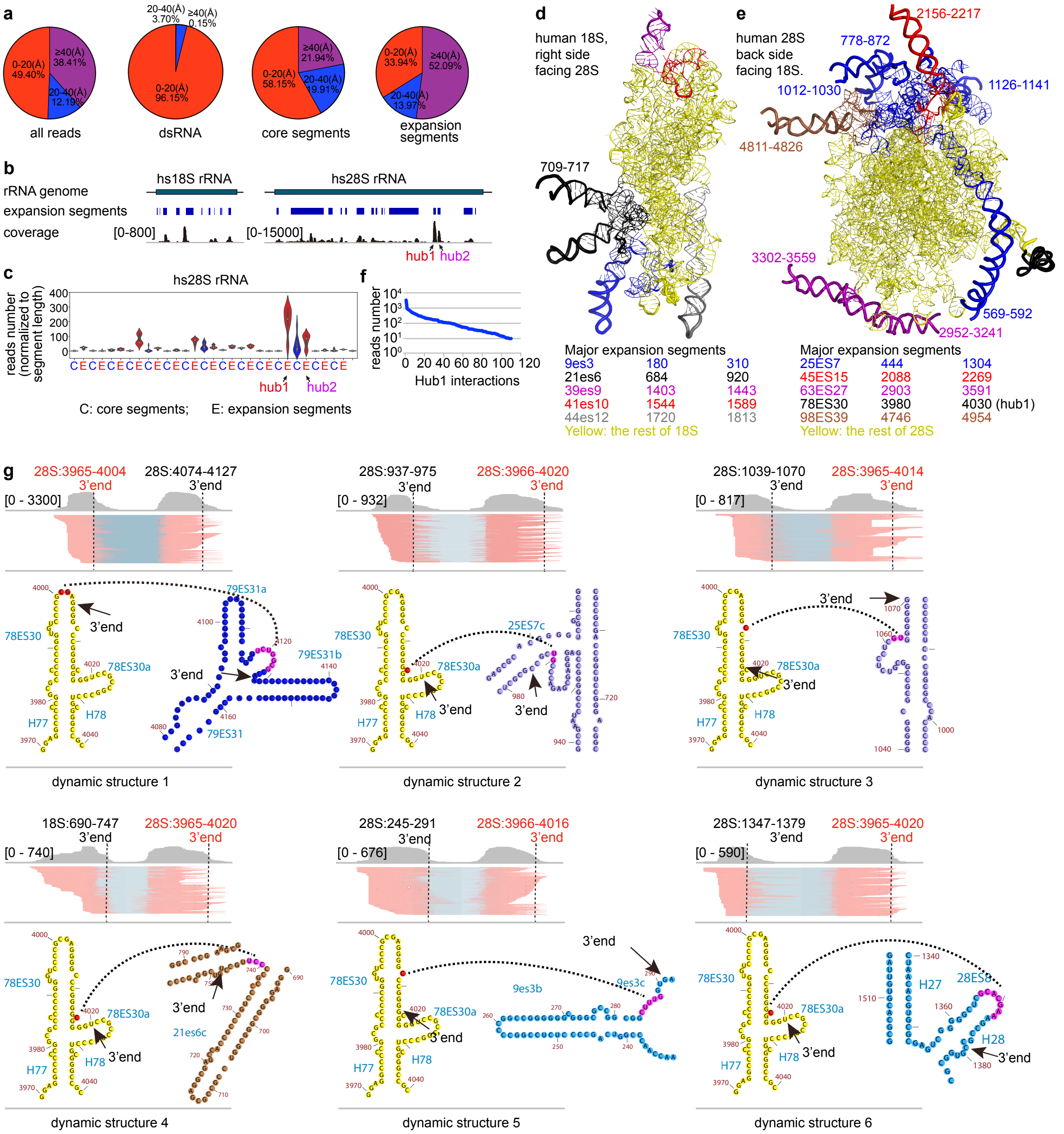

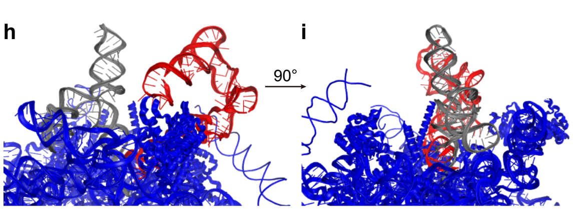


**Supplementary Figure 11. SHARC-exo captures dynamic structures in the human ribosome. a**, Percentages of reads with between-arm distances in three ranges: 0-20, 20-40 and ≥40Å. For crosslinks constrained by dsRNAs, the vast majority of distances (96.15%) are within 20Å. For tertiary contacts in the core and expansion segments, more distances are >20 or >40 Å. **b**, Genome coverage track showing that SHARC-exo reads with larger distance between two arms (≥40Å) are primarily mapped to18S and 28S rRNA expansion segments. The scales are different between 18S and 28S. **c**, Violin plot of per-nt reads coverage along the 28S, in core and expansion segment intervals. **d-e**, Locations of the major expansion segments on the ribosome cryo-EM structures (only expansion segments >50nts are shown). Thin lines with bases represent well positioned regions in the cryo-EM structure, while thick lines without bases represent high B values, i.e. flexible regions. Missing segments are listed next to the break points. Some of the expansion segments, even though are well positioned in this model, may be more flexible in cells, e.g. the roots of 21ES6, 44es12, and 63ES27. For example, 63ES27 missed two regions 2952-3241 and 3302-3559, which add up to 548 nucleotides, the lost of all the missed expansion segments. **f**, Numbers of reads in the hub1-interacting DGs, ranked by coverage. **g**, Details of all the 6 DGs supporting dynamic conformations between 78ES30 and its targets. **h-i**, Model of the alternative interaction between ES30 and ES31, illustrating the lack of clashes with the surface of the ribosome. Two views rotated horizontally by 90° are shown. Red: H76-H78 and ES30. Gray: H79 and ES31.

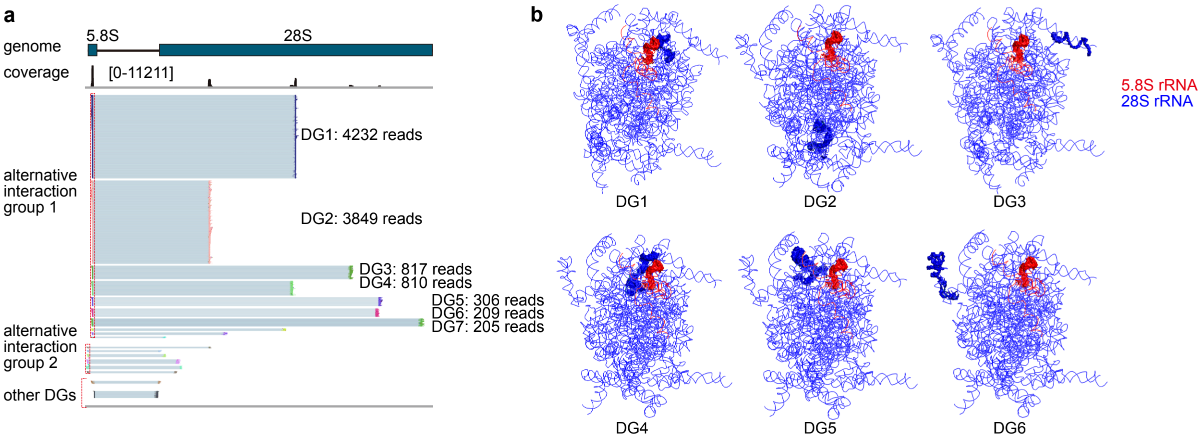


**Supplementary Figure 12. SHARC-exo captures intermolecular interactions between 5.8S and 28S rRNAs.** a, IGV plot showing the interactions between 5.8S and 28S rRNA. Only the duplex groups (DGs) with more than 10 reads were showed here. **b**, Physical locations of top 6 5.8S-28S rRNA interactions on the ribosome cryo-EM structure (PDB: 4V6X). Interacting regions are shown in spheres while the rest are in lines. The part of 5.8S involved in all the alternative contacts is exposed to the surface of the ribosome, making it possible to reach distant 28S helices.

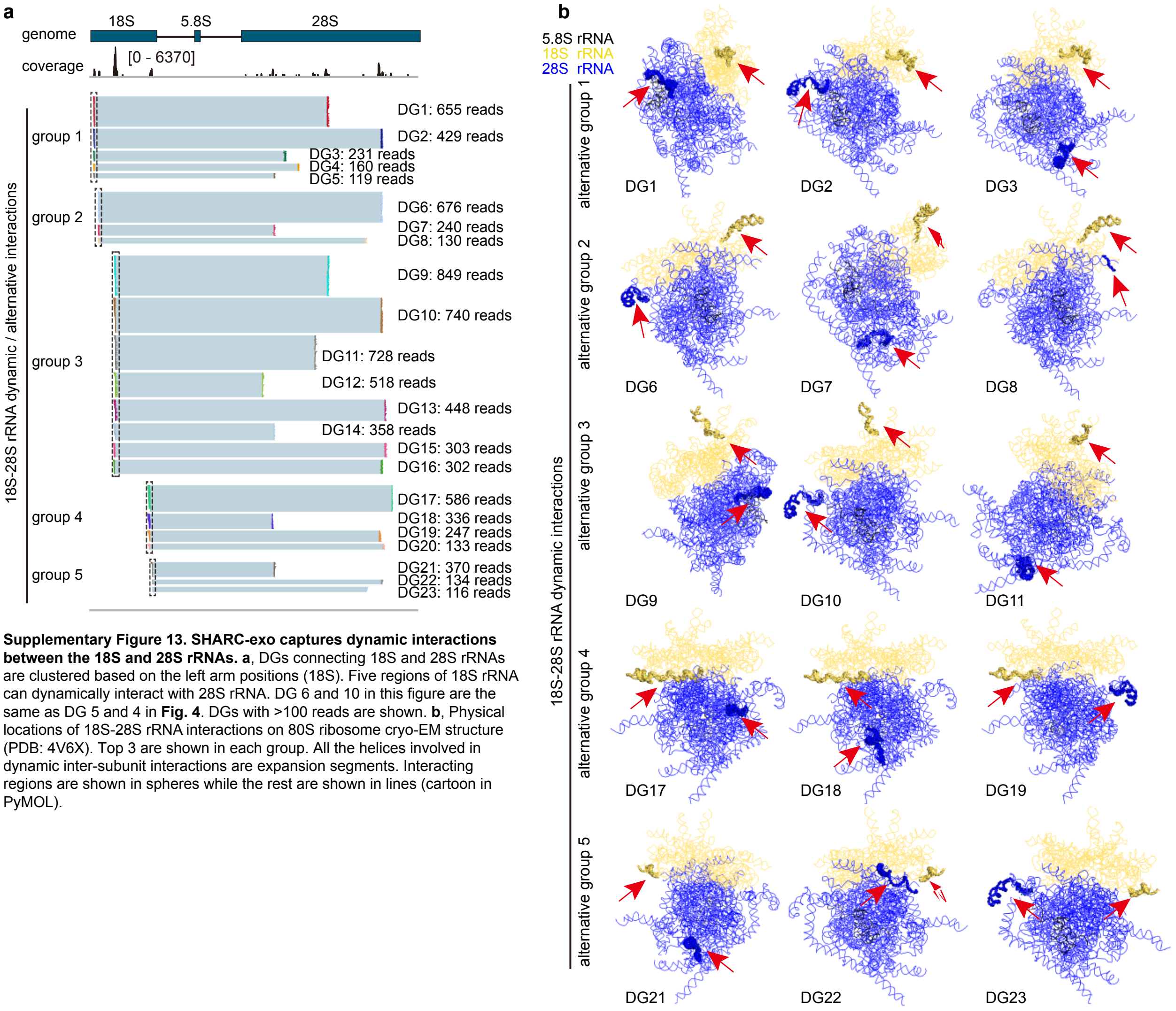


**Supplementary Figure 13. SHARC-exo captures dynamic interactions between the 18S and 28S rRNAs. a**, DGs connecting 18S and 28S rRNAs are clustered based on the left arm positions (18S). Five regions of 18S rRNA can dynamically interact with 28S rRNA. DG 6 and 10 in this figure are the same as DG 5 and 4 in **Fig. 4**. DGs with >100 reads are shown. **b**, Physical locations of 18S-28S rRNA interactions on 80S ribosome cryo-EM structure (PDB: 4V6X). Top 3 are shown in each group. All the helices involved in dynamic inter-subunit interactions are expansion segments. Interacting regions are shown in spheres while the rest are shown in lines (cartoon in PyMOL).

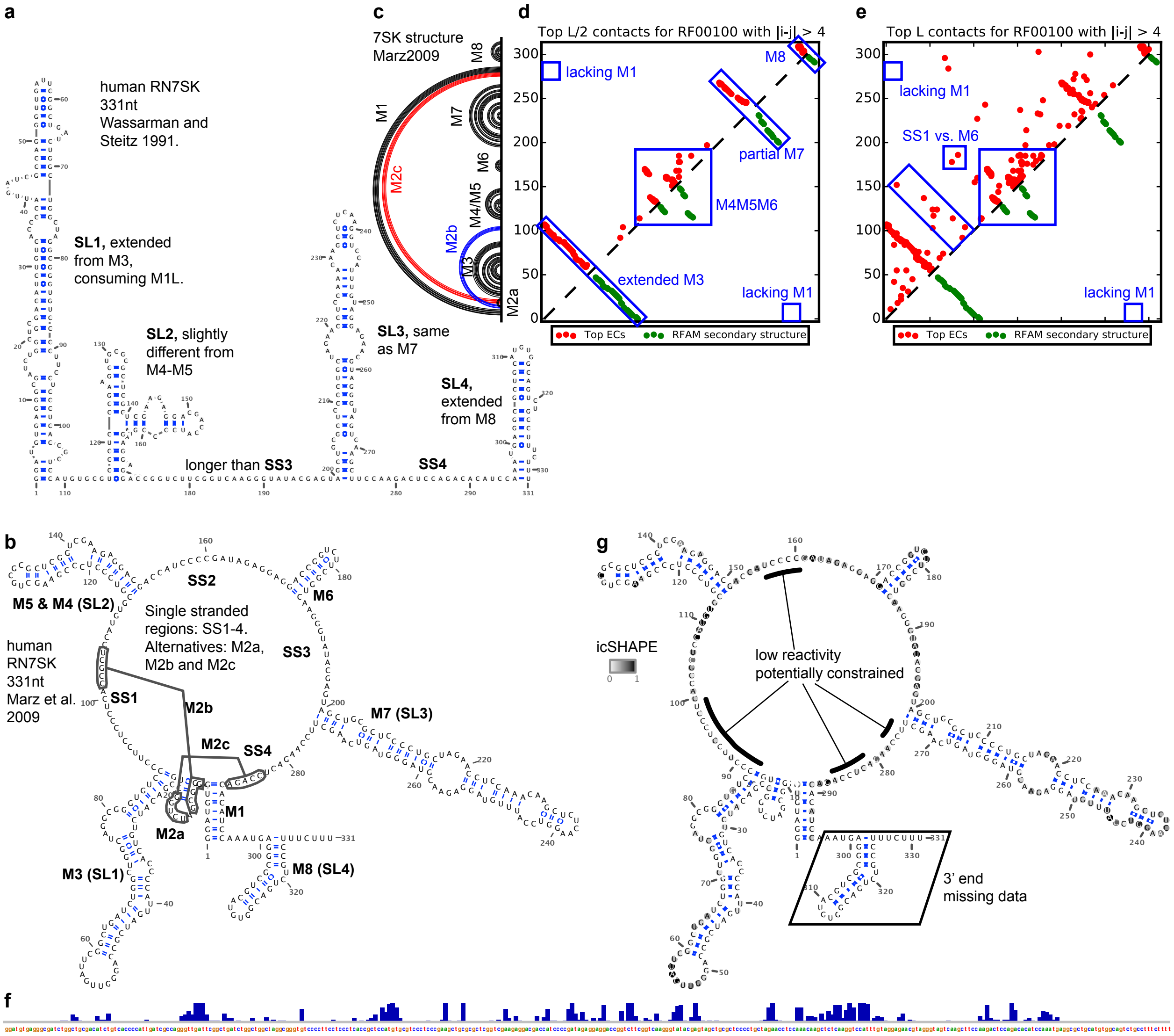


**Supplementary Figure 14. icSHAPE analysis of the human 7SK RNA. a**, Secondary structure model from Wassarman and Steitz 1991. SL1-4 are based on recent nomenclature in crystallographic studies, not the same as the original helices. **b**, Secondary structure model from Marz et al. 2009 (PMID: 19734296). The 8 helices are labeled M1-8, and their alternative names are indicated in the parentheses (SL1-4). The 3 M2 alternative conformations M2a-c and 4 single stranded regions SS1-4 are illustrated. **c-e,** Comparing the Marz model with (c) with EC and Rfam models (d-e). **d-e**, Contacts in the 7SK RNA identified by evolutionary coupling (EC) are shown on the top triangles (Weinreb et al. 2016. PMID: 27087444), while the Rfam model is on the bottom. The top L/2 contacts (d) are by definition more reliable than the top L contacts (e). Top L/2 contacts are almost identical with Rfam secondary structure (Rfam: RF00100), which is consistent with Marz 2009 model. However, the terminal helix M1 was not detected in either Rfam or EC. The extended M3 is also partially inconsistent with the M3 in the Marz model. **f-g**, icSHAPE data from Lu et al. 2016 (PMID: 27180905, panel c) mapped to the Marz 2009 secondary structure model (panel d). icSHAPE data were from HEK 293 cells, without any special treatment. No data were available in the first 5 and last 35 nts due to sliding window processing and primer binding. Constrained regions in the putative single-stranded regions are labeled with thick black curves. These low reactivity nucleotides are likely interacting with proteins or forming tertiary RNA structures that were not captured by phylogenetic analyses such as covariation or evolutionary coupling. However, icSHAPE alone can neither prove nor disprove long-range or tertiary structures.

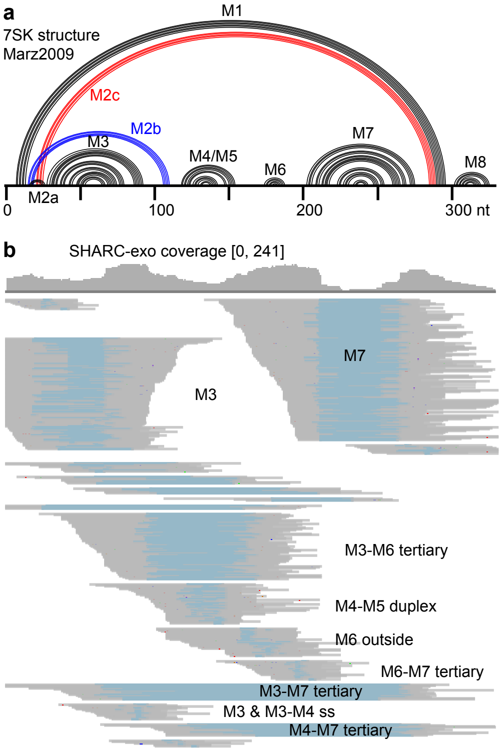


**c**

| DG | Reads | L5 | L3 | R5 | R3 |
|---|---|---|---|---|---|
| 0 | 156 | 162 | 211 | 254 | 290 |
| 1 | 123 | 1 | 24 | 66 | 97 |
| 2 | 73 | 49 | 97 | 155 | 186 |
| 3 | 44 | 89 | 124 | 137 | 159 |
| 4 | 31 | 109 | 160 | 169 | 197 |
| 5 | 21 | 163 | 196 | 200 | 217 |
| 6 | 17 | 17 | 66 | 266 | 284 |
| 7 | 17 | 48 | 88 | 93 | 109 |
| 8 | 13 | 99 | 126 | 266 | 286 |
| 9 | 10 | 238 | 267 | 280 | 325 |
| 10 | 7 | 77 | 110 | 116 | 139 |
| 11 | 10 | 19 | 49 | 68 | 132 |
| 12 | 11 | 3 | 26 | 29 | 49 |
| 13 | 8 | 34 | 62 | 143 | 166 |
| 14 | 6 | 55 | 82 | 186 | 225 |

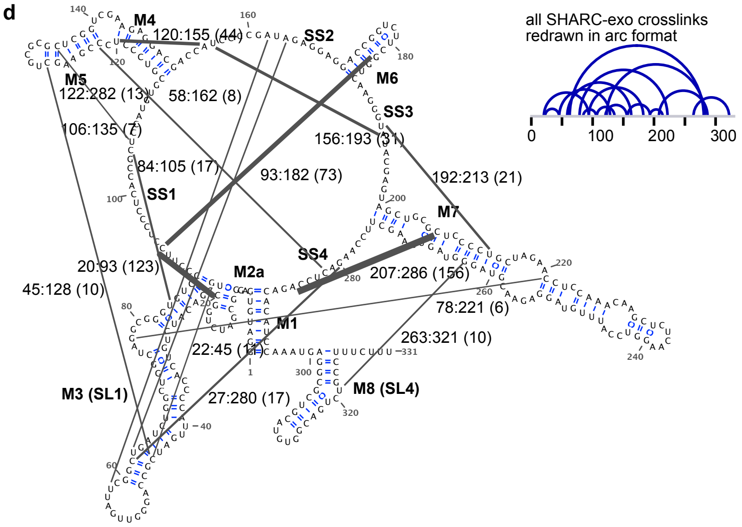


**Supplementary Figure 15. SHARC-exo data for the human 7SK RNA. a,** The Marz model for the 7SK RNA. Blue arcs: M2b. Red arcs: M2c. **b**, SHARC-exo reads coverage and DGs supporting the major helices and interhelical contacts. M8 was not represented in the data due to its small size and tight structure. **c**, Numbers of reads and start/end coordinates for the two arms (L5/L3 for left arm 5' and 3' ends. R5/R3 for right arm 5' and 3' ends). **d,** Mapping the crosslinks to the secondary structure model of 7SK. Same as Fig. 6a, but with more details, including the two nucleotides in contact and the numbers of reads in parentheses. Thickness of the black lines are proportionate to the square root of numbers of reads supporting each contact. An arc format is shown on the right.

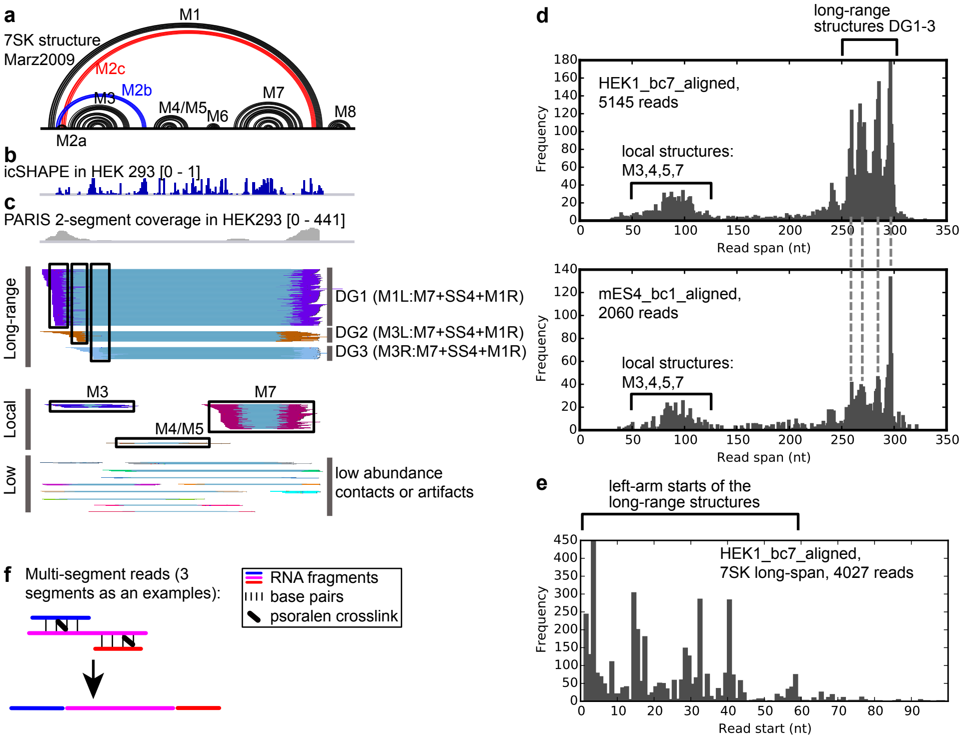


**Supplementary Figure 16. PARIS analysis 7SK RNA secondary structure. a,** The Marz 2009 secondary structure model for the 7SK RNA. Blue arcs: M2b. Red arcs: M2c. **b**, icSHAPE reactivity score from Lu et al. 2016. **c**, PARIS coverage and single-gap DGs clustered by CRSSANT, divided into the long-range, local and low-abundance groups. The 3 long-range DGs represent contacts between the 5' end and the 3' end. DG1 confirms the existence of M1, and the high reads coverage suggest that it is highly abundant, if not the only conformation. DGs 2 and 3 are not consistent with any secondary structures in the Wassarman and Steitz 1991 model, Marz 2009 model, EC model (Weinreb 2016), or the Rfam model, suggesting that they came from previously unknown crosslinkable contacts. The PARIS data here lacked the nucleotide resolution in the SHARC-exo, making it difficult to detect the exact crosslinked nucleotides. M6 and M8 were missed, probably due to lack of the preferred psoralen crosslinking sites (staggered uridines). The low abundance local duplexes were likely from dynamic intermediates of 7SK folding or technical artifacts of proximity ligation. **d**, Analysis of the span of all PARIS gapped reads in human and mouse cells (HEK293 and mouse ES (mES) cells, Lu et al. 2016, PMID: 27180905). There are two types of reads that correspond to local (M3,4,5 and 7) and long-range structrues. In particular, the long-range structures are further sub-divided into 4 groups, roughly corresponding to the 3 DGs. The DGs are defined by overlap on the two arms, and therefore do not correspond exactly with the groups shown here based on the read span. **e,** Plotting the start position of the long-range reads as a histogram, showing several peaks that roughly correspond to the DGs 1-3. These reads start at several distinct locations between 0-60, but also end at the same region, M7+SS4+M1R (see panel c). **f**, Diagram of multi-segment reads. Stronger crosslinking produces complexes with more than 2 RNA fragments, which can be ligated together and sequenced. Such reads indicate that these fragments are in proximity in the same molecule and the same conformation. These multi-segment reads and the 2-segment long-range structures (DG1-3) support interhelical packing of the 7SK RNA.

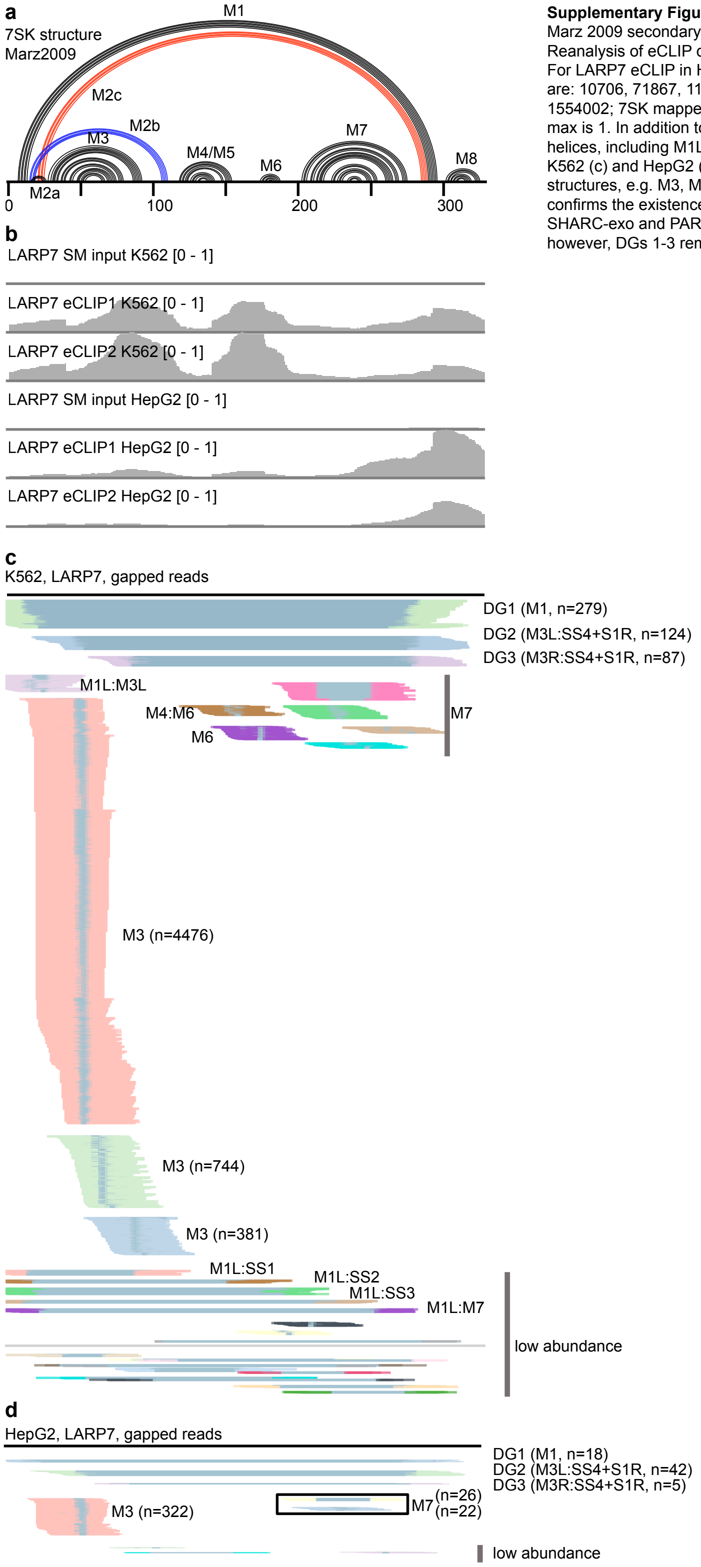


**Supplementary Figure 17. LARP7 eCLIP reveals long-range contacts in the 7SK RNA. a,** The Marz 2009 secondary structure model for the 7SK RNA. Blue arcs: M2b. Red arcs: M2c. **b**, Reanalysis of eCLIP data from K562 and HepG2 cells (Van Nostrand et al. 2016, PMID: 27018577). For LARP7 eCLIP in HepG2, total mapped reads are 5674934, 1381014, 446830; 7SK mapped reads are: 10706, 71867, 11047. For LARP7 eCLIP in K562, total mapped reads are 7302572, 1790120, 1554002; 7SK mapped reads are: 50306, 517215, 673789. The samples were normalized so that the max is 1. In addition to the primary binding site on the 3' end M8 region, LARP7 also binds the 5' end helices, including M1L, M2, M3, and between M4/M5/M6. **c-d**, Gapped reads from LARP7 eCLIP in K562 (c) and HepG2 (d) cells were clustered into DGs using CRSSANT. In addition to capturing local structures, e.g. M3, M6, M7, these gapped reads revealed long-range structures DG1-3. DG1 again confirms the existence of M1 helix, while DGs 2-3 are consistent with newly identified contacts by SHARC-exo and PARIS. The total numbers of reads for 7SK were lower for the HepG2 cells, however, DGs 1-3 remain identifiable.

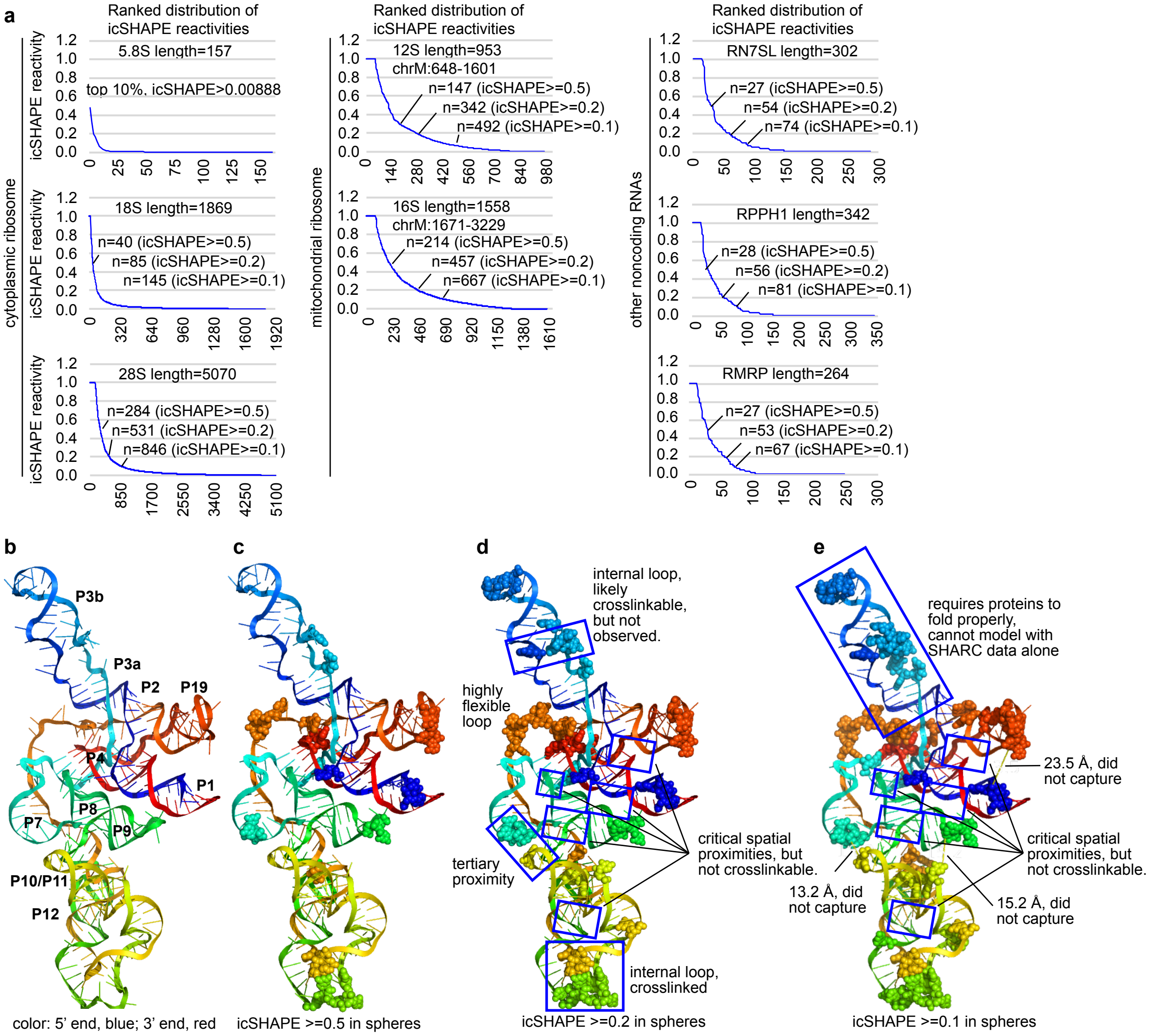


**Supplementary Figure 18. Analysis of RNA icSHAPE reactivity and limitations of SHARC-exo. a,** icSHAPE reactivity for abundant noncoding RNAs was extracted from our recently published sequencing data (Lu et al. 2016, PMID: 27180905)**.** Numbers of reactive nucleotides at different cutoff levels are labeled. The 18S and 5.8S rRNAs are far less reactive compared to other RNAs. The lower reactivity of 18S compared to 28S is probably due to the lower fraction of expansion segments. Typically <10% nucleotides have >0.5 SHAPE reactivity in each RNA. The higher reactivity of the mitochondrial ribosome is likely due to its smaller size and more primitive form. which allows SHAPE reagent to access. These distributions both confirmed the applicability of SHARC-exo to various RNAs, and also showed the limitations. **b-e,** Mapping of icSHAPE reactive nucleotides onto the RNase P RNA (RPPH1) cryo-EM structure model (PDB: 6AHR). RNA was colored blue to red from the 5' end to the 3' end. Even at a very low threshold, many of the critical regions remain non-crosslinkable, due to secondary structure or protein constraints (panel e).

Supplementary Table 1. SHARC-exo library statistics. Unique fraction: reads_unique/reads_raw. Reads_min20: reads with minimal length of 20nts. Total_min20: summing the two libraries for each condition. Min20 fraction: reads_min20/reads_unique
Gapped fraction: (gap1+gapm+chimeric+overlapping_chimeric)/input. Bad alignments represent a small fraction of all alignments, and therefore they are ignored.

| | | Preprocessing | | | | | | Mapping | | | | | | | | |
|---|---|---|---|---|---|---|---|---|---|---|---|---|---|---|---|---|
| samples | barcodes | reads_raw | reads_unique | unique fraction | reads_min20 | min20 fraction | total_min20 | total | input | nongapped | gap1 | gapm | chimeric | overlapping chimeric | bad alignments | gapped fraction |
| HeLa5mMDPI_exo0h | R701 | 8.995.612 | 8.023.454 | 89,19% | 5.564.497 | 69,35% | 11.616.794 | 11.334.506 | 10.807.624 | 9.970.086 | 620.240 | 8.006 | 175.680 | 13.473 | 20.139 | 7,56% |
| HeLa5mMDPI_exo0h | R701 | 9809777 | 8703404 | 88,72% | 6.052.297 | 69,54% | | | | | | | | | | |
| HeLa5mMDPI_exo2h | R723 | 2.132.552 | 1.922.211 | 90,14% | 739.272 | 38,46% | 1.537.923 | 1.327.685 | 1.278.072 | 1.222.774 | 31.552 | 299 | 21.247 | 178 | 2.022 | 4,17% |
| HeLa5mMDPI_exo2h | R723 | 2313864 | 2073503 | 89,61% | 798.651 | 38,52% | | | | | | | | | | |
| HeLa5mMDPI_exo12h | R702 | 14.606.757 | 12.658.542 | 86,66% | 8.160.003 | 64,46% | 17.028.746 | 16.066.509 | 15.464.518 | 14.517.522 | 728.442 | 8.453 | 186.761 | 7.030 | 16.310 | 6,02% |
| HeLa5mMDPI_exo12h | R702 | 15913543 | 13724191 | 86,24% | 8.868.743 | 64,62% | | | | | | | | | | |
| HeLa12.5mMDPI_exo0h | R703 | 6.385.176 | 5.838.913 | 91,44% | 4.438.825 | 76,02% | 9.290.218 | 9.363.603,00 | 8.903.281 | 8.145.260 | 610.889 | 7.994 | 110.184 | 15.334 | 13.620 | 8,36% |
| HeLa12.5mMDPI_exo0h | R703 | 6992022 | 6367057 | 91,06% | 4.851.393 | 76,20% | | | | | | | | | | |
| HeLa12.5mMDPI_exo2h | R708 | 7.135.624 | 6.690.540 | 93,76% | 4.583.161 | 68,50% | 9.569.696 | 8.518.901 | 8.024.575 | 7.193.377 | 630.395 | 14.432 | 168.430 | 5.584 | 12.357 | 10,20% |
| HeLa12.5mMDPI_exo2h | R708 | 7779168 | 7270955 | 93,47% | 4.986.535 | 68,58% | | | | | | | | | | |
| HeLa12.5mMDPI_exo12h | R707 | 11.974.629 | 10.483.576 | 87,55% | 5.523.174 | 52,68% | 11.513.789 | 10.699.524 | 10.094.922 | 9.027.326 | 861.318 | 16.856 | 160.613 | 12.517 | 16.292 | 10,41% |
| HeLa12.5mMDPI_exo12h | R707 | 13039931 | 11351734 | 87,05% | 5.990.615 | 52,77% | | | | | | | | | | |
| HeLa12.5mMDPI_exo24h | R704 | 10.070.228 | 7.198.662 | 71,48% | 814.144 | 11,31% | 1.658.879 | 1.145.588 | 1.100.338 | 1.062.448 | 23.236 | 255 | 12.661 | 150 | 1.588 | 3,30% |
| HeLa12.5mMDPI_exo24h | R704 | 10912598 | 7659816 | 70,19% | 844.735 | 11,03% | | | | | | | | | | |
| HeLa25mMDPI_exo2h | R709 | 5.379.101 | 5.247.760 | 97,56% | 4.282.122 | 81,60% | 8.953.498 | 9.045.708 | 8.332.858 | 7.103.605 | 993.214 | 16.793 | 181.188 | 16.009 | 22.049 | 14,49% |
| HeLa25mMDPI_exo2h | R709 | 5875549 | 5723068 | 97,40% | 4.671.376 | 81,62% | | | | | | | | | | |
| HeLa25mMDPI_exo24h | R717 | 5.102.976 | 4.714.432 | 92,39% | 2.951.322 | 62,60% | 6.158.342 | 5.777.797 | 5.468.182 | 4.994.332 | 354.226 | 4.906 | 94.987 | 7.082 | 12.649 | 8,43% |
| HeLa25mMDPI_exo24h | R717 | 5565513 | 5121082 | 92,01% | 3.207.020 | 62,62% | | | | | | | | | | |
